# RNA polymerases as moving barriers to condensin loop extrusion

**DOI:** 10.1101/604280

**Authors:** Hugo B. Brandão, Xindan Wang, Payel Paul, Aafke A. van den Berg, David Z. Rudner, Leonid A. Mirny

## Abstract

To separate replicated sister chromatids during mitosis, eukaryotes and prokaryotes have structural maintenance of chromosome (SMC) condensin complexes that were recently shown to organize chromosomes by a process known as DNA loop extrusion. In rapidly dividing bacterial cells, the process of separating sister chromatids occurs concomitantly with ongoing transcription. How transcription interferes with the condensin loop extrusion process is largely unexplored, but recent experiments show that sites of high transcription may directionally affect condensin loop extrusion. We quantitatively investigate different mechanisms of interaction between condensin and elongating RNA polymerases (RNAP) and find that RNAPs are likely steric barriers that can push and interact with condensins. Supported by new Hi-C and ChIP-seq data for cells after transcription inhibition and RNAP degradation, we argue that translocating condensins must bypass transcribing RNAPs within ~2 seconds of an encounter at rRNA genes and within ~10 seconds at protein coding genes. Thus, while individual RNAPs have little effect on the progress of loop extrusion, long, highly transcribed operons can significantly impede the extrusion process. Our data and quantitative models further suggest that bacterial condensin loop extrusion occurs by two independent, uncoupled motor activities; the motors translocate on DNA in opposing directions and function together to enlarge chromosomal loops, each independently bypassing steric barriers in their path. Our study provides a quantitative link between transcription and 3D genome organization and proposes a mechanism of interactions between SMC complexes and elongating transcription machinery relevant from bacteria to higher eukaryotes.

## Introduction

The structural maintenance of chromosome (SMC) complexes are an evolutionarily conserved family of protein complexes including condensin, cohesin, SMCHD1, Smc5/6 and others, present in most organisms from eubacteria to humans (1). These proteins are involved in processes as diverse as DNA damage repair, sister chromatid cohesion, and organization of mitotic and interphase chromosomes. SMC complexes are characterized by a three-part ring, composed of a dimer of SMC subunits each with an ATPase domain and a long coiled-coil domain, a kleisin linker which closes the ring, and accessory proteins which bind to the linker to perform specific functions, depending on the organism (1).

Recent *in vivo* studies have provided evidence that SMC complexes have a motor activity, allowing them to translocate processively on a chromatin fiber and perform active chromatin reorganization by loop extrusion. In the proposed loop extrusion mechanism (2–8), SMC complexes (or oligomers of SMC complexes) bind to DNA at a single site, bridge two flanking DNA segments forming a loop, and then progressively enlarge the loop by translocating away from the loading site. Thus, loop extrusion is thought to result from the activity of two connected motors translocating in opposite directions that expand a DNA loop.

Single molecule studies provide support for this or a similar mechanism by demonstrating that budding yeast condensin SMCs are mechano-chemical motors that can translocate along DNA (9), extrude DNA loops (10), and can actively compact DNA (11–13). While the molecular details of this process are yet to be fully understood (14, 15), loop extrusion appears to be a mechanism that can explain a wealth of chromosomal phenomena in eukaryotes and bacteria.

In eukaryotes, during mitosis, loop extrusion by condensin can explain the compaction and resolution of sister chromatids and may underlie the formation of arrays of loops and nested loops central to mitotic chromosomes. Evidence also suggests that loop extrusion by cohesin SMCs underlies the formation of chromosomal domains during interphase (16–20).

In bacteria, condensin SMC complexes help resolve newly replicated origins and appears to do so by juxtaposing the left and right chromosome arms of the newly replicated sister chromosomes. Amazingly, DNA juxtaposition extends from origin to terminus generating a single 4 Mb “loop” (7, 21, 22). Studies using chromosome conformation capture (Hi-C), and chromatin immunoprecipitation combined with deep sequencing (ChIP-seq), have shown that condensins are preferentially loaded onto chromosomes via ParB proteins bound at *parS* sites located primarily adjacent to the origin of replication (23–25). Once loaded, condensins progress away from the *parS* sites (8, 26) along both chromosomal arms, thus juxtaposing them (7, 21), resulting in the characteristic “X” shaped pattern on the Hi-C maps of many bacteria (27). *In vivo* experiments recently showed that translocation by condensin is an active process (8, 28): condensin complexes travel processively and bidirectionally away from the *parS* loading site, in a manner that appears to be ATP-dependent, at speeds exceeding 800 bp/s (29). This active juxtaposition of chromosome arms by the bacterial condensin SMC condensin complex suggests a mechanism of loop extrusion in bacteria (7, 8, 28, 30) nearly identical to the proposed loop extrusion process in eukaryotes.

Beyond their function in directly shaping spatial chromosome structure, SMCs can potentially also respond to various signals allowing the reorganization of chromosomes in response. As an example, recent studies provide strong evidence that transcription can affect genome structure and SMC action (31–36). It remains unknown, however, how an SMC loop extruder interacts with the transcription machinery, and how these nanometer scale interactions affect global chromosome structure.

Here, we study the effect of transcription on chromosome structure by developing models of condensin dynamics and validating them using experimental data. Central to these models is the hypothesis that the speed of condensin translocation is affected by transcription depending on the relative orientation of genes and the direction of extrusion. We propose that once a condensin encounters an actively transcribed gene, it slows down due to interactions with the transcription machinery, with the slowing down being greater if condensin and RNA polymerase meet in a head-to-head versus head-to-tail interaction.

Our models predict condensin juxtaposition trajectories that are in excellent quantitative agreement with Hi-C data for wild-type and engineered bacterial strains where the condensin loading site has been moved to different genome positions. Our analysis further supports the idea that loop extrusion by bacterial condensins is mediated by at least two independently acting and uncoupled motor activities. To understand the molecular mechanisms that underlie the directional effect of slow transcription (~40-80 bp/s) on the much faster condensin translocation (~800 bp/s), we develop a mechanistic “moving barriers” model for interactions of SMC complexes with transcription machinery. The analytical solution of the stochastic “moving barriers” model allows us to integrate diverse experimental data to predict chromosome structure arising through the interplay of loop extrusion and transcription. We find strong evidence that SMC molecules can bypass elongating RNA polymerases impeding their translocation within 2 seconds of an encounter at rRNA operons, and within 10 seconds at protein coding operons. This finding has important implications for understanding the mode of DNA translocation by condensins, and their ability to overcome steric barriers. We also investigated changes in DNA juxtaposition following transcription inhibition and acute RNAP degradation. Our analysis revealed that both transcription-dependent and transcription-independent effects impact condensins’ genome-wide chromosome juxtaposition activity. Our quantitative models of transcription-condensin interactions tested on bacterial data have widespread implications for chromosome organization, and can provide a framework to study the effect of transcription on chromosome organization in higher organisms

## Results

### Predicting condensin’s spatiotemporal trajectory from gene directions and positions

In the Hi-C map of wild-type *B. subtilis*, contacts from DNA segments close in the linear genome sequence, as in all organisms, give rise to the primary interaction diagonal, which extends from the bottom left to top right of the map (Fig. 1A). A secondary diagonal, which runs perpendicularly to the primary, typical of many bacterial Hi-C interaction maps (7, 22, 27), indicates a symmetric juxtaposition of two chromosome arms about the *parS* sites located next to the origin of replication (*ori*) (Fig. 1A, inset). However, engineered strains of *B. subtilis*, in which all the endogenous *parS* sites are deleted, and a single site is inserted at other positions, reveal different shapes of the Hi-C secondary diagonal. In such strains, the wild-type secondary diagonal is missing and is replaced with a diagonal emanating from the new *parS* location (7, 8). Without exception, these new interaction signatures are tilted or curved away from the *ori* (Fig. 1B). These curved diagonals represent an asymmetry in interactions between DNA flanking these displaced *parS* sites. In all cases, larger tracks of terminus-proximal DNA interact with shorter tracks of origin-proximal DNA. In the loop extrusion model, chromosome juxtaposition occurs by two motor activities of condensin translocating away from the *parS* site (Fig. 1C); at the molecular level, the secondary diagonal visible by Hi-C arises from individual trajectories of condensin’s loop extrusion motors, averaged over a population of cells. Thus, in the context of this model, a curved diagonal suggests that at certain loci one of the loop-extruding motors translocates more slowly that the other one (Fig. 1D).

**Figure 1:**
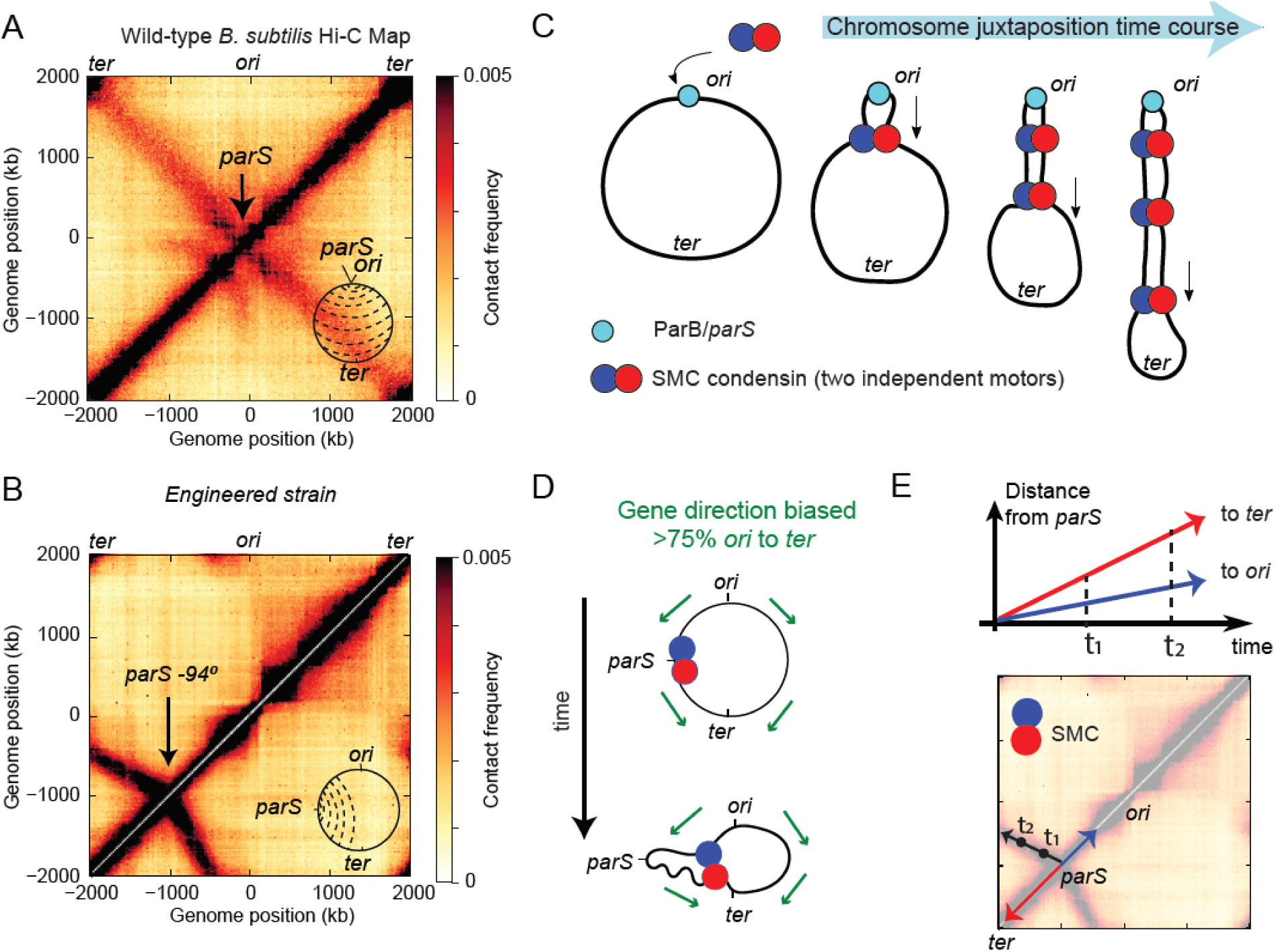
The posited role of transcription in shaping asymmetric SMC translocation rates. **(A)** Hi-C map of a wild type *B. subtilis* PY79. The inset depicts the juxtaposition of chromosome arms (dashed lines) centered on the *ori* to *ter* axis. (**B**) Hi-C map of a *B. subtilis* strain (see Supporting Information for strain tables) with a single *parS* site at −94° which has a secondary diagonal that points biasedly away from the *ori*. **(C)** Loop extrusion model schematic depicting the active juxtaposition of chromosome arms performed by the *B. subtilis* condensin loop extruding complex. (**D**) Gene directions point biasedly towards the *ter* (green arrows); for SMC motors translocating towards the *ori* (blue), there will be increased frequencies of head-on collisions with transcripts as compared to SMCs translocating towards the *ter* (red). (**E**) Interpretation of Hi-C: condensin translocates bidirectionally from the *parS* site juxtaposing flanking DNA; motion towards the *ter* is faster than towards the *ori* resulting in the asymmetric secondary diagonal.

Recalling that over 75% of genes in *B. subtilis* are co-oriented with replication (37), we posited that transcription could account for the tilt of the secondary diagonal by slowing down condensin translocation (Fig. 1D). Since condensin translocation towards the *ori* will be more frequently opposing transcription, the increased numbers of “head-to-head” encounters of condensin with RNAP potentially leads to a slower overall translocation rate for *ori*-oriented condensins. This results in a gene-direction based effect on condensin speed (Fig. 1E). This hypothesis is supported by recent experimental evidence in *C. crescentus* and *B. subtilis* where the relative orientations of genes to condensin loading sites have been altered (8, 28).

To test whether the interplay between condensin translocation and transcription can shape chromosome structure, we developed a model where condensin trajectories can be predicted based solely on gene locations and orientations (Fig. 2A). In this model, condensins form a loop extrusion complex that has two motors, each translocating independently and deterministically along the DNA with the maximum speed, v_max_; the complex begins at a single point (the *parS* site) and each motor progresses in opposing directions with the following rules: When a motor encounters a gene its instantaneous speed v is changed such that

**Figure 2:**
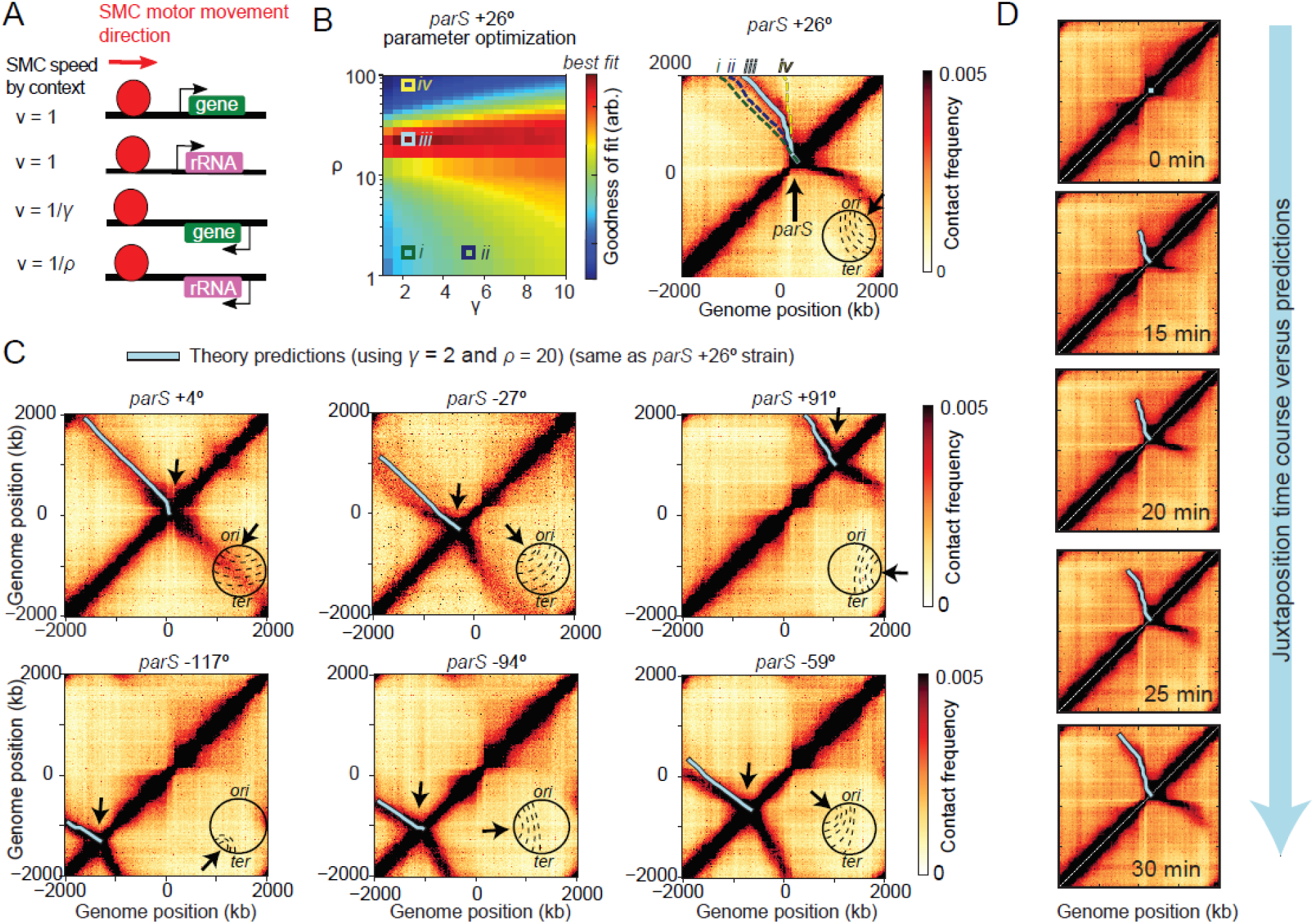
Model of SMC translocation based on gene position and orientation. (**A**) Rules for the model of condensin translocation rates based on gene orientation. (**B**) A parameter sweep of the model (left panel) shows the agreement with Hi-C data as a function of *γ* and *ρ* parameters; illustrative example trajectories (i-iv) are superimposed on the Hi-C map to show how juxtaposition traces change as a function of these parameters (right panel). (**C**) Examples of using the *parS+26*° strain-specific optimum parameters (*γ* = 2 and *ρ* = 20) (see Fig. S4 for globally optimum trajectories) to predict the juxtaposition trajectories of other engineered *B. subtilis* strains. (**D**) The model of condensin translocation using only gene positions and orientations captures the spatiotemporal behavior of Hi-C secondary diagonal formation during a time-course Hi-C experiment where condensin loading was induced at t = 0 min.

v = *v*_*max*_ if the condensin motor moves in the direction of gene transcription,

v = *v*_*max*_/*γ* if the condensin motor moves in the direction opposing gene transcription,

v = *v*_*max*_/*ρ* if the condensin motor moves in the direction opposing the rRNA loci, which are the most highly transcribed genes,

(v = *v*_*max*_ in the absence of annotation).

The value γ is interpreted as the fold-increase in time required for a condensin motor to traverse a gene against the direction of transcription. ρ is similarly interpreted but reserved for the highly transcribed rRNA operons which are found at 7 distinct loci in *B. subtilis PY79*. Below, we generalize this model to incorporate locus-specific rates of transcription. Crucially, we assumed that speeds of condensin motors in the same extrusion complex are independent of each other, i.e. if one motor is slowed down, the other continues unaffected at its own speed; we revisit this assumption later. As the two motors move away from the *parS* site, they bring together flanking DNA, generating the Hi-C secondary diagonal (Figs. 1C, 1D). By computing the displacement from *parS* versus time of each motor, base pair by base pair using gene position and direction data (see Supporting Information), it is possible to trace the expected extrusion complex trajectory parametrized by time on the Hi-C map (Fig. 1E); the trajectory depends on the values of γ and ρ as shown (Fig. 2B, right).

We hypothesized that if transcriptional interference with loop extrusion is a universal phenomenon which depends largely on gene orientation, then a single set of the parameters γ, ρ might predict the shapes of secondary diagonals in different engineered bacterial strains with ectopic condensin loading sites. By sweeping over parameter values and comparing predicted extrusion traces to Hi-C experiments, we can determine the best values of γ and ρ (Fig. S1A). For example, we can use the best values of γ and ρ found for one strain, i.e. *parS*+26^0^, (Fig. 2B) to adequately predict the condensin trajectories in 9 other strains (Fig. 2C; Fig. S1B). Moreover, and importantly, we find that the optimal solutions for γ and ρ across strains have similar values (Fig. S1A). Combining parameter-fit values of the different strains (Fig. S2A), we find that a single set of parameters (γ =3.5, ρ = 20) provides the overall best predictive power for the secondary diagonals and extrusion traces (Fig. S2B) and resembles the strain-specific optimum trajectories (Fig. S1A).

To validate the temporal aspect of our above model of condensin loop extrusion, we tested whether it agreed with time-course Hi-C of chromosome juxtaposition (8). In the time-course experiments, loading of condensin at the *parS* site was induced at time t = 0 minutes, and the progression of the juxtaposition front was monitored by Hi-C over 5-minute time intervals. Using the average juxtaposition rate of 800 bp/s measured previously (8), we calibrated the relative condensin speeds into absolute speeds. In our model, knowing the average condensin speed, v_avg_, we infer the maximum speed, v_max_, using

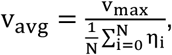

where *N* is the number of base pairs of a genome arm, and *i* is the relative translocation time to move across a locus (where η_i_ = γ, ρ if the base-pair, η_i_, belongs to a gene or rRNA locus that is oriented opposite to condensin’s translocation, and η_i_=1 otherwise). Plotting the model predictions made for each arm using the two globally optimal parameter values *γ* = 3.5, *ρ* = 20, known experimental values for *v*_*avg*_ = 800 bp/s, *N* = 2×10^6^ (base-pairs), we find excellent quantitative agreement with the time-course of experimental Hi-C (Fig. 2D; Fig. S3), and infer that the maximum speed of condensin is *v*_*max*_ ≈1500±200 bp/s.

Thus, this minimal model which uses only gene positions and orientations agrees well with experimental data; it captures the major aspects of the Hi-C secondary diagonals and suggests that transcription orientation is a key factor in controlling the speed of extrusion. However, the model does not establish a direct link between condensin translocation and the process of transcription. Accordingly, it remains a possibility that other DNA motifs or processes, correlated with gene orientations, influence condensin speeds (38). This calls for more direct experimental tests for the role of genes and transcription. Nonetheless, we can conclude from this analysis that, either via transcription or other mechanisms, origin to terminus sequence biases strongly influence condensin translocation and alters chromosome organization in a predictable and universal way.

### Bidirectional condensin translocation is performed by two independent motor activities

The model above and previous analyses (8) suggests that *B. subtilis* condensin complexes bi-directionally translocate along chromosomal arms enlarging chromosomal loops by two independent motor activities. We sought to rigorously test whether this assumption of independence is necessary, and the degree to which it holds true. We modified the model such that the instantaneous waiting times for each locus (*η*_*i*_ = *γ*, *ρ*) were partially correlated between the two motors, thereby breaking the assumption of independent translocation for each motor. Correlation was quantified by the mixing parameter *f* (*f* = 0 for independent motion, *f* = 1 for fully correlated motion) (See Supporting Information for details).

As done previously, we swept the parameters *γ*, *ρ*, for various fixed values of *f* in the mixing model (ranging from *f=*0 to *f=*1) to obtain goodness of fit values (Fig. S4A). Intriguingly, the model with *f=0* (independent translocation) had the highest overall goodness of fit value (Fig. S4B) and exhibited visually better predictions (Fig. S5). Thus, our model strongly suggests that the translocation process occurs via two independent motor activities, which do not sense impediments to their counterpart in the other translocation direction. This finding argues that dimerization (or oligomerization) of condensin may be required to obtain two distinct motor activities, consistent with previous experimental evidence and theoretical models (8, 14, 39, 40), or that a single one-sided condensin dynamically switches directions of translocation leading to apparent “two-sided” extrusion, with effectively independent motor activities (41).

### Transcription slows down the condensin translocation rate at highly transcribed genes

To directly test the effect of transcription on condensin translocation, we studied how transcription inhibition affects chromosome structure as assayed by Hi-C. In a strain with a *parS* site close to a large cluster of rRNA operons (*parS+26*°), adding rifampicin to exponentially growing cells, which inhibits the transition from transcription initiation to elongation (42), results in a partial straightening out of the secondary diagonal as shown previously (8, 28) (Fig. 3A). We reasoned that if transcription elongation shapes the overall tilt in the secondary diagonal, our quantitative model (Fig. 2A) should reveal a decrease in both the *γ* and *ρ* values following transcription inhibition.

**Figure 3:**
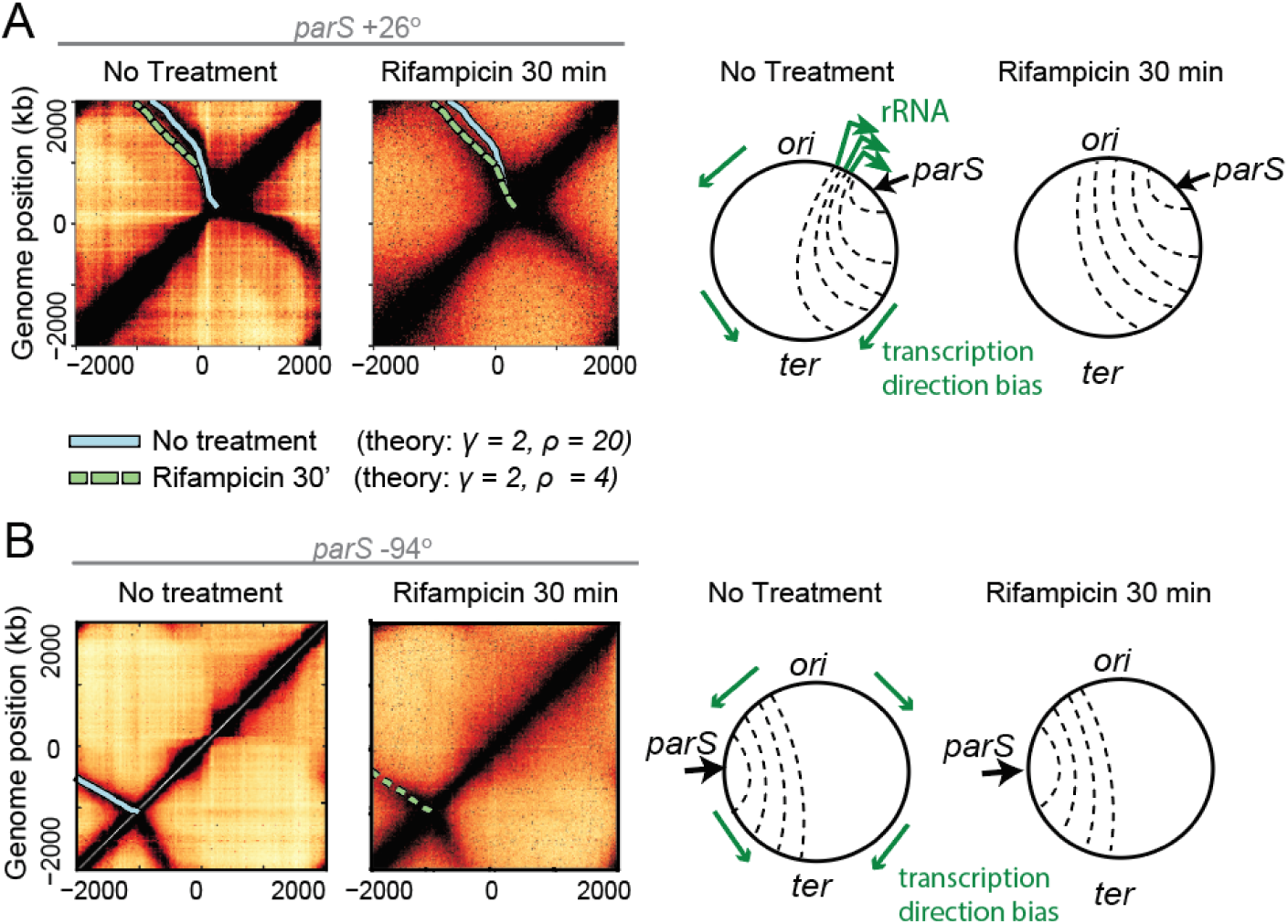
Transcription-dependent and independent features of SMC-mediated chromosome juxtaposition. (**A**) Hi-C data from previous experiments (Wang et al., 2017) showing the effect of transcription inhibition by rifampicin for 30 minutes on chromosome arm juxtaposition; superimposed SMC translocation model trajectories (solid and dashed lines) suggest a transcription-elongation independent effect on asymmetric SMC (i.e. the factor γ is unchanged before and after treatment; see Fig. S6A); schematic representations of the changes to chromosome juxtaposition are shown on the right. (**B**) Hi-C before and after rifampicin treatment experiments for a strain (*parS+94*°) with condensin loading far from the highly transcribed rRNA clusters of the other strain (*parS*+26^0^); the unchanging secondary diagonal angle confirms a transcription independent effect.

We fit the previously published data on transcription inhibition to obtain the best fit values for *γ*, *ρ* before and after treatment (Fig. S6A). To our surprise, we find that *γ* = 2, *ρ* = 4.5 best describe the data after treatment and *γ* = 2, *ρ* = 20 before treatment (Fig. 3A; Fig. S6A). This suggests that the overall tilt (“baseline asymmetry”) in the Hi-C secondary diagonal away from rRNA loci, captured by our parameter *γ*, is largely independent of transcription elongation (i.e. γ = 2 before and after treatment). Conversely, *ρ*, which quantifies the slowdown of condensin translocation going head-to-head with transcription at rRNA loci, largely depends on elongation.

To independently investigate this observation, we performed additional transcription inhibition experiments using a strain with a *parS* site at the −94° position. In this strain, the *parS* site is far from the highly transcribed rRNA loci that affect condensin movement in the former strain (*parS*+26°). Consistent with the effect suggested by our model, we observed virtually no changes to the angle of the Hi-C secondary diagonal in the newly tested strain after 10 min (Fig. S6B) or 30 min (Fig. 3B) of rifampicin treatment. This suggests that protein-coding genes (non-rRNA) have little effect on the on the speed of condensin translocation. These observations and the partial but not complete straightening out of the secondary diagonal in the *parS*+26° experiments led us to consider two possible models: either non-transcribing RNAPs (e.g. trapped at transcription start sites by rifampicin) are directional barriers to condensin translocation, akin to CTCFs as directional barriers to cohesin in eukaryotes (4), or an RNAP independent mechanism generates the “baseline asymmetry” of chromosome juxtaposition at loci outside of rRNA operons.

To differentiate between these the two models, we analyzed changes to chromosome structure following the degradation of RNAP. We reasoned that degrading RNAP will have no effect on the secondary diagonal tilt if the mechanism that generates the asymmetric juxtaposition is RNAP independent. We generated a strain in which the sole copy of the β’ (beta prime) subunit of RNAP was fused to YFP and an SsrA tag (β’-YFP-SsrA) and could be conditionally targeted for degradation (43)(Fig. S7). We induced degradation of the β’ fusion in a strain with a *parS* site at the −59° position and monitored the levels of protein over 90 minutes; β’ levels dropped to 5% of their initial value, as assayed via quantitative immunoblotting (Fig. S7C) and imaging of β’-YFP-SsrA fluorescence in single cells (Fig. S7B). Strikingly, a time-course Hi-C after induction of RNAP degradation revealed only a minor change in the tilt of the secondary diagonal (Fig. S7A). As with rifampicin treatment, the most significant changes to the Hi-C maps manifested as a “blurring” of Hi-C features along the main diagonal and the disappearance of high intensity spots along the secondary diagonal (Fig. S8A). After RNAP degradation for 90 minutes, contact probability at short distances (<200 kb) decayed more quickly as compared to normal growth conditions (Fig. S9A), and was indistinguishable from contact probabilities of cells treated for 30 minutes with rifampicin (Fig. S9B). Thus, RNAP and transcription are necessary for creating the texture in the Hi-C maps, which results in increased DNA-DNA contacts within ~200 kb of separation, however, it does not strongly affect the overall tilt. Altogether, the degradation of RNAP experiments rule out the role of paused RNAPs in establishing the “baseline asymmetry” of chromosome juxtaposition; this further indicates that the condensin translocation slowdown towards the *ori* (at non rRNA loci) is largely independent of RNAP.

In hindsight, we understand that our phenomenological model (Fig. 2A) worked so well because in *B. subtilis* gene density is high and homogeneous (Fig. S8B), and there are relatively few highly transcribed genes (44); the average RNAP density is of 0.1 RNAP/kb for most genes (45), in contrast with ~10 RNAP/kb for rRNA genes (45) in our growth conditions (see Supporting information). Thus, the parameter *ɣ* reflects a systematic *ori* to *ter* bias in the condensin translocation speed (i.e. the “baseline asymmetry”) which correlates with gene direction but is largely RNAP independent. We posit that such a bias may come from the process of DNA replication and will be a topic of future study. In contrast to *ɣ*, however, the parameter *ρ*, which reflects condensin slow down at rRNA loci, does depend strongly on transcription elongation. We thus choose to focus on understanding how the parameter *ρ* emerges from transcription elongation. Accordingly, we test mechanistic models to help explain how highly transcribed genes become directional barriers to condensin translocation.

### The moving barriers mechanism of condensin-transcription interactions

Plausible mechanisms of SMC and RNAP interaction must solve the following puzzle: how can condensin’s effective speed of translocation (measured at >800 bp/s via Hi-C and ChIP-seq (8)) be so strongly attenuated (>20-fold) by RNAP transcription (which moves at 40-90 bp/s (46)) depending only on the relative orientation of the two processes?

Previous studies have suggested that a passive SMC ring can be pushed by transcribing RNAP (34, 47). This idea emerged from the observation that cohesin SMCs are enriched at sites of convergent transcription in yeast (47, 48) and in mammalian cells (35); this is further supported by other experiments demonstrating localization of cohesin mediated by transcription in cells (32) and *in vitro* (49). The central assumption of the models is that transcription machinery forms an impermeable moving barrier to the SMC ring, pushing it along the direction of transcription.

By combining the moving barrier idea with active translocation by condensin, we can intuitively explain the directional effect of RNAP elongation on SMC translocation at rRNA loci, and the emergence of the direction parameter, ρ. In the case of head-to-tail interactions between condensin and RNAP (Fig. 4A, left), a condensin motor translocates at a high rate (e.g. ~1500 bp/s) until it encounters a transcribing RNAP moving in the same direction at a much lower speed (e.g. 80 bp/s as measured in *E. coli* (46)). Since RNAP is assumed to be an impermeable barrier (we generalize this later to allow for partial permeability), when condensin encounters an RNAP, it slows down its translocation rate to match the RNAP until the end of the operon. Dissociation of RNAP at the end of an operon allows the condensin to continue translocating at its original high speed. In contrast, in the case of head-to-head interactions (Fig. 4A, right), when a translocating condensin encounters a transcribing RNAP, condensin is stalled and pushed back to the transcription termination site (condensin has a very low stall force measured *in vitro* (10) compared to RNAP (50)). Once RNAP dissociates at the transcription termination site, condensin is left to attempt crossing the operon again; the condensin will only successfully cross the operon if no RNAPs are encountered during its run through the operon. In this “moving barriers” model, the relative directional slowing down of condensin arises from the fact that multiple attempts may be required for the condensin to successfully cross an operon in a head-to-head orientation.

**Figure 4:**
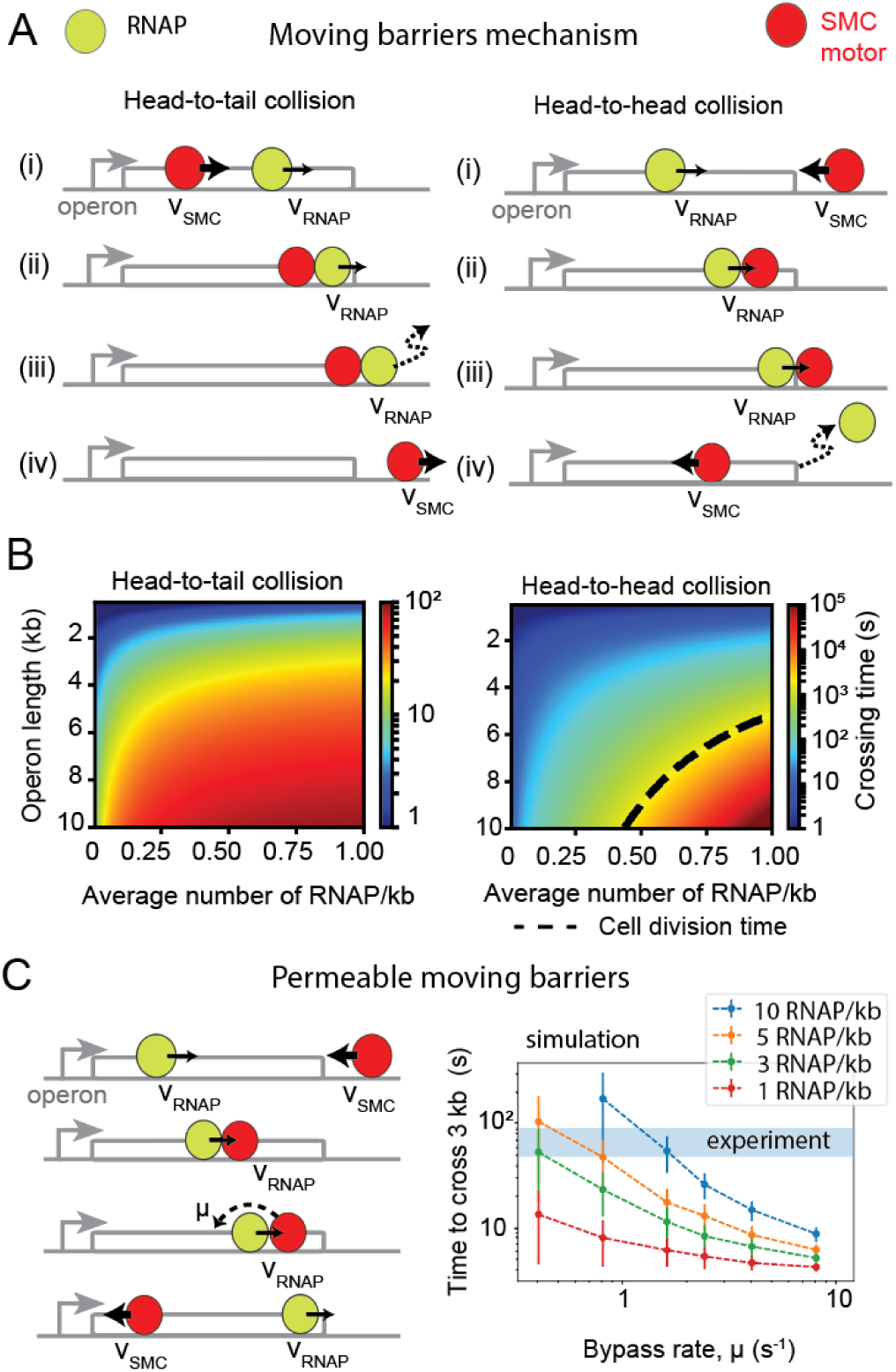
Mechanistic models of extrusion-transcription interference. (**A**) The “moving barriers” model of condensin transcription interactions: transcription complexes are posited as impermeable barriers to condensin movement which results in different translocation dynamics when crossing operons in the co-oriented or convergent fashion. Condensin translocates (i) at its native speed (v_SMC_) until (ii) it reaches a slowly moving RNAP, then (iii) it proceeds in the direction and at the speed of RNAP (v_RNAP_) until (iv) RNAP reaches the end of the operon whereby condensin continues translocation at its original speed and direction. (**B**) Analytically computed average times to cross an operon for each of the head-to-tail (left panel) and head-to-head (right panel) case. as function of operon length and RNAP density. Despite similar “rules”, in the head-to-tail case (left panel) the SMC always reaches the end of the operon, whereas in the head-to-head case (right panel), a successful traversal by SMC may occur only if no RNAP is encountered within a cell division time (dashed line). (**C**) Extension of the “moving barriers” model allowing for condensin to bypass (“hop”) over transcribing RNAP (left, schematic); simulations of the locus-crossing times with varying permeability (“bypass”) rates and RNAP densities (right panel); blue region indicates the experimentally estimated time for condensin to cross a 3 kb rRNA gene locus (with density ~10 RNAP/kb).

The moving barriers concept can be incorporated into a quantitative model, allowing to compute the parameter, *ρ*. To obtain theoretical estimates for the times to cross a locus in the head-to-head versus head-to-tail cases, we solved the moving barriers model analytically (Fig. 4B) (see Supporting Information). The ratios of the calculated head-to-head to baseline operon crossing times (see Supporting Information) give us a theoretical local value of *ρ* as a function of operon lengths, and RNAP density, and produces a strong directional effect with *ρ* ≫ 1, as desired. However, for characteristic rRNA operon lengths (10 kb) and average rRNA locus densities (~10 RNAP/kb), the calculated directionality parameter produces values of *ρ* > 10^4^ (Fig. S9A), far exceeding the sought range of *ρ* ≈ 20-100 (Fig. S1A). These calculations raise the possibility that translocating condensins can somehow bypass (“hop over”) elongating RNAP.

As a consistency check, we investigated whether the moving barriers model would support the observations that transcription inhibition (Fig. 4B) and RNAP degradation (Fig. S7A) resulted in only a minor change to the tilt of the secondary diagonal. Using the moving barriers model, we calculated the relative contribution to our parameter *ɣ* due to transcription elongation at regular operons (i.e. non rRNA). For operons of length ~3 kb, and average RNAP densities of ~0.1 RNAP/kb, the ratio of head-to-head versus head-to-tail crossing times is ~1.3 (see Supporting information). The ~1.3-fold relative slow-down suggests that active transcription can only account for a *ɣ* value up to ~1.3. Since *ɣ* is found to be between 2 and 7 (Fig. S1A), this suggests that transcription elongation can contribute only up to 30% of the observed tilt. If condensins can bypass RNAPs (as we will see below), then the upper limit of 30% will be further reduced. Thus, the model agrees with the apparent lack of change in the secondary diagonal tilt. Interestingly, this calculated fold-increase in crossing times between the head-to-tail and head-to-head encounters agrees well with other recent experimental results (28). In *C. crescentus*, the enrichment of SMCs within genes was up to 1.4-fold larger depending on whether the genes transcribed “against” or “with” the direction of condensin translocation (see Fig. 5 in Ref. (28)). This suggests that the moving barriers concept is not only applicable to *B. subtilis* genes but is a general feature of SMC interactions with transcription in other organisms.

**Figure 5:**
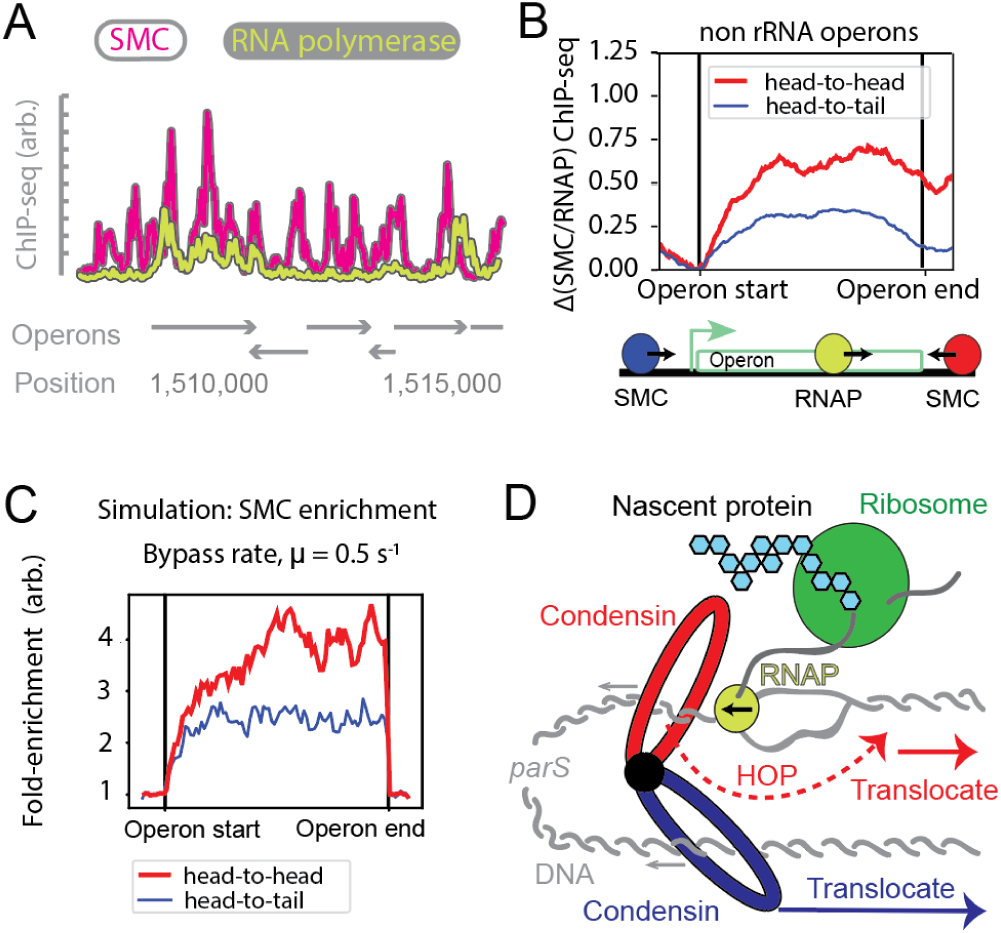
Evidence for the “moving barriers with permeable boundaries” model of condensin-transcription interactions. (**A**) Comparison of SMC ChIP-seq tracks (anti-SMC) with RNAP tracks (anti-GFP, RpoC-GFP) showing RNAP colocalizes with SMC, but SMC peaks may occur without RNAP; operon locations are shown. (**B**) Fold-difference of ChIP-seq signal for SMC tracks normalized by RNAP tracks; average signal is shown for all genes of length up to length 1 kb, separated by the direction of SMC translocation relative to transcription direction (see Supporting information). (**C**) Simulation SMC ChIP-seq for 1 kb gene, demonstrating the differential accumulation of SMC within the gene body for the “moving barriers with permeable boundaries” model; a permeability rate of 0.5 s^−1^ well describes the ~2 fold-change in experimentally observed head-to-tail versus head-to-head SMC accumulation. (**D**) Summary model: condensins (possibly oligomers) translocate away from the *parS* loading site by two independent (blue and red) motor activities; condensin motors can bypass steric barriers (like transcription machinery, or other DNA bound proteins) which are larger than the condensin lumen; while condensin attempts to bypass a steric barrier, it may be “pushed” by other translocating factors like RNAP leading to transcription-dependent SMC translocation rates.

### Translocating condensins can efficiently bypass sites of active transcription

To study the possibility that translocating condensin can bypass transcribing RNAP, we generalized our moving barriers model by introducing a finite permeability to the barrier. In the permeable moving barriers model, condensins that are hindered by a transcription complex can bypass it with a characteristic rate, µ (Fig. 4C, right panel; Fig. S10A), i.e. pausing at each RNAP for on average 1/ µ seconds. The limiting case *μ* → 0 s^−1^ is the impermeable barriers model (Fig. 4A), and the limit *μ* → ∞ s^−1^ is where condensins do not interact with RNAP at all.

We studied the model analytically and performed 1D simulations of the RNAP and condensin translocation with varying permeability (or bypass) rates, µ, and computed the average times for condensin to cross operons of various lengths (1 kb to 10 kb). We searched for permeability rates which would reproduce rRNA head-to-head and head-to-tail locus crossing times measured by Hi-C as well as the parameter *ρ* ≈ 20. Time-course Hi-C data indicate that to cross the clusters of rRNA operons near the *ori* (e.g. *parS+26*° strain) it takes <1 minute for condensins travelling along the direction of transcription (Fig. S3A), and between 8-15 minutes against transcription (Fig. S3B). Assuming RNAP densities of ~10 RNAP/kb, we found from simulations that the permeability rate µ ≈ 0.8-1.6 s^−1^, was most consistent with the experimental data on rRNA locus crossing times, and µ ≈ 0.6-1.7 s^−1^ from the analytical model (Fig. 4C; Fig. S10B; see Supporting Information). Reassuringly, this range of rates also reproduced the value *ρ* ≈ 20 required to reproduce condensin traces in Hi-C data (Fig. S10B). Together, these results suggest that in crossing the rRNA locus, condensins stall for 0.5-2 seconds at each elongating RNAP molecule before bypassing it. Curiously, the rate at which SMCs bypass RNAP molecules (0.5-1.6 s^−1^) is very close to the rate of ATP consumption by SMCs (~0.7±0.1 ATP/condensin/s) (29) suggesting that ~1 ATP cycle (or SMC “step”) is required for condensin to bypass an elongating transcription complex. Interestingly, these estimates suggest that most genes (due to low levels of transcription compared to rRNA genes, and high permeability values) will not significantly slow down condensin translocation; this is consistent with the small effect of transcription inhibition on the “baseline asymmetry” of chromosome juxtaposition (i.e. parameter γ).

We next sought to independently validate the permeability rate estimates measured above. The analytical formulation of the permeable moving barriers model makes several predictions. While most genes will not measurably change the overall condensin trajectories, they will leave signatures in SMC accumulation patterns measurable by ChIP-seq. We make three predictions: First, there will be a positive correlation between RNAP and SMC ChIP-seq signals. Second, the model predicts a non-uniform SMC accumulation pattern within operon bodies: wherever RNAP accumulates within a gene body, our theory predicts that SMCs will also accumulate. Additionally, even for uniform RNAP distributions, simulations of head-to-head encounters suggest that for high RNAP densities and inferred permeability rates (e.g. 10 RNAP/kb and µ = 0.8 s^−1^), there will be a strong accumulation of SMC at transcription termination sites, whereas for lower RNAP densities and similar rates (e.g. 1 RNAP/kb and µ = 0.8 s^−1^) the SMC distribution will be more uniform (Fig. S10C); the pattern and strength of accumulation is a function of the permeability rate, RNAP density, and distance from the transcription start site (Fig. S10C). Third, and most importantly, the model predicts a stronger ChIP-seq enrichment for condensins crossing operons in the head-to-head versus head-to-tail directions (Fig. S10B, S10C).

Thus, to test the permeable moving barriers model we performed ChIP-seq for RNAP and condensin. As expected, we found a strong positive correlation between RNAP and SMC ChIP-seq signals (Fig. S11A) (Pearson correlation coefficient, R=0.51, p<10^−28^). Then, visualizing SMC and RNAP ChIP-seq signals alongside genome annotations, we found that wherever RNAP accumulates, SMC also accumulates, consistent with the analytical theory (Fig. 5A). Interestingly, the reverse was not always true (Fig. 5A); this suggests that RNAPs can be barriers to translocating condensins, but that other DNA bound proteins may also be barriers; consequently, this makes our estimates of the permeability rates lower bounds on the true rates. Lastly, to probe for the gene body and gene direction dependent SMC accumulation, we performed an aggregate analysis of SMC accumulation within operons. We found that SMC accumulates two times more strongly in operons where the transcription direction opposes condensin’s translocation direction (Fig. 5B; Fig. S11B) consistent with our simulations (Fig. 5C) and analytical model (Supporting Information). The simulations suggest a range µ ≈0.1-0.8 s^−1^ are in the best agreement with the SMC ChIP-seq data using estimated numbers of RNAP (~1-3 RNAP/transcription burst) (51) (Fig. 5C; Fig. S10B); the analytical model suggests µ ≈ 0.12-0.36 s^−1^ for the same values (see Supporting Information). Thus, two independent sets of data (derived from Hi-C and ChIP-seq) produce similar results within the permeable moving barrier model framework for rRNA loci and protein-coding loci. Together, these analyses lend strong support for the idea that elongating RNA polymerases can push translocating condensins and supports our findings that condensins bypass the elongating RNAPs with high efficiencies (Fig. 5D).

## Discussion

Collectively, our analysis leads to the permeable moving barriers model (Fig. 5D). The model, which is quantitative in nature, makes several yet-untested predictions. It suggests that by increasing the density of transcribing RNA polymerases beyond a critical value, actively transcribed loci will become directionally impermeable to condensin translocation within physiological time scales. For example, it is highly improbable for a condensin to cross a 10 kb operon with a density of ~20 RNAP/kb, in the head-to-head orientation within ~35 minutes (or one cell division time) but it will bypass the same operon in the head-to-tail orientation within 30 seconds. Prior experimental observations have demonstrated that condensin translocation speed can be directionally slowed down by the rRNA locus (8, 28), and can result in a gene direction dependent build-up of SMC at other highly expressed operons (28), but the impermeability of a transcribed locus to condensin has not been shown. A Hi-C experiment whereby the transcription levels are increased at a specific locus may test the prediction of a directionally impermeable locus, and our quantitative analyses can provide a guide to estimate conditions (based on transcript length and RNAP density) when a locus can totally block condensin translocation.

Predictions of the permeable moving barriers mechanism can also be tested by single molecule experiments similar to those that observed ATP-dependent condensin translocation (9) and loop extrusion (10) *in vitro*. For example, in head-to-head “collision” type of experiments, bypassing of elongating RNAP by condensin, and pushing/pausing of condensins for the predicted ~2 second may be measured; however, while observing the directional effect on a single RNAP on condensin could be difficult, our model suggests that a train of transcribing RNAP will better help measure the permeability value (i.e. the rate at which condensin bypasses DNA-bound obstacles like RNAP). Our analyses of SMC ChIP-seq data and estimates of the permeability rates (i.e. µ ≈ 0.1-0.8 s^−1^ for protein coding sequences versus µ ≈ 0.8-1.6 s^−1^ for rRNA loci) also raise the interesting possibility that different types of genes (i.e. coding versus non-coding) may have different permeabilities. The lower permeability rate in protein coding genes could arise from a higher steric hindrance imposed by ribosomes that bind to nascent RNA during transcription in protein coding, but not rRNA genes. This can be tested *in vitro* by attaching fluorophores or beads of different sizes to a transcribing RNAP, potentially achieving different rates of permeability.

The permeable moving barriers mechanism proposed here for bacteria may be a general mechanism with implications in eukaryotes as well. There is growing evidence that transcription can affect cohesin SMC localization in yeast (47) and mammalian cells (32, 35), and can potentially interfere with the process of cohesin loop extrusion. However, if eukaryotic RNAPs are as permeable to SMC as bacterial RNAPs, transcription in eukaryotes may not have a major effect on chromosome organization by loop extrusion as compared to other molecules that specifically (and directionally) impede loop extrusion, like CTCF (20, 52). Nevertheless, since SMC and CTCF interactions in eukaryotes play a regulatory role in gene expression (53), it is of interest to explore the potential role of SMC-transcription interactions in gene regulation in both bacteria and eukaryotes.

The remarkable ability of loop-extruding condensin SMCs to bypass large elongating transcription complexes is important from both molecular and evolutionary standpoints. While the molecular mechanisms of loop extrusion by SMCs remain to be elucidated, our results suggest that to overcome large steric barriers either: (A) a sufficiently large opening (large enough to fit the entire transcription complex) emerges in the SMC complex lumen during an SMC ATPase cycle, and SMC loop extrusion proceeds by maintaining DNA in a topological embrace (with mechanisms like “inchworm”, “pumping”, “segment-capture”, “shackled-walker” (9, 14, 25, 54–56), or (B) the SMC rings transiently open (57), disengage the topological embrace of the DNA, and re-engage after passing the steric barrier (closer to the “walker”/ “rock-climber” models (39, 40)).

Furthermore, our work suggests a possible link between the SMCs’ ATP hydrolysis rate, and the rate at which the SMC complex can bypass sites of active transcription or other obstacles. We found that condensins pause for about one (or a few) ATP hydrolysis cycles before bypassing RNAP. This suggests that the rate of bypassing will differ between types of SMC complexes depending on their respective ATP hydrolysis rates. For instance, yeast cohesin SMCs are shown to have lower ATP hydrolysis rates *in vitro* of < 0.2 ATP/cohesin/s (58, 59) whereas yeast condensins have higher rates of ~1.5 ATP/condensin/s (9) compared to *B. subtilis* SMC’s 0.7 ATP/condensin/s (29). Indeed, yeast cohesin molecules with an abolished ATP activity accumulate more strongly at sites of convergent transcription (60), consistent with this predicted effect.

Another surprising result is that loop extrusion activity occurs by two effectively uncoupled motor activities *in vivo*; i.e. occlusion of one motor does not affect the translocation of the other. This could indicate that linked dimers of condensins each separately perform directional translocation (thus loop extrusion) (8, 40). We note, however, that this does not preclude the possibility that a single SMC complex performs the two motor activities: for instance, an anchored “one-sided” SMC extruder can alternate the anchoring site and the “DNA reeling arm”, effectively performing two-sided extrusion with uncoupled kinetics (41); this will be a topic of future study.

Irrespective of the molecular details, the ability to overcome steric barriers is likely an important evolutionary adaptation for SMCs to organize chromosomes (e.g. to resolve sister chromatids in bacteria, form domains and compacted genomes in eukaryotes) largely unobstructed by active transcription. Permeability of RNAPs to loop extrusion also suggests that SMCs should be able to effectively bypass other large steric barriers such as nucleosomes in eukaryotes (consistent with experimental evidence, where nucleosome depletion does not affect condensin’s ability to form a mitotic chromosome (61)), as well as long plectonemes (22, 31) and other DNA-bound proteins (as suggested for the SMC homolog MukBEF (62)) in bacteria. It remains to be seen how bigger molecular complexes, e.g. replication machinery, can interfere with the process of loop extrusion.

In summary, our analyses suggest that bacterial condensin’s loop extrusion activity occurs by two effectively independent, and uncoupled motor activities *in vivo*. Further, it appears that two major processes may be at play in shaping the genome-wide condensin trajectories and hence chromosome organization. The first is a transcription independent mechanism that slows down loop extrusion when condensin proceeds towards the origin. The second is a transcription elongation dependent effect at highly transcribed loci like rRNA operons. Most crucially, our models with their inferred parameters, show how SMC translocation speeds can vary as they progress through the genome; we show the speed of extrusion is slowed down by interactions with transcription machinery and depends on the relative directions of transcription and extrusion. Our permeable moving barrier models show that trains of RNA polymerases can serve as directional barriers to extrusion, with individual RNA polymerases having only a modest effect on translocating condensin, by pausing and pushing it back for a mere ~2 seconds at rRNA loci, and ~10 seconds at protein coding loci (Fig. 5D). In all, our work provides a quantitative and predictive framework to study the dynamics of SMC complexes and their interactions with other translocating DNA-bound complexes *in vivo* and *in vitro*.

## Contributions

H.B.B., X.W. and D.Z.R. conceived of the project; H.B.B. processed and analyzed data, developed the theory and models, supervised by L.A.M. and D.Z.R; X.W. made strains, performed ChIP-seq and Hi-C experiments and processed data; P.P. made the RNAP degron strains, performed immunoblotting, Hi-C and microscopy; A.B. developed models and theory. All authors interpreted results. H.B.B and L.A.M wrote the article with input from all authors. Data have been deposited on the Gene Expression Omnibus with accession number GSEXXXXXX.

## Acknowledgements

We would like to thank Johannes Nuebler, Ed Bannigan, John Marko, Stuart Sevier, Jean-Benoit Lalanne, Maxim Imakaev, and Geoff Fudenberg for helpful insights and suggestions. We are especially grateful to Tung Le for sharing data and discussions. H.B.B. acknowledges the Natural Sciences and Engineering Research Council of Canada for a PGS-D fellowship. This work was supported by NIH grants GM086466 and GM073831 (D.Z.R), NSF, Physics of Living Systems (15049420) grant and the Center for 3D Structure and Physics of the Genome of NIH 4DN Consortium DK107980 (L.A.M), and start-up funds from Indiana University (X.W).

---

**Figure S1:**
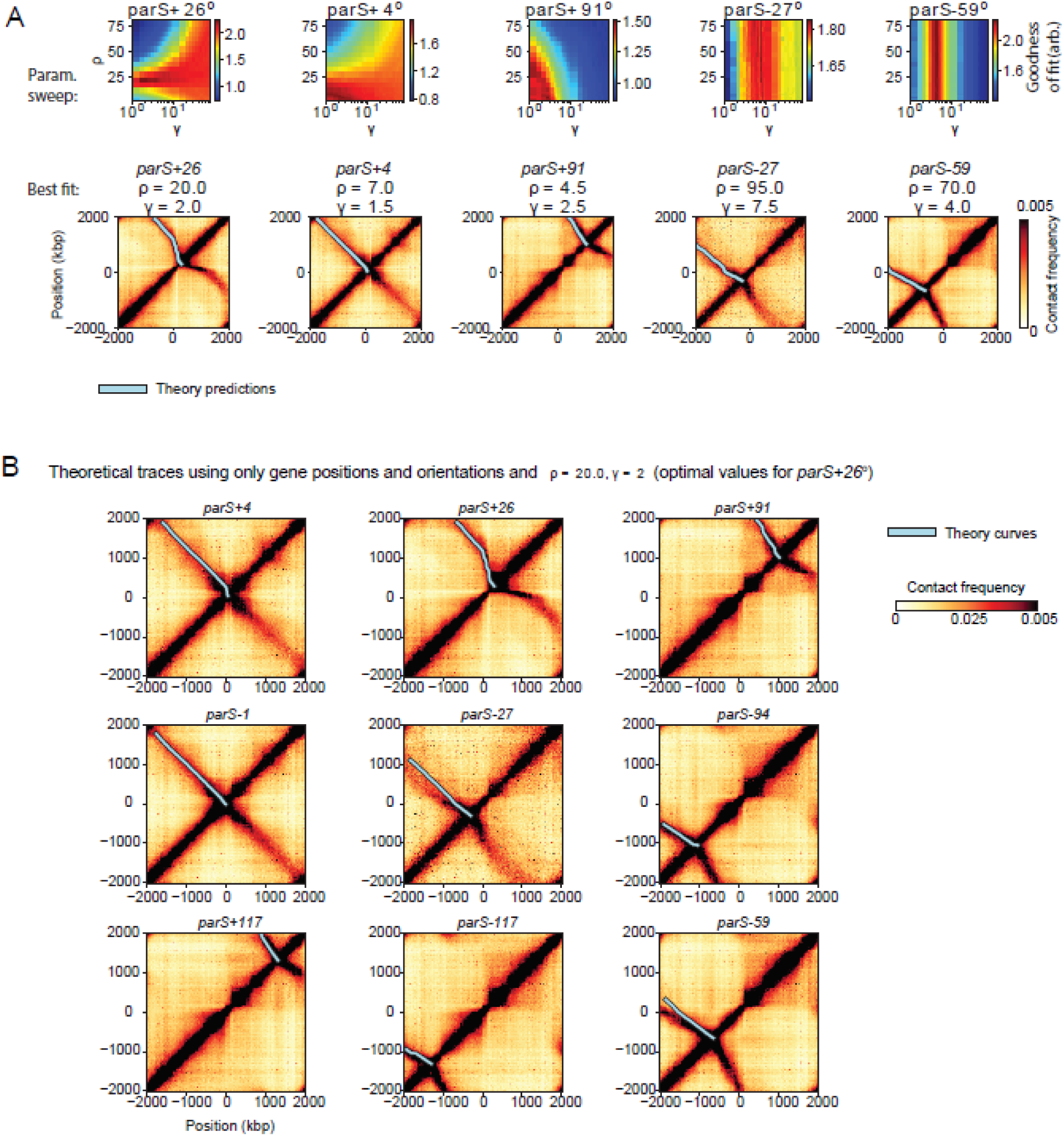
Strain-specific optimal parameters for the gene orientation and direction SMC trajectory model. **(A)** Strain-specific optimal parameter values are obtained from a parameter sweep (top), and the optimal trajectories are superimposed on the respective Hi-C maps (bottom). **(B)** Condensin extrusion trajectories are created using the optimum parameters (*γ* = 2 and *ρ* = 20) for a representative strain (*parS*+26°) to predict extrusion traces in 9 other strains with high fidelity.

**Figure S2:**
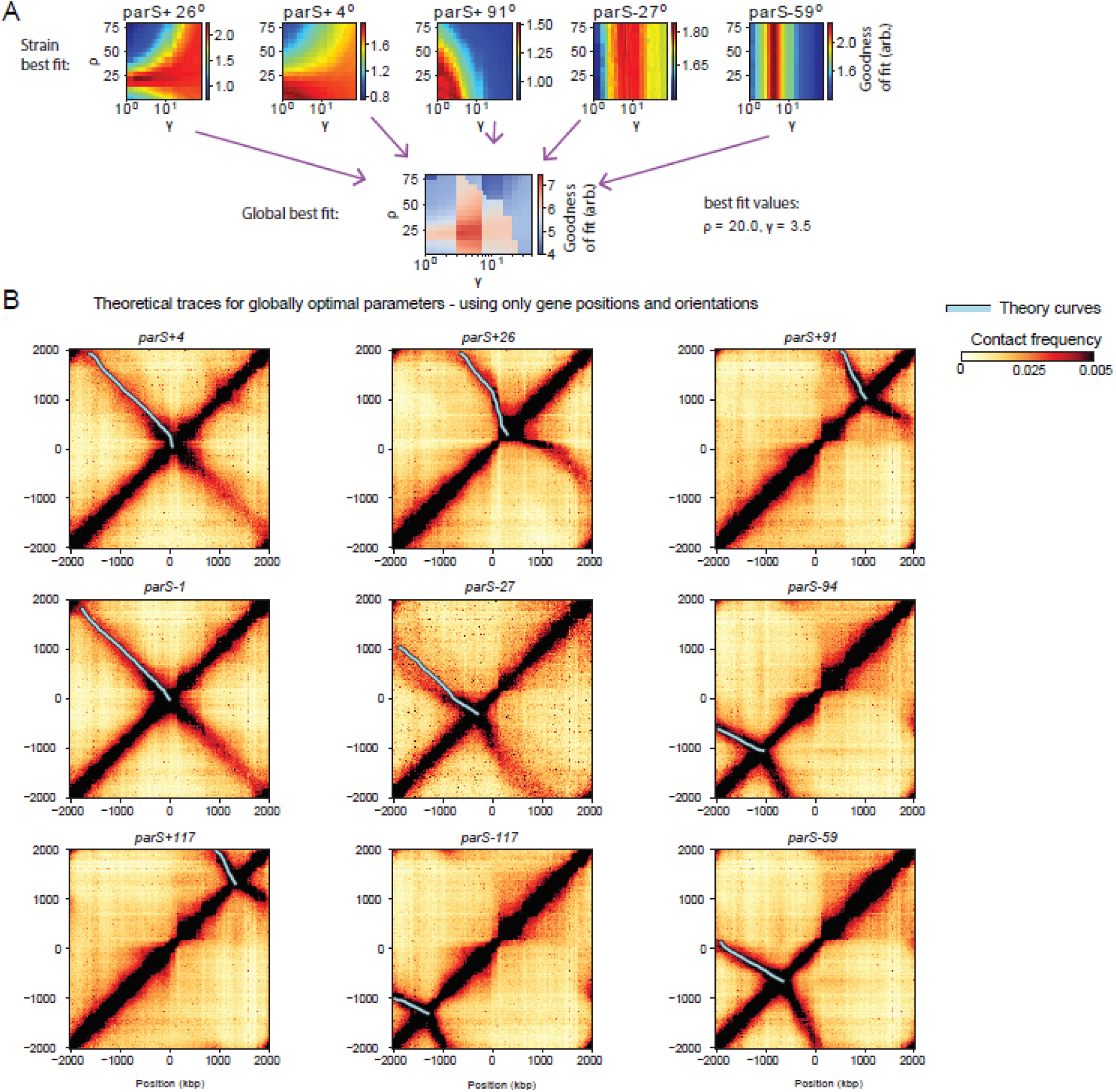
Obtaining globally optimal parameters for the gene position and direction SMC trajectory model. **(A)** Adding together strain-specific parameter goodness of fit values gives the “globally optimal” goodness of fit values (see Supporting Information for details). **(B)** Best-fit (globally optimal) extrusion traces plotted against each strain. The best fit value used for all traces is: *ϒ*=3.5, *ρ*=20 (corresponding to Fig. S2A).

**Figure S3:**
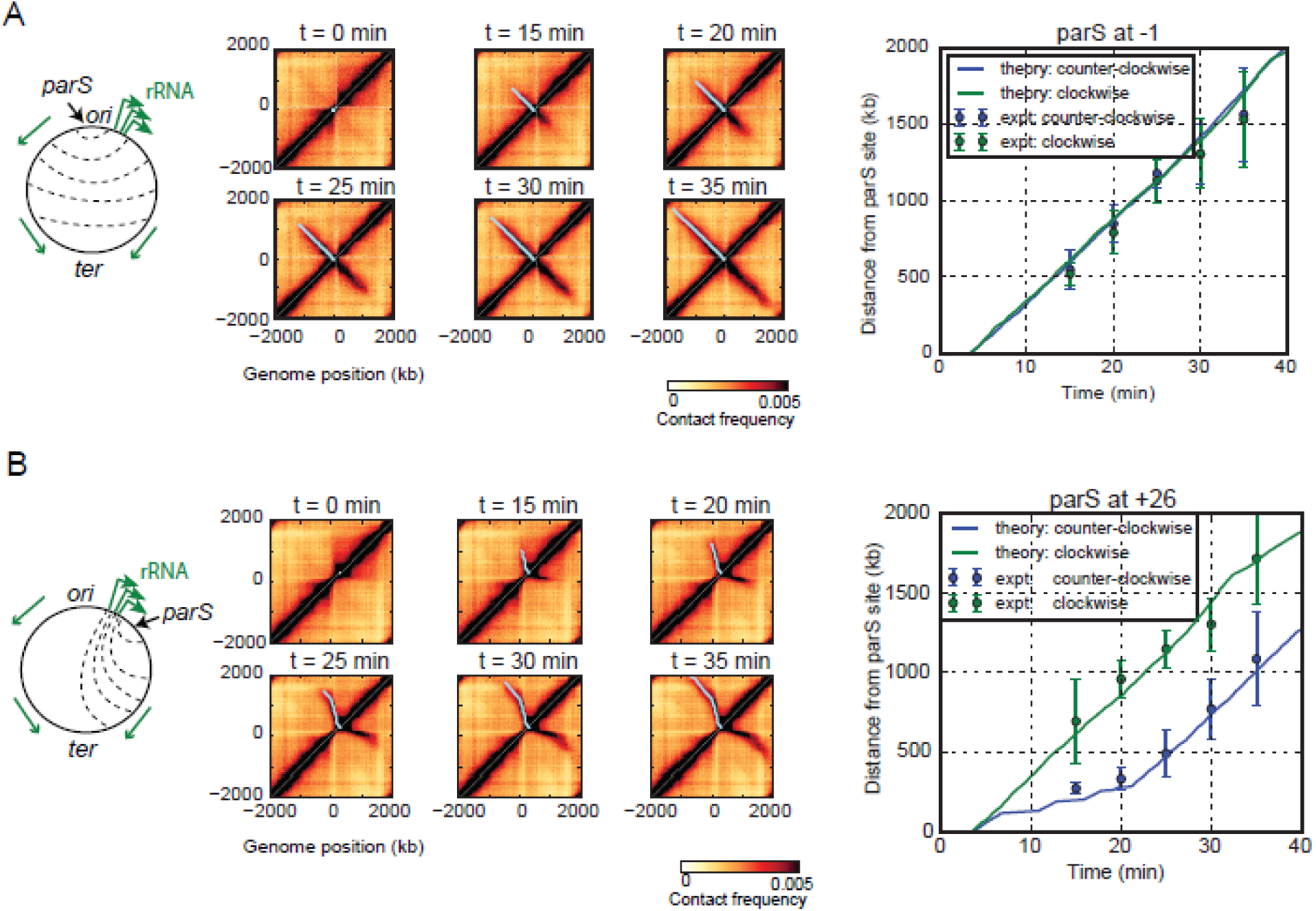
Comparison of gene-position and direction model of SMC translocation to time-course Hi-C data. A comparison of the predicted time-trace of extrusion to experimental data for (**A**) a strain with a single parS site at the −1° position and (**B**) a strain with a single *parS* site at the +26o. Traces were computed using v_avg_ = 50 kb/min (as measured in Wang et al., 2017), and the global optimum parameters as in Fig. S2A. The Hi-C maps show a time-course following induction of ParB (the condensin loader) at t = 0 min. The theoretical trace (light blue line) is superimposed on the Hi-C map. On the right, the model time-course predictions for distance of the extruding motor away from the *parS* site versus time are shown against measurements of extent of juxtaposition (see Wang et al., 2017); the mean and standard deviation of the measured values are displayed for comparison with the theoretical value. Note that the standard deviations do not represent true “errors” on the measurements but are shown to demonstrate the range of expected “maximum extent” values given a set of cutoff thresholds.

**Figure S4:**
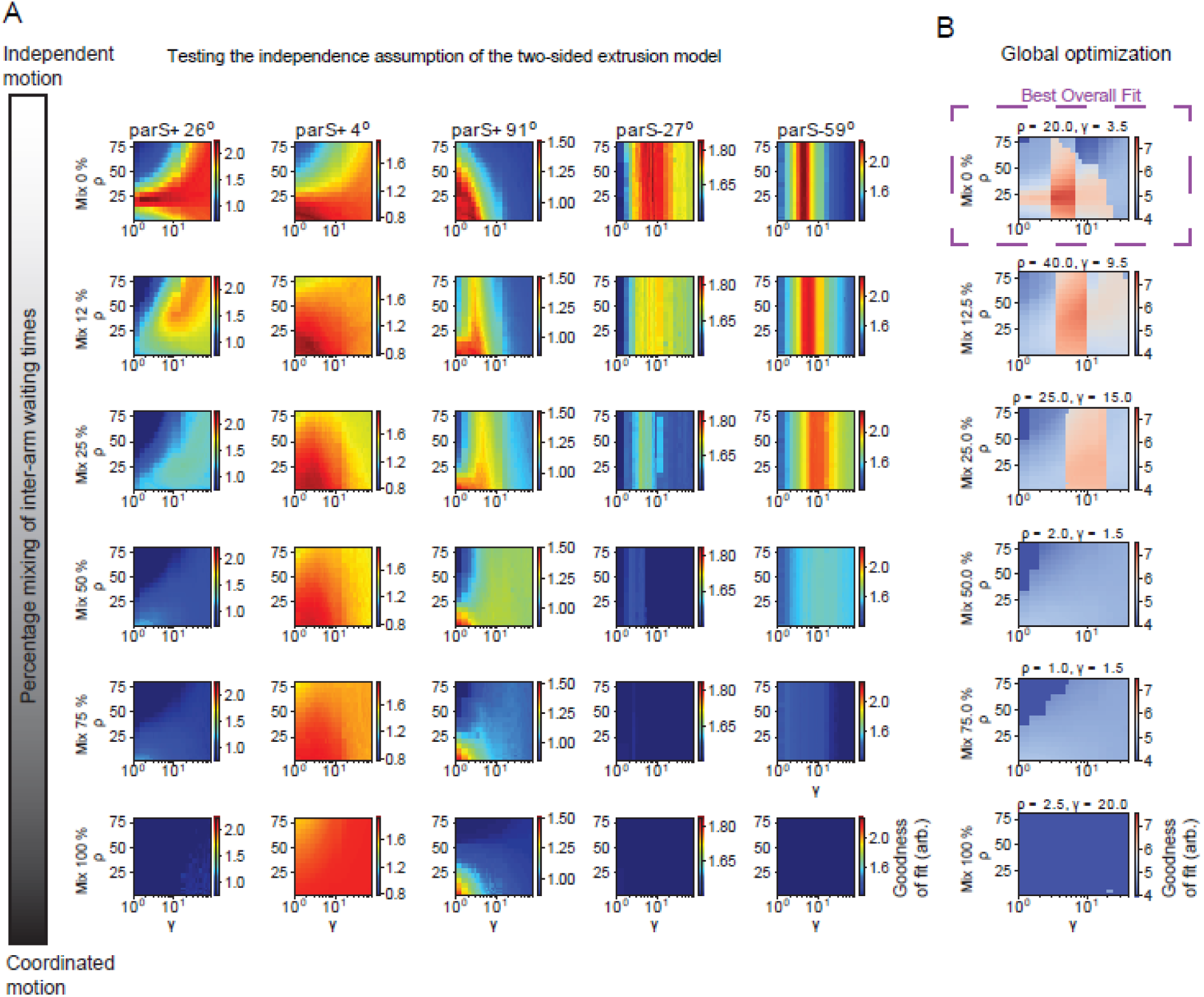
Testing the independence assumption of the two SMC motor activities. (**A**) Parameter sweep (parameters, *γ*, *ρ*) for SMC translocation trajectories calculated using the gene directions and position model with an inter-motor activity “mixing fraction”. Different rows correspond to various degrees of “mixing” interactions (mixing 0% is for independently calculated trajectories of SMC away from the *parS* site; mixing 100% is the perfectly correlated case of identical trajectories). (**B**) The global optimum best fit surfaces for each mixing model calculated using the sum of best fit surfaces for each of the 5 different bacterial strains in panel A; interestingly, the overall best fit occurs for the “mixing = 0%”, which contains both the overall highest, and most defined best fit values (i.e. dark red); the models’ overall goodness of fit decreases progressively from the “Mix=0%” to “Mix=100%”; the overall best fit values for and are indicated above each surface.

**Figure S5:**
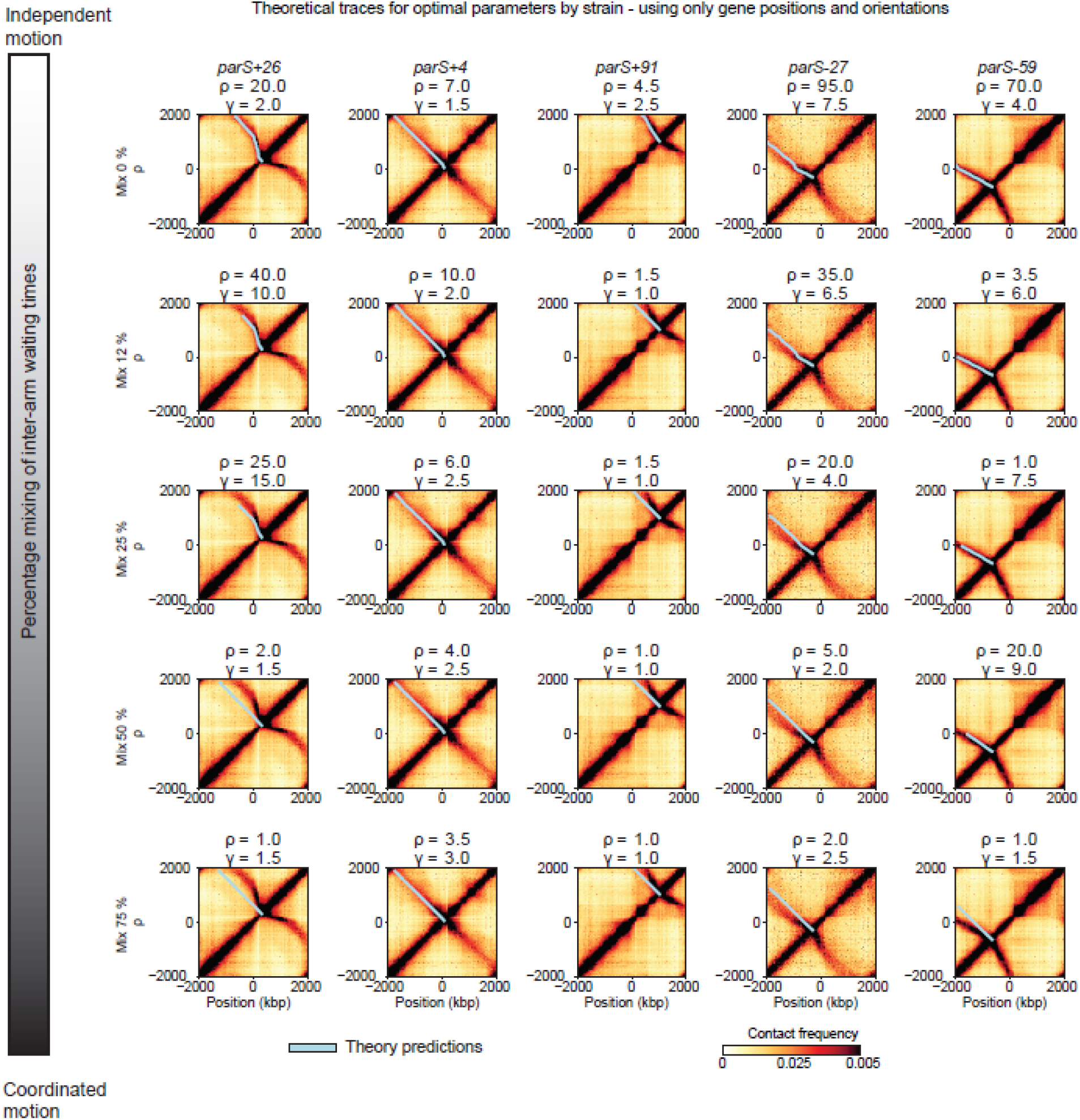
Best-fit (optimal by strain) extrusion traces calculated for the gene position and direction model for different mixing fractions. The best fit values of gamma and rho are indicated above each Hi-C map (corresponding to Fig. S4A panels).

**Figure S6:**
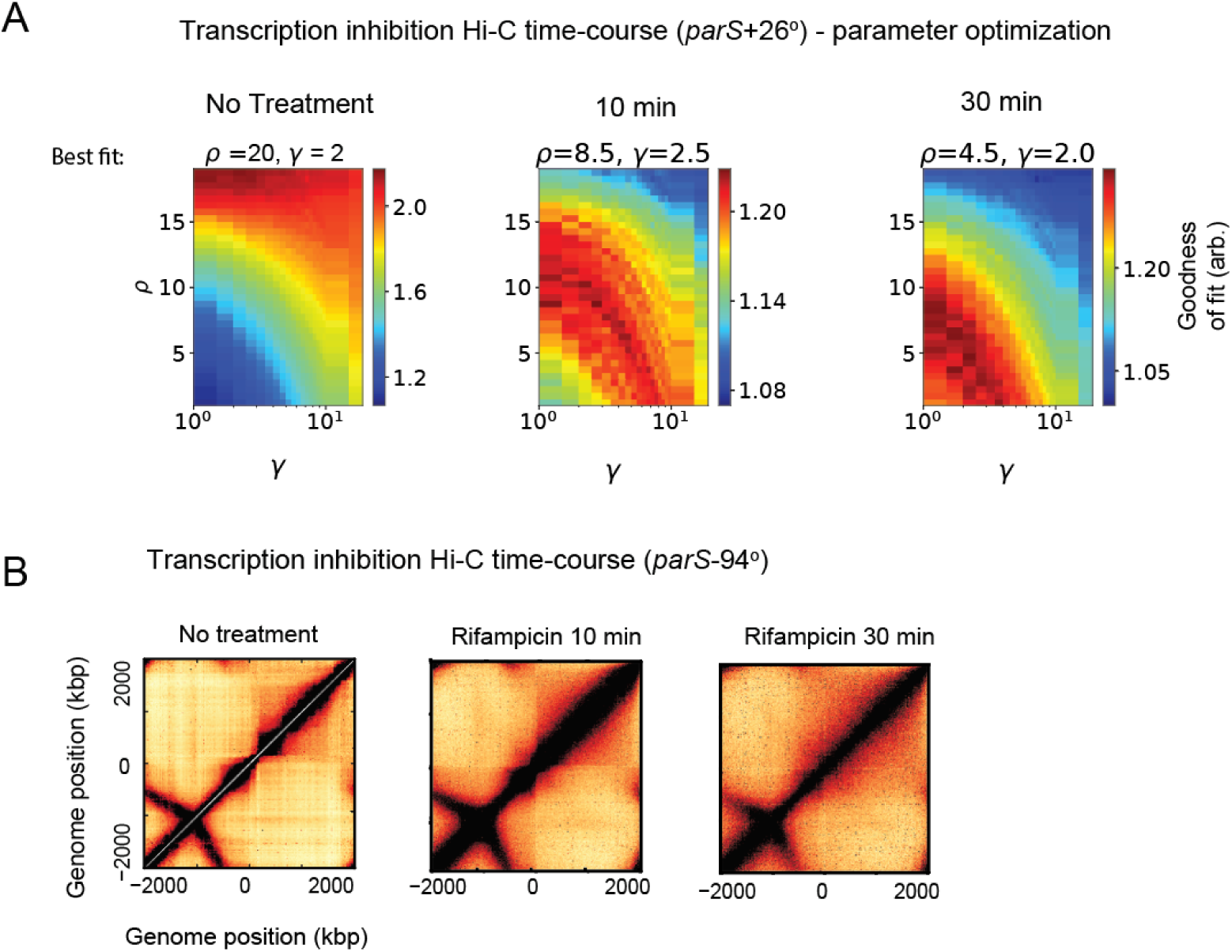
Time-course of transcription inhibition (rifampicin treatment) experiments. (**A**) Parameter sweep for SMC trajectories (gene position and orientation model), following rifampicin treatment of a strain with a *parS* site at the +26° position. Surprisingly, the optimal parameter value for *ϒ* remains largely unchanged, but *ρ* decreases significantly (to a level close to the global optimum value of *ϒ*) (**B**) Time-course following rifampicin treatment for a strain with a *parS* site at the −94° position – no significant changes to the angle of the secondary diagonal are apparent.

**Figure S7:**
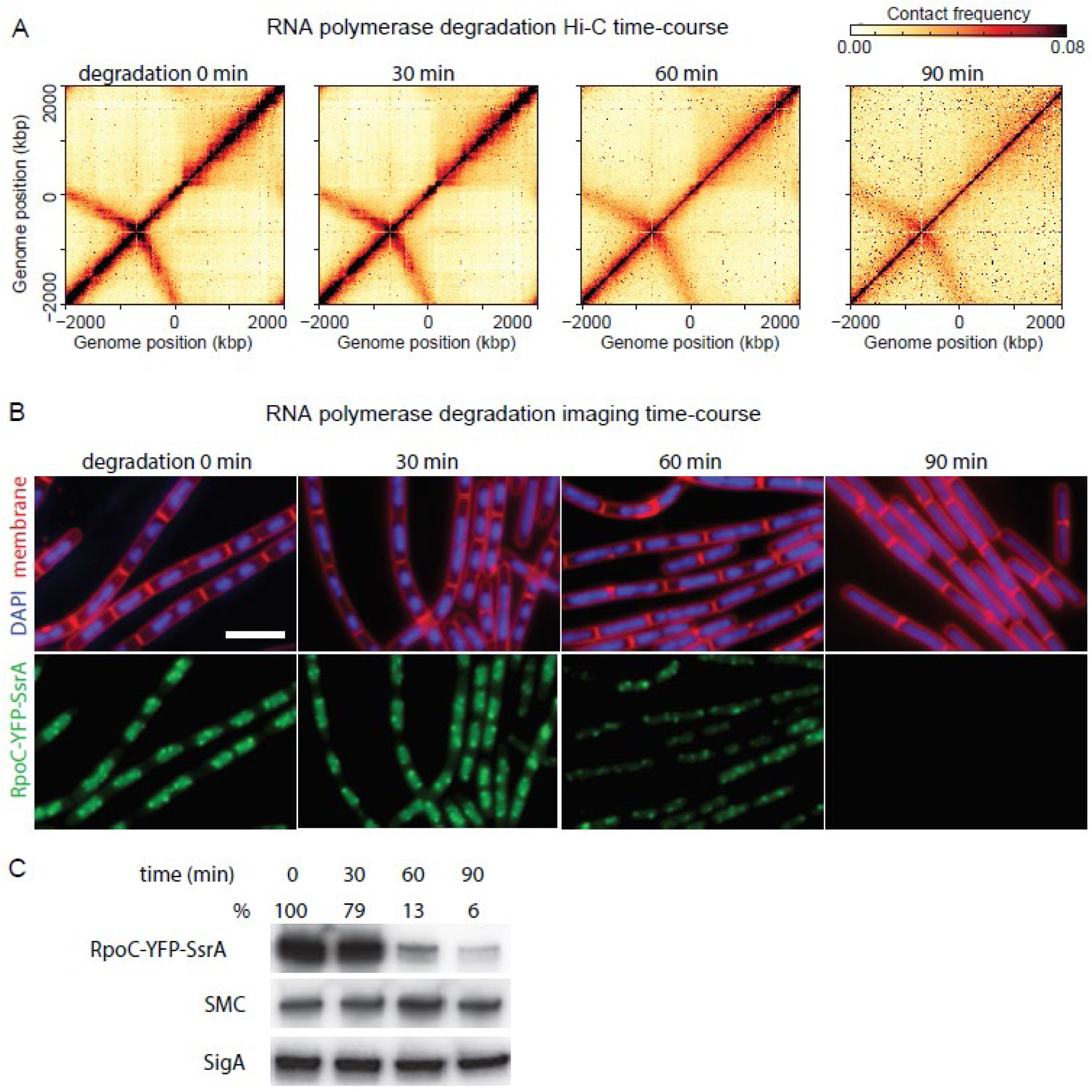
Effect and quantification of RpoC-YFP-SsrA degradation. (**A**) Time-course Hi-C following induction of RNAP (RpoC-YFP-SsrA) degradation. (**B**) Representative images of DAPI-stained nucleoids (blue), FM4-64-stained membrane (red), and RpoC-YFP-SsrA (green) in cells induced for RpoC-YFP-SsrA degradation for the indicated times. The YFP images were using the same scale across all different time points. Similar to rifampicin treatment (Wang et al, 2017), degradation of RpoC-YFP-SsrA caused nucleoids to fill up the whole cell. Bar, 4 µm. (**C**) Immunoblot analysis of RpoC-YFP-SsrA degradation. The same samples were used for Hi-C experiment in (**A**). GFP, SMC and SigA antibodies were used for the three panels respectively. The percentage of remaining RpoC-YFP-SsrA after degradation is indicated. Immunoblots were analyzed using ProteinSimple AlphaView software. The protein levels of SMC and SigA were largely unchanged in the time course of the experiment.

**Figure S8:**
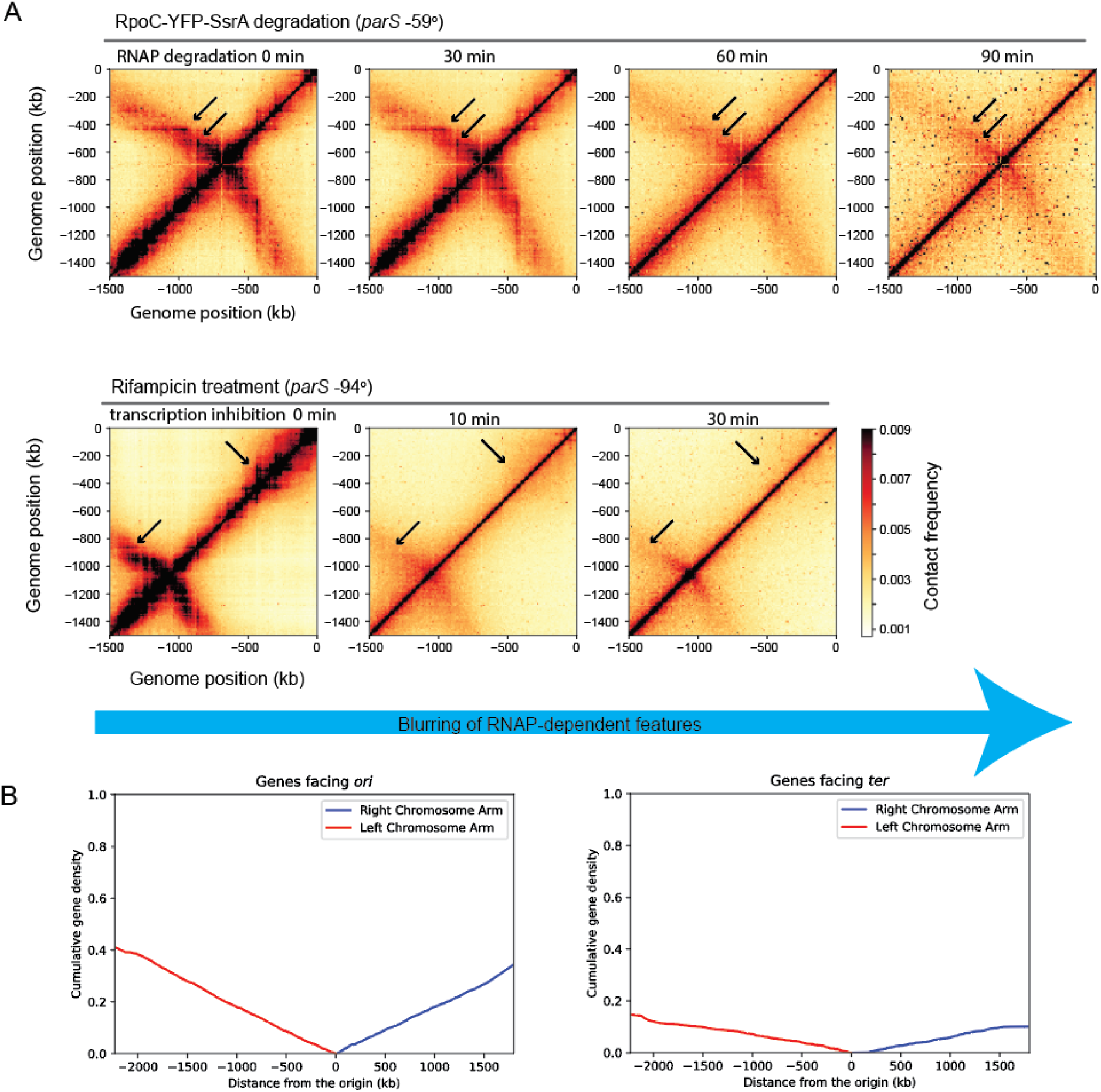
Transcription dependent features on Hi-C maps, and gene orientation density map. (**A**) “Zoomed in” portion of Hi-C map showing features (black arrows) of the Hi-C data which disappear after transcription inhibition or RNAP degradation. Specific “points” of high interaction frequency on the secondary diagonal often co-localize with highly transcribed genes. Features (“X”-shapes and “stripes”) on the primary diagonal largely also disappear in the absence of RNAP or transcription. (**B**) Cumulative density of genes pointing towards the *ori* or *ter* as a function of distance from the *ori*. The uniform increase of the cumulative density demonstrates that the 75% *ori* to *ter* bias in gene directions is largely homogeneous throughout the genome.

**Figure S9:**
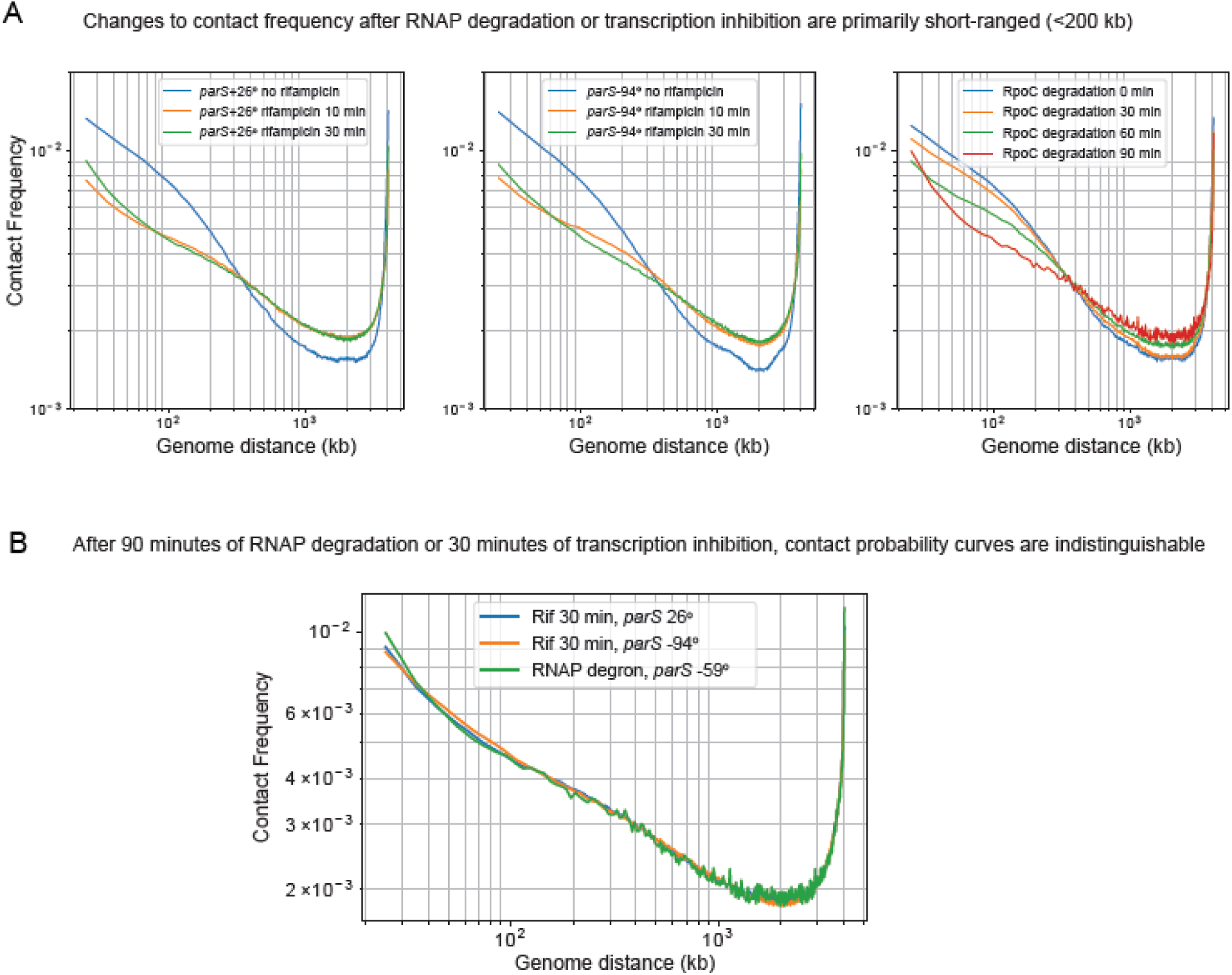
Comparing contact probability curves. (**A**) A comparison of Hi-C interaction frequencies versus genomic distance between pairs of loci in 3 different strains over a time-course of Hi-C data of rifampicin or RNAP degraded cells. (**B**) After rifampicin treatment (30 min) or RNAP degradation (90 min) contact probabilities of 3 different strains become indistinguishable from each other; this suggests that both RNAP degradation and transcription inhibition by rifampicin have similar effects on short-range chromosome contacts (<200 kb).

**Figure S10:**
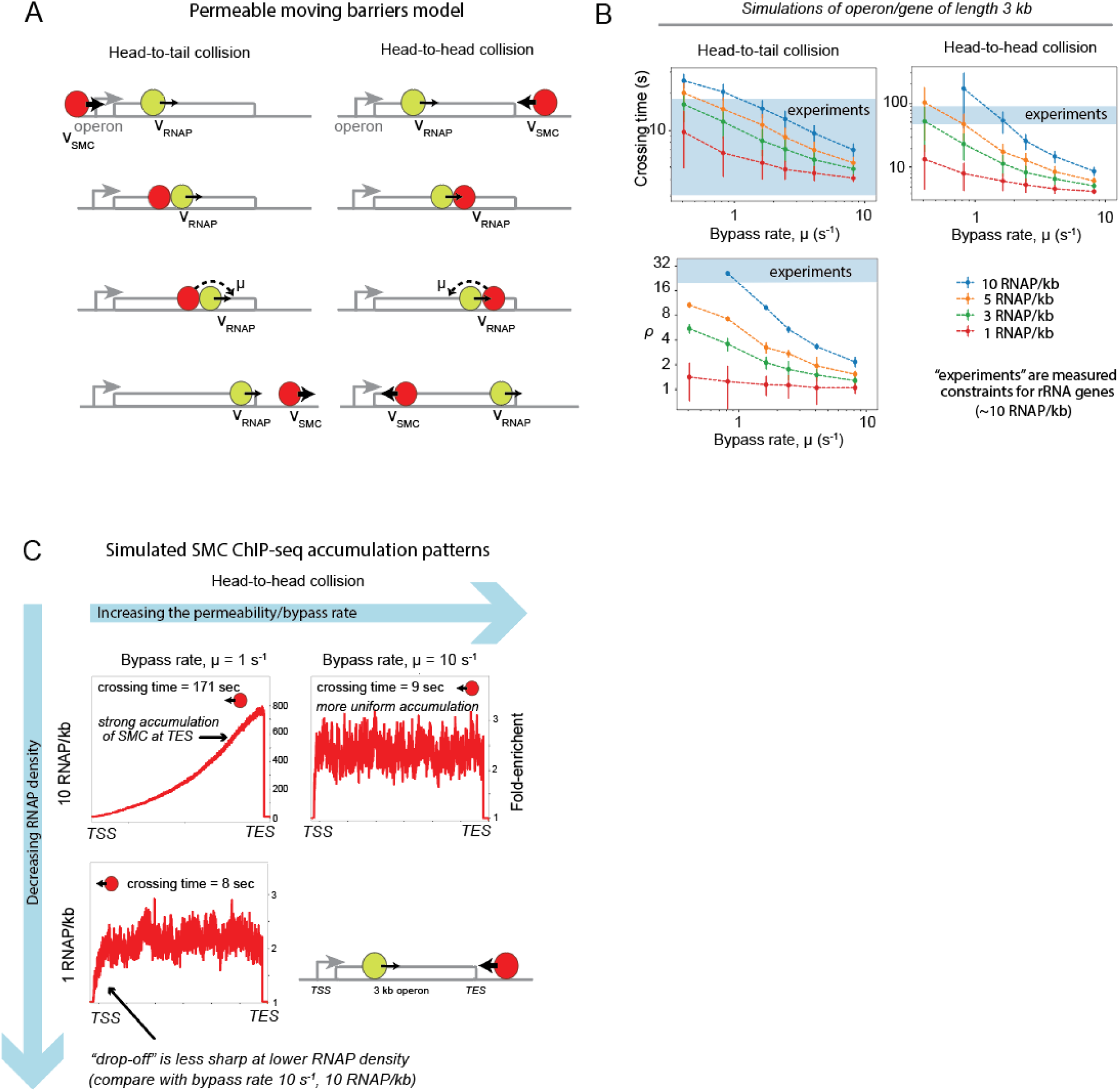
Permeable Moving Barriers Model. (**A**) Schematic illustration showing the moving barriers model with permeable barriers for both the cases of head-to-head and head-to-tail SMC/RNAP interactions. (**B**) Simulations of the times to cross the gene in the head-to-head and head-to-tail cases for a range of RNAP densities on a 3 kb gene. The blue line shows the experimentally inferred range of times to cross an rRNA gene of length 3 kb. From the head-to-head and head-to-tail times, the parameter ρ is also estimated (see Supporting Information); interestingly, the simulations suggest that for rRNA genes, condensins must be able to bypass RNAP at rates between 0.8 s^−1^ and 1.6 s^−1^. (**C**) Simulated SMC accumulation patterns within genes bodies for varying RNAP densities and permeability rates. For RNAP densities and rates below a critical value, the accumulation of SMC is uniform within gene bodies, where transcription end sites are labelled TES and transcription start sites as TSS. In cases of RNAP densities and permeability rates above the critical value, a strong accumulation of SMC is observed at the transcription end sites.

**Figure S11:**
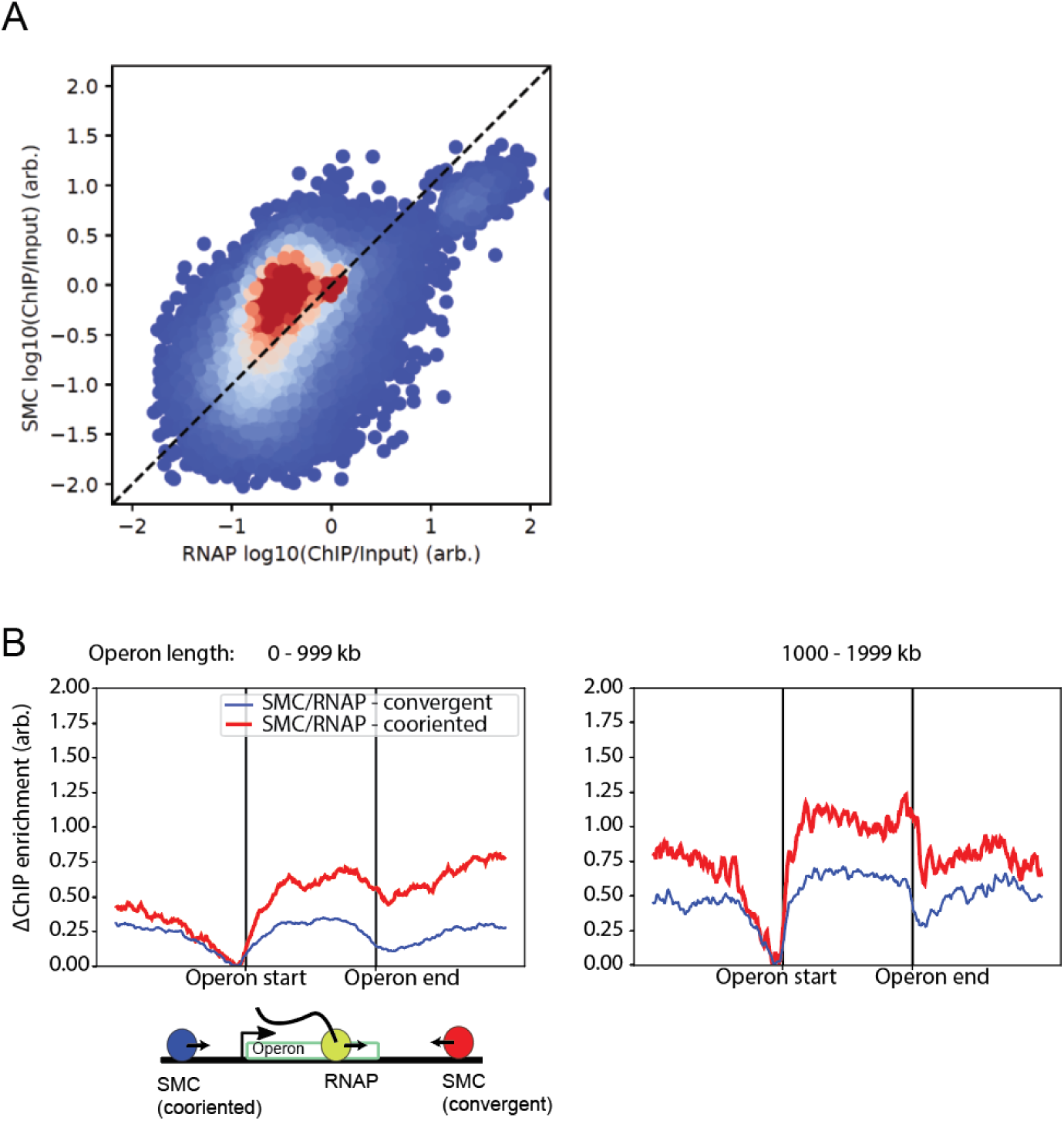
Experimental evidence supporting the moving barriers with permeability model. **(A)** SMC ChIP-seq data plotted against RNAP (RpoC-GFP) ChIP-seq data for the same strain show strong positive correlation between the two values (Pearson correlation R = 0.51, p < 10^−28^)**. (B)** Difference in ChIP-seq enrichment above unity for SMC tracks normalized by RNAP tracks.

## Data sources

**Table S1:** Genome assembly and annotations

| Organism | Usage | Type | Database | Source(s) |
| --- | --- | --- | --- | --- |
| *Bacillus subtilis*, PY79 | Hi-C, ChIP-seq mapping | Genome assembly | NCBI Reference Sequence NC_022898 | Schroeder and Simmons, 2013 |
| *Bacillus subtilis*, PY79 | Extrusion models, ChIP-seq aggregate analysis | Gene annotations | NCBI Reference Sequence NC_022898 | Schroeder and Simmons, 2013 |
| *Bacillus subtilis*, PY79 | Extrusion models, ChIP-seq aggregate analysis | Transcription unit annotations | BioCyc/ BioSubCyc | Karp et al, 2016; Travers et al., 2013; Romero and Karp, 2002 |

**Table S2:** Sources of Hi-C data for Fig. 1

| Figure | Strain | Type | Source |
| --- | --- | --- | --- |
| Figure 1A | PY79 Wild-type | Hi-C | Wang et al., 2015 |
| Figure 1B | BWX3221 (*parS* -94°) | Hi-C | Wang et al., 2015 |

**Table S3:** Sources of Hi-C data and extrusion traces for Fig. 2

| Figure element | Strain | Type | Source |
| --- | --- | --- | --- |
| Figure 2B | BWX3403 (*parS* +26°) | Hi-C | Wang et al., 2017 |
| Figure 2C (top left) | BDR2985 (*parS* +4°) | Hi-C | Wang et al., 2015 |
| Figure 2C (top center) | BWX3268 (*parS* -27°) | Hi-C | Wang et al., 2017 |
| Figure 2C (top right) | BNS1657 (*parS* +91°) | Hi-C | Wang et al., 2015 |
| Figure 2C (bottom left) | BWX3381 (*parS* -117°) | Hi-C | Wang et al., 2017 |
| Figure 2C (bottom center) | BWX3221 (*parS* -94°) | Hi-C | Wang et al., 2015 |
| Figure 2C (bottom right) | BWX3377 (*parS* -59°) | Hi-C | Wang et al., 2017 |
| Figure 2D | BWX3403 (*parS* +26°) | Hi-C | Wang et al., 2017 |
| Figure 2B, 2C, 2D | All strains | Extrusion traces | Transcription unit annotations (Table S1); Calculations (Supplement, Section 3.1) |

**Table S4:** Sources of Hi-C data and extrusion traces for Fig. 3

| Figure element | Strain | Type | Treatment | Source |
| --- | --- | --- | --- | --- |
| Figure 3A (*parS* +26°, left) | BWX3403 | Hi-C | No treatment | Wang et al., 2017 |
| Figure 3A (*parS* +26° right) | BWX3403 | Hi-C | Rifampicin 25ug 30 min | Wang et al., 2017 |
| Figure 3B (*parS* -94° left) | BWX3270 | Hi-C | No treatment | Wang et al., 2017 |
| Figure 3B (*parS* -94° right) | BWX3270 | Hi-C | Rifampicin 25ug 30 min | This work |
| Figure 3 (all panels) | All strains | Extrusion traces | All treatments | Transcription unit annotations (Table S1); Calculations (Supplement, Section 3.2) |

**Table S5:**
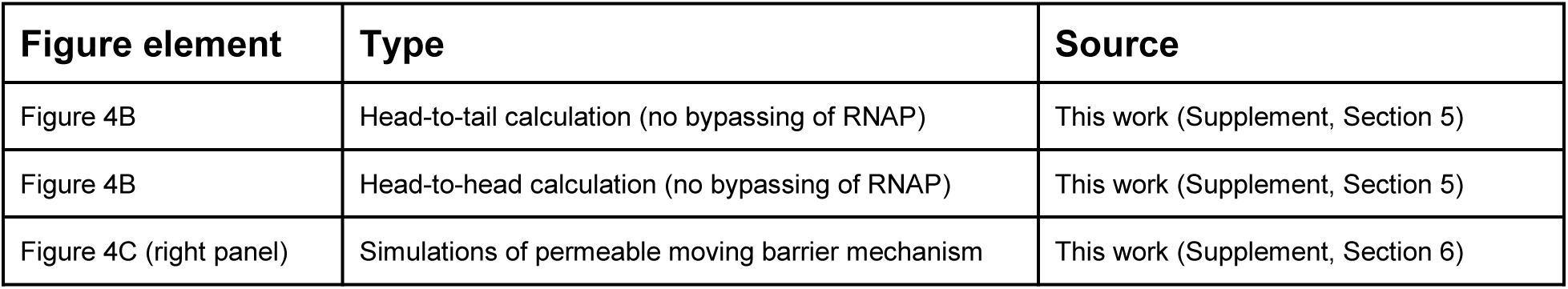
Sources of data and calculations for Fig. 4

**Table S6:** Sources of ChIP-seq data and calculations for Fig. 5

| Figure | Strain | Type | Source |
| --- | --- | --- | --- |
| Figure 5A, 5B (RpoC-GFP) | PY79 Wild-type | ChIP-seq tracks (anti-GFP) | Wang et al., 2017 |
| Figure 5A, 5B (SMC) | PY79 Wild-type | ChIP-seq tracks (anti-SMC) | Wang et al., 2017 |
| Figure 5B | PY79 Wild-type | ChIP-seq aggregate analysis | Transcription unit annotations (Table S1); Calculations (Supplement Section 2.2) |
| Figure 5C | - | Simulations: permeable moving barriers model | This work (Supplement Section 6.4) |

**Table S7:**
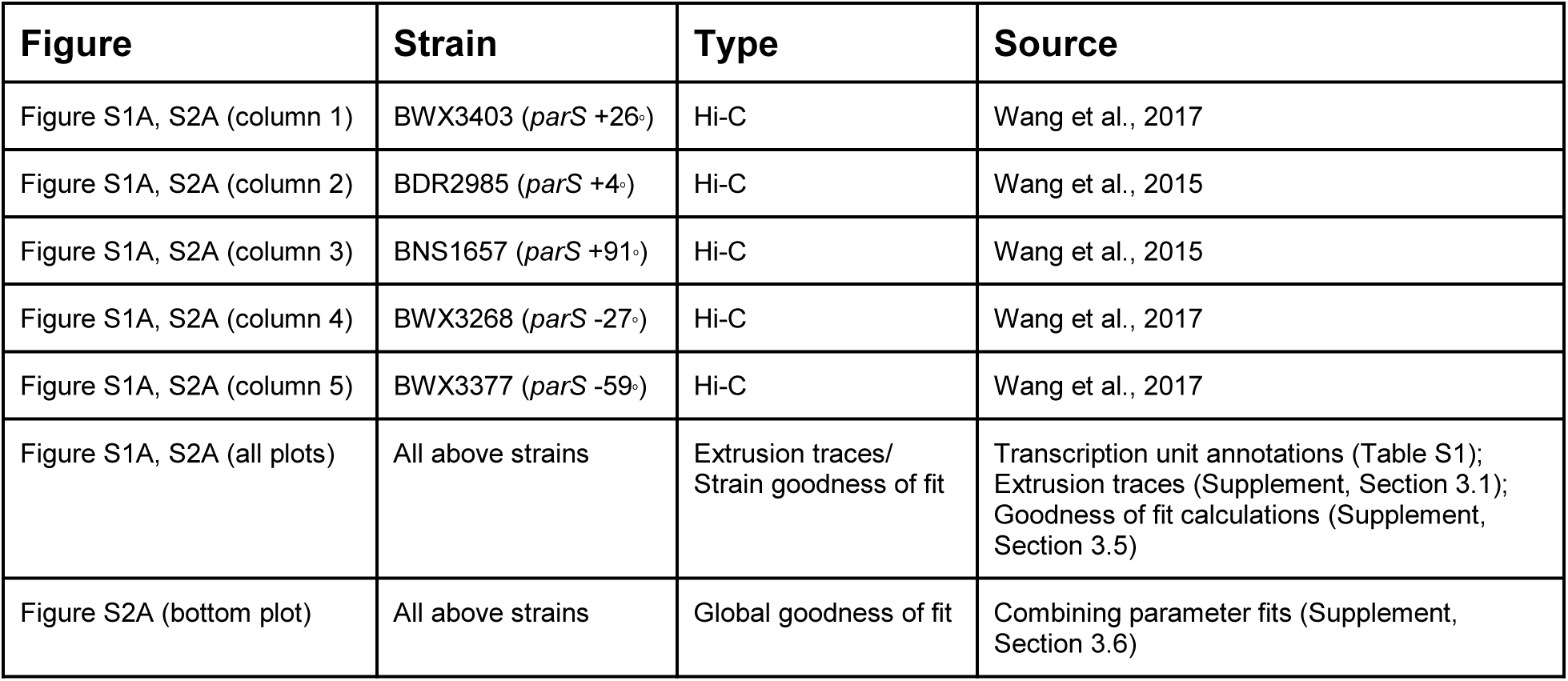
Source of Hi-C data and calculations for Figs. S1A, S2A

**Table S8:** Source of Hi-C data and calculations for Fig. S1B, Fig. S2B

| Figure | Strain | Type | Source |
| --- | --- | --- | --- |
| Figure S1B, S2B (top left) | BDR2985 (*parS* +4°) | Hi-C | Wang et al., 2015 |
| Figure S1B, S2B (top center) | BWX3403 (*parS* +26°) | Hi-C | Wang et al., 2017 |
| Figure S1B, S2B (top right) | BNS1657 (*parS* +91°) | Hi-C | Wang et al., 2015 |
| Figure S1B, S2B (middle left) | BDR2996 (*parS* -1°) | Hi-C | Wang et al., 2015 |
| Figure S1B, S2B (middle center) | BWX3268 (*parS* -27°) | Hi-C | Wang et al., 2017 |
| Figure S1B, S2B (middle right) | BWX3221 (*parS* -94°) | Hi-C | Wang et al., 2015 |
| Figure S1B, S2B (bottom left) | (*parS* +117°) | Hi-C | Wang et al., 2017 |
| Figure S1B, S2B (bottom center) | BWX3381 (*parS* -117°) | Hi-C | Wang et al., 2017 |
| Figure S1B, S2B (bottom right) | BWX3377 (*parS* -59°) | Hi-C | Wang et al., 2017 |
| All panels | All strains | Extrusion<br>traces | Transcription unit annotations (Table S1); Calculations<br>for extrusion traces (Supplement, Section 3.1) |

**Table S9:**
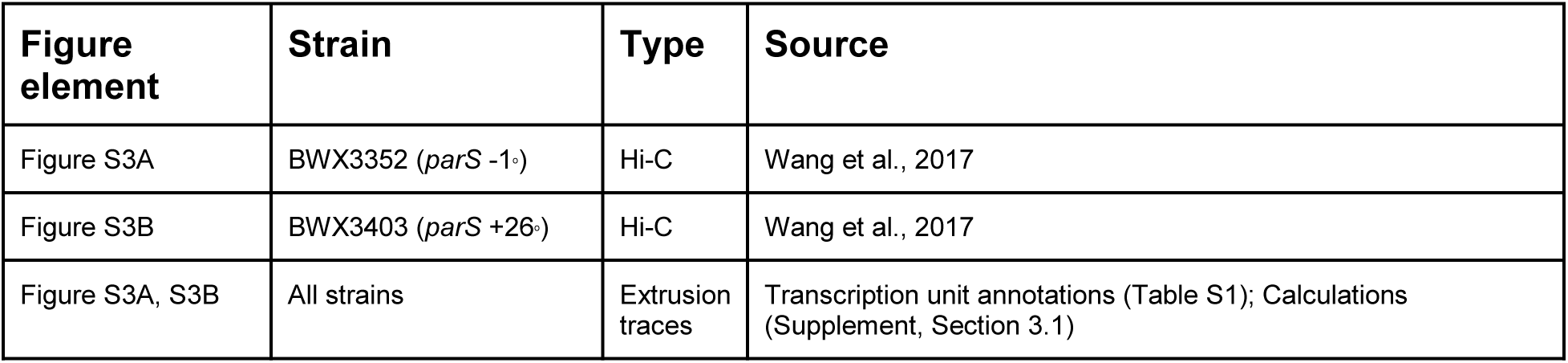
Source of Hi-C data and extrusion traces for Fig. S3

**Table S10:**
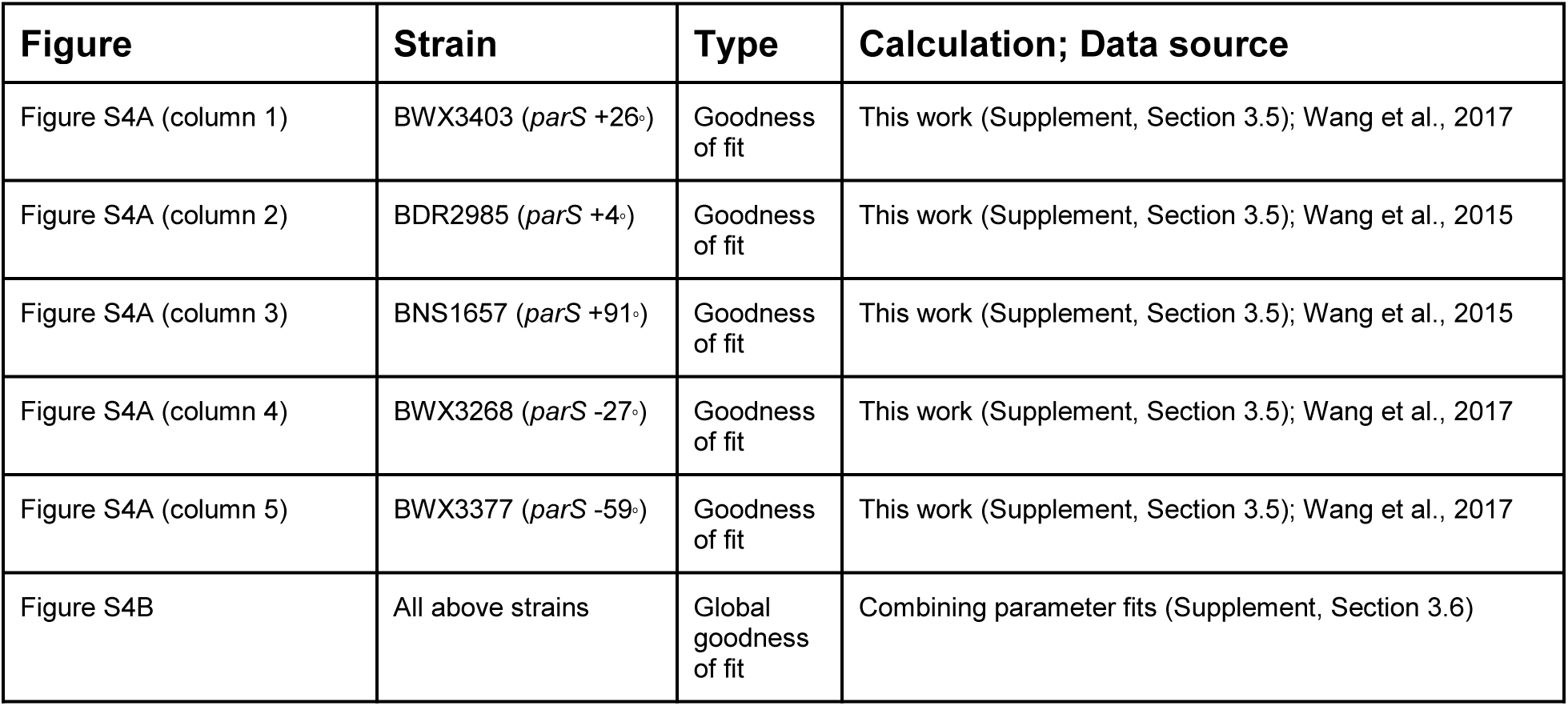
Source of data and calculations for Fig. S4

| **Figure** | **Strain** | **Type** | **Calculation; Data source** |
| --- | --- | --- | --- |
| Figure S4A (column 1) | BWX3403 (*parS* +26°) | Goodness of fit | This work (Supplement, Section 3.5); Wang et al., 2017 |
| Figure S4A (column 2) | BDR2985 (*parS* +4°) | Goodness of fit | This work (Supplement, Section 3.5); Wang et al., 2015 |
| Figure S4A (column 3) | BNS1657 (*parS* +91°) | Goodness of fit | This work (Supplement, Section 3.5); Wang et al., 2015 |
| Figure S4A (column 4) | BWX3268 (*parS* -27°) | Goodness of fit | This work (Supplement, Section 3.5); Wang et al., 2017 |
| Figure S4A (column 5) | BWX3377 (*parS* -59°) | Goodness of fit | This work (Supplement, Section 3.5); Wang et al., 2017 |
| Figure S4B | All above strains | Global goodness of fit | Combining parameter fits (Supplement, Section 3.6) |

**Table S11:** Source of Hi-C data for Fig. S5

| **Figure** | **Strain** | **Type** | **Source** |
| --- | --- | --- | --- |
| Figure S5 (column 1) | BWX3403 (*parS* +26°) | Hi-C | Wang et al., 2017 |
| Figure S5 (column 2) | BDR2985 (*parS* +4°) | Hi-C | Wang et al., 2015 |
| Figure S5 (column 3) | BNS1657 (*parS* +91°) | Hi-C | Wang et al., 2015 |
| Figure S5 (column 4) | BWX3268 (*parS* -27°) | Hi-C | Wang et al., 2017 |
| Figure S5 (column 5) | BWX3377 (*parS* -59°) | Hi-C | Wang et al., 2017 |
| Figure S5 (all plots) | All above strains | Extrusion traces | Transcription unit annotations (Table S1); Mixing model traces (Supplement, Section 3.4) |

**Table S12:** Source of Hi-C data and calculations for Fig. S6

| **Figure element** | **Strain** | **Type** | **Treatment** | **Calculation; data source** |
| --- | --- | --- | --- | --- |
| Figure S6A (left) | BWX3403 (*parS* +26°) | Goodness of fit/ Hi-C | No treatment | Supplement, Section 3.5; Wang et al., 2017 |
| Figure S6A (center) | BWX3403 (*parS* +26°) | Goodness of fit/ Hi-C | Rifampicin 25ug/uL for 10 min | Supplement, Section 3.5; Wang et al., 2017 |
| Figure S6A (right) | BWX3403 (*parS* +26°) | Goodness of fit/ Hi-C | Rifampicin 25ug/uL for 30 min | Supplement, Section 3.5; Wang et al., 2017 |
| Figure S6B (left) | BWX3270 (*parS* -94°) | Hi-C | No treatment | Wang et al., 2017 |
| Figure S6B (center) | BWX3270 (*parS* -94°) | Hi-C | Rifampicin 25ug/uL for 10 min | This work |
| Figure S6B (right) | BWX3270 (*parS* -94°) | Hi-C | Rifampicin 25ug/uL for 30 min | This work |

**Table S13:** Source of data for Fig. S7

| **Figure element** | **Strain** | **Type** | **Treatment** | **Source** |
| --- | --- | --- | --- | --- |
| Figure S7A | BWX4921 (*parS* -59°) | Hi-C | RpoC-YFP-SsrA degradation | This work |
| Figure S7B | BWX4921 (*parS* -59°) | Microscopy | RpoC-YFP-SsrA degradation | This work |

**Table S14:**
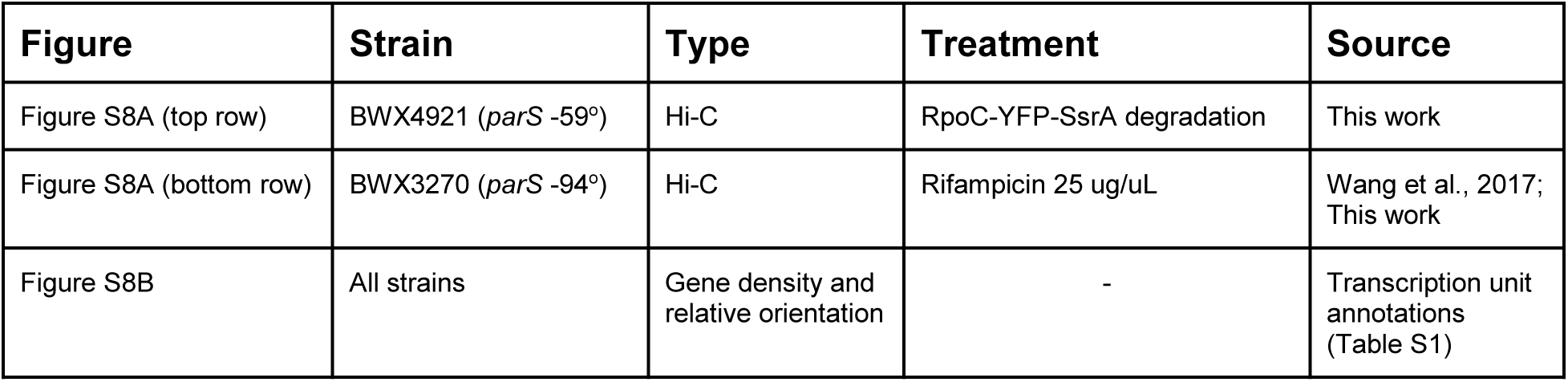
Source of Hi-C data for Fig. S8

| **Figure** | **Strain** | **Type** | **Treatment** | **Source** |
| --- | --- | --- | --- | --- |
| Figure S8A (top row) | BWX4921 (*parS* -59°) | Hi-C | RpoC-YFP-SsrA degradation | This work |
| Figure S8A (bottom row) | BWX3270 (*parS* -94°) | Hi-C | Rifampicin 25 ug/uL | Wang et al., 2017; This work |
| Figure S8B | All strains | Gene density and relative orientation | - | Transcription unit annotations (Table S1) |

**Table S15:**
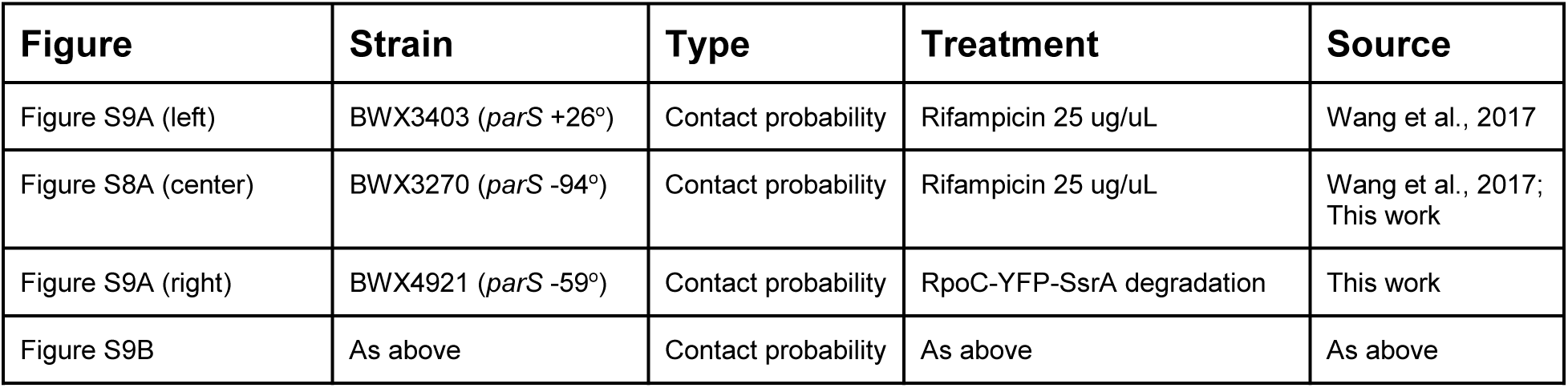
Source of Hi-C data for Fig. S9

**Table S16:** Source of calculations for Fig. S10

| **Figure element** | **Type** | **Source** |
| --- | --- | --- |
| Figure S10A | Locus crossing time calculations | This work (Supplement, Section 6.4) |
| Figure S10A | Estimation of parameter $\rho$ | This work (Supplement, Section 6.5) |
| Figure S10B | SMC ChIP-seq simulations | This work (Supplement, Section 6.4) |

**Table S17:**
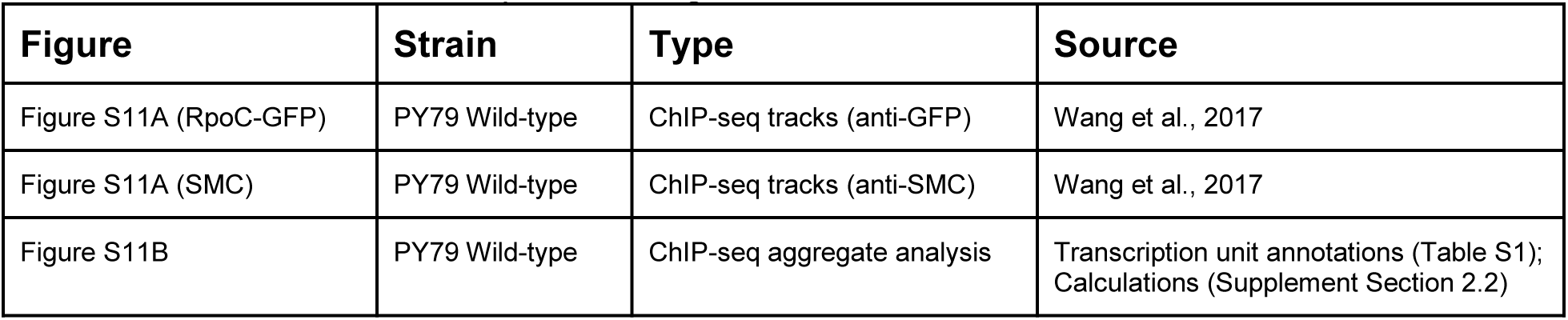
Source of ChIP-seq data for Fig. S11

| **Figure** | **Strain** | **Type** | **Source** |
| --- | --- | --- | --- |
| Figure S11A (RpoC-GFP) | PY79 Wild-type | ChIP-seq tracks (anti-GFP) | Wang et al., 2017 |
| Figure S11A (SMC) | PY79 Wild-type | ChIP-seq tracks (anti-SMC) | Wang et al., 2017 |
| Figure S11B | PY79 Wild-type | ChIP-seq aggregate analysis | Transcription unit annotations (Table S1);<br>Calculations (Supplement Section 2.2) |

## 1 Materials and Methods

### 1.1 Cell growth and treatment conditions

*B. subtilis* strains were derived from the prototrophic strain PY79 (Youngman et al. 1983), and were grown at 37°C in defined rich Casein Hydrolysate (CH) medium (Harwood and Cutting 1990). Hi-C and ChIP-seq experiments were performed for cells in mid-exponential growth phase (optical density of 0.3-0.5 at *A*_600_). Transcription elongation inhibition experiments were performed with rifampicin at a concentration of 25 µg/ml for the indicated minutes.

### 1.2 Hi-C experiments and data processing

The high-throughput chromosome conformation capture (Hi-C) protocol was performed as described (Wang et al, 2015; Wang et al, 2017) adapted as specified from (Le et al, 2013). Briefly, cells were crosslinked with 3% formaldehyde for 30 minutes at room temperature and quenched with 125 mM glycine for 5 min, followed by a PBS buffer wash. Per Hi-C reaction, 5×10^7^ cells were used. Following cell lysis and chromatin solubilization, chromatin was digested with HindIII for 2 hours at 37°C. Digested chromatin ends were filled with Klenow and Biotin-14-dATP, dGTP, dCTP, dTTP at 25°C for 75 minutes. Products were then ligated overnight at 16°C with T4 DNA ligase. Formaldehyde crosslinking was reversed overnight in the presence of proteinase K at 65°C. The DNA was extracted twice with phenol/chloroform/isoamylalcohol (25:24:1) (PCI), precipitated with ethanol, and resuspended in QIAGEN EB buffer. Biotin from non-ligated ends was removed using T4 polymerase for 4 hours at 20°C, followed by extraction with PCI. DNA was then sheared by sonication for 12 minutes with 10 seconds on, 10 seconds off cycles, with 60% amplitude using a Qsonica Q800 water bath sonicator. Sheared DNA was used for library preparation with the NEBNext Ultra kit (E7370S) according to the manufacturer’s instructions for end repair, adapter ligation, and size selection. Biotinylated DNA fragments were purified using 10 µL streptavidin beads. 5 µL of DNA-bound beads were used for PCR in a 50 µL reaction for 14 cycles. PCR products were purified using Ampure beads and sequenced using a Nextseq 500.

Paired-end sequencing reads were mapped to the genome of *B. subtilis* PY79 (NCBI Reference Sequence NC_022898.1) using the same pipeline described (Wang et al., 2015). Contact maps were generated by subdividing the 4,033,459 bp PY79 genome into 404 bins: 403 bins (starting from position 0 bp) contained mapped end reads from contiguous, non-overlapping 10,000 bp stretches of DNA; the final bin contained mapped end reads from the remaining 3,459 bp of the genome. Frequencies of binned paired-end sequence reads were normalized using the iterative correction procedure (Imakaev et al, 2013). Plotting and visualization of Hi-C contact maps were performed using Python 3.6.0 (described below).

### 1.3 ChIP-seq experiments and data processing

Chromatin immunoprecipitation with deep sequencing (ChIP-seq) was performed as described previously (Wang et al, 2015) and modified as specified from (Graham et al, 2014). Briefly, cells were cross-linked using 3% formaldehyde for 30 minutes at room temperature. Cells were then quenched with 125 mM glycine for 5 min, washed, and lysed as described (Graham et al, 2014). Chromosomal DNA was sheared to an average size of 200 bp by sonication using a Qsonica (Q800) water bath sonicator. After removal of cell debris by centrifugation, 50 µL of lysate was removed to serve as an “Input” control. The remaining lysate was then incubated overnight at 4°C with anti-GFP (Rudner et al., 1999) or anti-SMC (Lindow et al., 2002) antibodies and subsequently incubated by Protein A-Sepharose resin (GE HealthCare) for 1 hour at 4°C. After washes and elution, crosslinks in the immunoprecipitate were reversed with an incubation at 65°C overnight. Then, both the “Input” and “ChIP” sample DNA were treated with RNase A, Proteinase K, and extracted using PCI, resuspended in QIAGEN EB Buffer as described (Graham et al, 2014). Library preparation was performed with the NEBNext Ultra Kit (E7370S) and sequenced using the Illumina MiSeq platform. Between 2-5 million reads were collected for each sample.

Paired-end sequencing reads from ChIP and Input samples were mapped to the genome of *B. subtilis* PY79 (NCBI Reference Sequence NC_022898.1) using CLC Genomics Workbench (CLC bio, QIAGEN). To create ChIP and Input tracks from paired-end sequence reads, a count of 1 read was added to all base pairs between the 3’ and 5’ positions of each mapped end (i.e. for a total of 51 counts for each of the paired end, where 51 is the number of sequenced reads from the Illumina machine). Reads that were not uniquely mappable were assigned randomly between the sites in question. In this way, we could assign reads to repetitive sequences such as the ribosomal rrn loci to give an estimate for the occupancy at those loci. For plotting, every sample was first normalized to the total number of reads. ChIP-enrichment was then calculated from the ratio of ChIP-seq signal over the Input signal (ChIP/Input). Normalization, subsequent processing and plotting of ChIP-seq data were performed using Matplotlib v2.2.2 (https://matplotlib.org/), Numpy 1.13.1 (https://www.numpy.org/) and Scipy 0.19.1 (https://www.scipy.org/) in Python 3.6.0 (https://www.python.org/).

### 1.4 Microscopy experiments and data processing

Fluorescence microscopy was performed using a Nikon Eclipse Ti2 microscope equipped with Plan Apo 100x/1.45NA phase contrast oil objective and an sCMOS camera. Membranes were stained with FM4-64 (N-(3-Triethylammoniumpropyl)-4-(6-(4-(Diethylamino) Phenyl) Hexatrienyl) Pyridinium Dibromide, Molecular Probes) at 3 µg/ ml. DNA was stained with DAPI (4’,6-diamidino-2-phenylindole, Molecular Probes) at 2 µg/ml. Images were cropped and adjusted using MetaMorph software. Final figure preparation was performed in Adobe Illustrator.

### 1.5 External data sources

See Tables S1–S17.

## 2 Data processing

### 2.1 ChIP-seq data visualization and aggregation over operons

ChIP-seq data displayed in the main figures were plotted as follows. We first computed the ratio for ChIP/Input data, and then removed any “NaN” or “Inf” values; for such values, we set the number scale for these regions to 1. To display the “raw” ChIP/Input data (Fig. 5A), we used a 1-D Gaussian filter with a window size of 1 kb (using Scipy’s scipy.ndimage.gaussian_filter, with sigma=1000).

Averaging of the ChIP-seq signal over genes and operons was performed by first identifying the start and end locations of each feature. Features were obtained from annotations (genome or transcription unit annotations) as in Table S1. We first classified features as cooriented or convergent with the SMC extrusion direction (i.e. head-to-head, or head-to-tail). Then, we applied the analysis separately for each of these two cases using the procedure below.

For each unique feature, we calculated its length, *L*, by taking the difference between the annotated start and end positions. We then defined a window of total length 3*L* centered on the feature’s mid-point (i.e. *L* basepairs upstream, followed by the feature of length *L*, and *L* basepairs downstream), and obtained the capped ChIP/Input signal for that region. Next, we coarse-grained the signal in that region by re-binning it into 999 distinct bins (i.e. 333 bins for signal downstream, 333 bins for the feature, and 333 bins upstream); in this way, we can average the capped ChIP/Input signal for differently sized genes or operons.

Re-binning was done using the function zoomArray found in mirnylib/numutils.py using the default parameters (http://bitbucket.org/mirnylab/mirnylib, 2018-02-01). Briefly, it works by block-averaging the signal: Let’s first define the desired final length of the array as 3*L*, and the input region length as *A* (assuming *A* ≥ 3*L*). Then, if *A* is an integer factor of 3*L*, we simply coarse-grain the capped ChIP/Input signal by averaging it over consecutive, non-overlapping bins of length 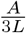 (i.e. for an integer bin *x* of the final array, we average over bins 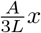 to 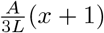 in the input array). In the cases where length *A* is not initially an integer factor of 3*L*, we apply a cubic spline interpolation to the input signal such that the final length of the interpolated signal region is integer divisible by 3*L*. Interpolation is performed using the scipy.ndimage.zoom function; the applied zoom factor is the value *zoom* = ┌*A/L*┐ (i.e. *A/L* rounded upwards to the nearest integer). Following interpolation of the capped ChIP/Input signal to an integer length of 3*L*, we block average the signal as described above.

After coarsegraining all desired features to a final length of 999 bins, we averaged the results using the Numpy nanmean function, and plotted the results with Python’s matplotlib v2.2.2.

### 2.2 Hi-C data visualization

All Hi-C data displayed in the paper were shown with a linear colour scale. Unless specifically indicated in the colour bar, all Hi-C contact frequencies were thresholded to a value of 0.005 to improve clarity and visibility of the secondary diagonal for display purposes. Additionally, although the most proximal 30 kb along the main diagonal of the Hi-C map was masked out for the iterative correction procedure, as is typical for bacterial and eukaryotic Hi-C data processing (Le et al., 2013; Imakaev et al, 2012), the masked-out values were filled in with the maximum of the colour scale (i.e. 0.005 or otherwise) for the purposes of display to improve the visibility of the data and avoid map rendering artefacts due to empty pixels.

We opted to display Hi-C maps with the *ori* at the center, with the main diagonal going from the bottom left to top right. As such, all contact matrices are re-centered using Numpy’s roll() function separately along each dimension, and flipud(). i.e. for a contact matrix, M, and numpy imported as np, we call M = np.flipud(np.roll(np.roll(M,-shiftBy,axis=0),-shiftBy,axis=1)). Specifically for *B. subtilis* PY79, the Hi-C contact map (M) is a 404×404 numpy array and we shift/re-center the map by 202 bins (i.e. with shiftBy=202).

### 2.3 Transcription units from predictions

Transcription unit annotations used in this study (see Table S1) were acquired from the BioCyc database (Karp et al., 2016). First, we obtained the transcription units using the database SmartTables (Travers et al., 2013). Transcription units were computationally predicted using software previously described (Romero and Karp, 2002) for the *B. subtilis PY79* genome and others. The SmartTable was configured first to list the genes of the transcription unit, and the right- and left-end positions of those genes. SmartTables were exported to a spreadsheet, and subsequent spreadsheet manipulations/parsing were performed using Python and the Pandas v0.22.0 data analysis library (https://pandas.pydata.org/). To determine the transcription unit start and end positions, we searched for the maximum and minimum postions of all the genes in the list. The orientation of the transcribed unit was determined by the orientation of the genes within the transcription unit; if the right-end positions of the genes were greater than the left-end positions, then we called these “forward facing”, otherwise, if the left-end positions were greater than the right-end positions, these were annotated as “backwards facing” operons.

## 3 Models of SMC loop extrusion

### 3.1 Calculating secondary diagonal traces from gene directions and positions

For the gene positions and directions extrusion model, we first start with computing a quantity we call the D-score; the D-score is the local gene density multiplied by the relative direction of transcription to condensin translocation. We separately compute a D-score for each genomic feature: i.e. one for operons (or genes) (*D*_*operon*_), one for rRNA loci (*D*_*rRNA*_), and one for tRNA loci (*D*_*tRNA*_). We will illustrate with an example using operons (e.g. *D*_*operon*_), but a similar procedure applies to rRNA and tRNAs annotations.

To compute *D*_*operon*_ from operon genome annotations, we first initialize an array of zeros of size equal to the genome length (in basepairs). For each operon, we obtain its start and end positions and add +1 for “forward” facing operons, or −1 for “backward” facing operons at each base pair position belonging to an operon; forwards and backwards are in reference to the linear genome coordinate and the relative translocation direction of condensin (i.e. forwards = “head-to-tail” type encounters, backwards = “head-to-head” type encounters). We apply a similar procedure for *D*_*tRNA*_ and *D*_*rRNA*_.

We then make the D-scores mutually exclusive (i.e. non-overlapping). Since operon annotations may overlap with rRNA annotations or tRNA annotations, for each basepair (or array position), *i*, we set the *D*_*operon*_(*i*) = 0 if *D*_*tRNA*_(*i*) ≠ 0 or *D*_*rRNA*_(*i*) ≠ 0. Next, we set *D*_*tRNA*_(*i*) = 0 if *D*_*rRNA*_(*i*) ≠ 0. This procedure is done to avoid double-counting the times to cross the loci that are multiply annotated (i.e. by operons, tRNAs and/or rRNAs).

From the D-score, we can then compute the relative locus crossing times for the clockwise and counter-clockwise directions using the parameter, *γ*, as defined in the main text. For each basepair (or array position), *i*, the traversal time is:

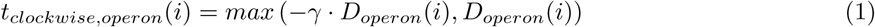

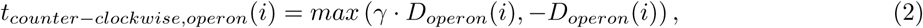

The clockwise and counter-clockwise traversal times for condensin to cross an rRNA or tRNA feature are computed similarly; for rRNA, we replace *D*_*operon*_ with *D*_*rRNA*_ and we replace *γ* by *ρ*; for tRNA, we use *D*_*tRNA*_ and use *γ* as we did for operons since although these are also highly transcribed loci, it was shown that due to their short lengths, they have little effect on local genome structure (Le & Laub, 2016).

The total locus traversal time at position *i* (where *i* is defined as an index along the genome coordinate) for a particular direction of condensin translocation is evaluated as the sum of traversal times for all the features at that locus:

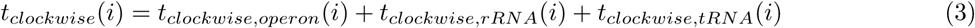

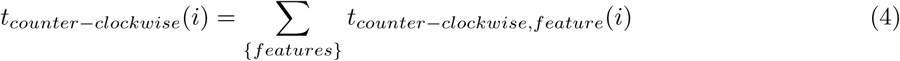

The total cumulative time as a function of distance, *x*, from the *parS* site is the cumulative sum of Eqs.3 and 4, starting from the *parS* position (i.e. *i* = *parS*):

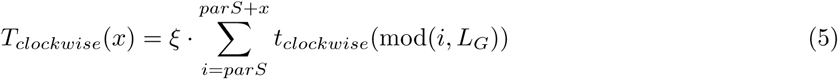

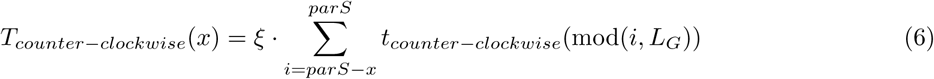

The total genome length is *L*_*G*_; mod() is the modulus function, which allows our index to run continuously along the circular genome (i.e. when the index position *i* reaches *L*_*G*_, the genome “end”, the next value is automatically *i* = 1). The calibration constant, *ξ*, is used to convert relative to real times (also descibed in detail in Section 4.1) and is computed using,

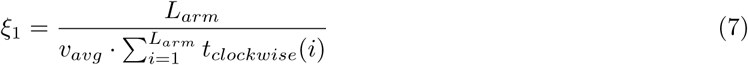

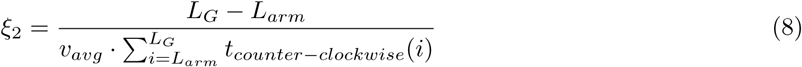

and,

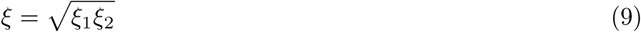

*L*_*arm*_ ≈ *L*_*G*_/2 ≈ 2 Mb is the length of a genome arm, and *v*_*avg*_ ≈ 50 kb/min is the experimentally measured average rate of condensin chromosome arm juxtaposition (Wang et al., 2017). We note here that *t*_*clockwise*_(*i*) and *t*_*counter−clockwise*_(*i*) are unitless (they are relative times), therefore *ξ*, *ξ*_1_ and *ξ*_2_ have units of time.

Eqs.5–9 allows us to plot time versus distance from *parS* site graphs for each extrusion motor direction. However, to overlay the predicted trajectory of the extrusion motor pair in time on the Hi-C map, we first need distance versus time graphs (and ultimately position versus time). Thus, we first invert Eqs.5 and 6. This is done numerically by linear interpolation (using Numpy’s interp function). We query a set of 10,000 time steps evenly sampled from 0 min to 50 min separately for both *t*_*clockwise*_(*x*) and *t*_*counter−clockwise*_(*x*). The interpolation results in matched pairs of spatial displacements (*x*_*clockwise*_(*t*), *x*_*counter−clockwise*_(*t*)) from the *parS* site at each of the queried time points, *t*. We note that (*x*_*clockwise*_(0), *x*_*counter−clockwise*_(0))=(0,0). Therefore, to plot/superimpose the predicted extrusion trajectory on the Hi-C contact map, we convert the distances into 2-D matrix positions (X,Y) using (X,Y)=(*parS* + *x*_*clockwise*_(*t*), *parS* − *x*_*counter−clockwise*_(*t*)).

### 3.2 Calculating secondary diagonal traces with “mixing” (i.e. non-independent SMC motors)

Unlike in the case of calculating the secondary diagonals of pairs of independently translocating SMC motors, extrusion traces cannot be easily pre-computed in the mixing model. We used the following algorithm to compute the “mixing model” trajectories. D-scores were computed as was done for in Section 3.1, and mixing times were calculated on an SMC step-by-step basis. D-scores give us the waiting time distributions for each one of the pair of SMC motors (i.e. the clock-wise and counter-clockwise moving motors).

We started the trajectory calculation by arbitrarily choosing one of the SMC motors to step forward first by one base-pair. We chose the clockwise moving motor (but this makes an unmeasurable difference for the overall trajectory at the observable distances). We define *T*_*clockwise*_(*i*) and *T*_*counter−clockwise*_(*j*) as the cumulative times for the clockwise motor and counter-clockwise motors respectively to reach genome positions *i* and *j*. We initialize these quantities as follows:

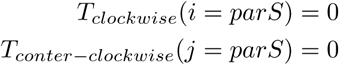

We calculated the cumulative time, *T*_*clockwise*_ required for the clockwise motor to reach the to take a 1 bp step using the mixing formula, where *f*_*mix*_ (which ranges from 0 to 1) is the degree to which each SMC motor is impaired by the other:

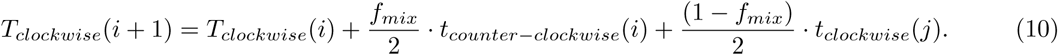

We also increase the position counter, *i*, for the clockwise motor by +1 (i.e. *i* ← *i* + 1). In all the subsequent steps, we check whether *T*_*clockwise*_ > *T*_*couner−clockwise*_ at the current positions of *i* and *j*. If so, then we update the counter-clockwise motor using:

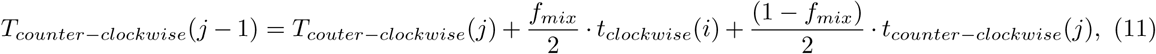

and the counter-clockwise position index (*j* ← *j* − 1). If not (i.e. *T*_*clockwise*_ < *T*_*couner−clockwise*_), we calculate the time to take another clockwise step again using Eq. 10. This procedure is repeated iteratively until either of the motors has reached some pre-determined position or the motors have reached some maximum alloted time for the extrusion. Once *T*_*counter−clockwise*_ and *T*_*clockwise*_ are computed, plotting of the secondary diagonal is performed as described above.

### 3.3 Calculating goodness of fit for extrusion models

To evaluate the goodness of fit for values of *γ* and *ρ* in the genome-annotation only and RNAP ChIP-seq calibrated models of loop extrusion, we first generated predicted traces of the secondary diagonal using the methods of Section 3.1 (and Section 3.2 for the mixing models). We queried 10^6^ time-points at the interpolation step to ensure that we had sub-kilobase spatial resolution in the predicted extrusion traces. The predicted traces were returned as a list of secondary diagonal positions, i.e. (X,Y)=(*parS* +*x*_*clockwise*_(*t*), *parS* − *x*_*counter−clockwise*_(*t*))), which we then rounded off to 10 kb resolution (to match our 10 kb binning of Hi-C maps). The rounding process resulted in many non-unique pairs of points; we filtered out the list of points to retain solely unique pairs.

Different experimental strains have different lengths of the secondary diagonal; this is due to the fact that the diagonals end roughly where condensin reaches the *ter* site, and will depend directly on the initial position of the *parS* site. As such, we resitricted the analysis and computation of the goodness of fit of each extrusion trace to a fixed number of Hi-C bin “steps”. The number of steps was fixed on a strain by strain basis, and were counted as follows: we used the same procedure outlined above with the model parameters *γ* = 1, *ρ* = 1. i.e. after generating the extrusion trace (sampled at 10^6^ evenly spaced temporal samples from 0 to 50 min), rounding the position to the nearest 10 kb bin, and filtering for unique bins; the total number of counts per strain were set equal to the number of unique (X,Y) pairs.

The goodness of fit value was computed as follows: First, to remove the dependence of contact probability decay with distance away from the main diagonal, we normalized the Hi-C contact matrix by dividing out the expected dependence on distance (i.e. we divided each diagonal parallel to the main diagonal by the mean value for that diagonal). From these “observed over expected” Hi-C maps (*M*_*o/e*_), we obtained the Hi-C map score *M*_*o/e*_(X,Y), and added the Hi-C scores for all values of the unique (X,Y) positions list. We divided the final sum by the number of queried values (i.e. to compute the mean).

This procedure was performed for varying *γ* and *ρ* values. To generate plots, we queried all values of *ρ* from 0.5 to 20 using increments of 0.5; for *γ*, we queried from 0.5 to 10 using increments of 0.5, and from 10 to 100 using increments of 5.

### 3.4 Combining parameter fit surfaces (calculating global optima)

To combine the experimental parameter fit (i.e. goodness of fit) surfaces, we calculated the arithmetic mean of the goodness of fit surfaces. We restricted the analysis to strains with sufficiently long secondary diagonals. This is because some strains (e.g. BWX3381 with a *parS* site at −117°) have a very short secondary diagonal, which resulted in poor goodness of fit surfaces (i.e. large variations in parameter values did not significantly change the overall sum of Hi-C values along the extrusion trace).

## 4 Extrusion model considerations - motivation for a microscopic picture

### 4.1 Inferring condensin’s maximum translocation rate

Previously, the average speed of condensin translocation has been measured *in vivo* in *B. subtilis* (Wang et al., 2017); the average speed of translocation is *v*_*avg*_ ≈ 833 bp/s. The maximum rate of condensin extrusion (*v*_*max*_) based on the genome annotation model is related to the average rate of extrusion (*v*_*avg*_) by:

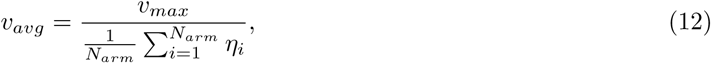

where *N*_*arm*_ is the number of basepairs of a genome arm, *η*_*i*_ is the fold-increase in time it takes condensin to cross any single basepair against the direction of the gene relative to the cooriented direction. The value of *η*_*i*_ is set based on the following rules:

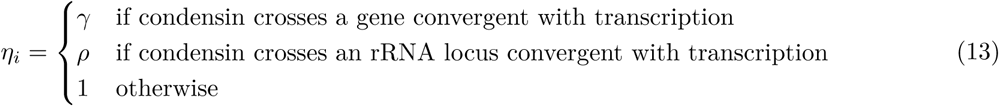

We arrive at this expression as follows. The average rate of extrusion down any single chromosome arm of length *L*_*arm*_ (which has units of length in bp), in terms of real time can be calculated through our genome annotation model up to a constant, *ξ*, which has dimensions of time and converts relative times per basepair (i.e. *η*_*i*_) to real time:

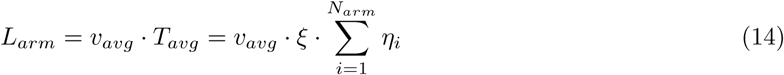

*T*_*avg*_ is the real time it takes for condensin to traverse the distance *L*_*arm*_ at the average speed, *v*_*avg*_. The same distance *L*_*arm*_ can be traversed at the maximum rate of extrusion, *v*_*max*_ (and time *T*_*max*_ according to our assumptions for *η*_*i*_ above), if all the genes are cooriented with transcription (i.e. *η*_*i*_ = 1 for all loci):

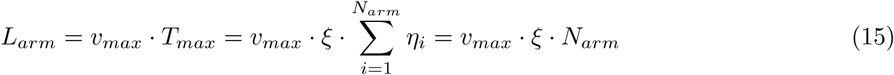

Equating the two relations, we get:

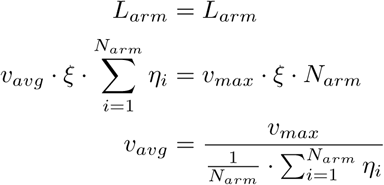

as above. The parameters for *η*_*i*_ that are found to be in the best agreement with Hi-C data are: *γ* ≈ 3 − 5 and *ρ* = 20. Plugging in these numbers into the relation above, using *v*_*avg*_ = 833 bp/s, gives the value of *v*_*max*_ ≈ 1500 *±* 200 bp/s.

### 4.2 Estimating numbers of RNA polymerases per operon

In our experimental growth conditions, the bacteria divide in ~34 mins or with a doubling time of 1.7-1.8 dbl/hr. Using Fig. 2A from Klumpp and Hwa (Klumpp & Hwa, 2008), we obtain a value of ~800-1000 RNAP/cell (for actively transcribing RNA polymerases producing mRNA). Since the cells are in exponential phase, there are roughly 3.1 origins of replication per cell as previously measured (see Graham et al, 2014); thus we estimate there are roughly 2.6-3 copies of the genome per cell on average (i.e. due to multi-fork replication). This corresponds to approximately 266-333/ RNAP/genome copy. Since the genome itself is ~4 Mb long in *B. subtilis* PY79, and ~90% of the genome is covered by operons, then each operon will receive on average (333 *RNAP*)/(4000 *kb* · 0.9) ≈ 0.09 *RNAP/kb*. This estimate of transcribing RNAP numbers is also supported by an independent observation via single-molecule studies (Golding et al., 2005). In their single molecule study, using a candidate gene of length ~4.5 kb, Golding et al. measure that the average times for a transcription burst is about 6 min, and the time between transcription bursts is roughly 37 min. In a given transcription burst, their measured average number of RNA polymerases was 2.2 RNAP/burst. This suggests that the time-average number of RNAP on a gene at any given time is 2.2 · (6 min/(37 min + 6 min))/(4.5 kb) ≈ 0.07 RNAP/kb, which is close to the previously calculated number. Throughout the text use the value of ~0.1 RNAP/kb for the average number density of RNAP at regular genes.

For the case of the rRNA loci, we estimate that the RNAP density is closer to ~5-10 RNAP/kb, which we estimate with two independent ways. First, using Fig. 2A from Klumpp and Hwa (Klumpp & Hwa, 2008), we obtain a value of ~800-1000 RNAP/cell (for transcribing rRNA genes). The calculation follows closely to the one above, where we divide out the numbers of genome copies in our growth conditions, and note that rRNA genes have a total length of ~50 kb in *B. subtilis PY79*. Thus, we estimate that 266-333/ RNAP/genome copy will fall nicely into 50 kb, which results in a density of ≈ 5 − 6 RNAP/kb. Next, we can obtain the relative values of RNAP at rRNA loci using our RNAP ChIP-seq data for wild-type *B. subtilis PY79*. Since the rRNA loci most heavily occupied by RNAP, we can obtain an estimate for the relative fold-enrichment of RNAP at rRNA loci compared to the rest of the genome by taking the median ChIP-seq signal for the top 50,000 RNAP ChIP-seq values, compared to the median ChIP-seq value for the rest of the genome. By our measurements, 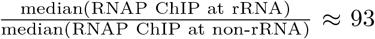. If we set normalize the median ChIP-seq signal at non-rRNA genes such that: median(RNAP ChIP at non-rRNA)≈0.07-0.1 RNAP/kb (see paragraph above), then it follows that the median ChIP-seq signal at rRNA loci corresponds to the range 6.5-9.3 RNAP/kb, which is close to the range of values we measured by the other method. Thus, we use ~5-10 RNAP/kb as our estimate of numbers of RNAPs transcribing rRNA loci in this paper.

## 5 Moving barriers model

### 5.1 Problem statement

Let’s assume that translocating RNA polymerase is an impenetrable barrier to the motion/translocation of the SMC condensin complex. The problem is to figure out the distribution of waiting times, or average time, for condensin to cross a gene body when (A) condensin translocates in the direction of transcription, and (B) when condensin translocates in the direction opposing transcription.

### 5.2 SMC translocation cooriented with transcription (head-to-tail interactions)

In the case of head-to-tail interactions between condensin and RNAP (Fig. 4A, left), a condensin motor subunit translocates at a high speed, *v*_*c*_ =1500 bp/s, on a gene (or operon) until it encounters a transcribing RNAP moving in the same direction at a much lower speed, *v*_*r*_ =80 bp/s. Since RNAP is assumed to be an impermeable barrier, upon RNAP/condensin encounter the condensin slows down its translocation rate to match the RNAP speed, *v*_*r*_, until the end of the gene. Dissociation of RNAP allows the condensin to continue translocating at its original high speed (Fig. 4A, left).

If the condensin meets the RNAP at base position, *x*, then the time for condensin to cross a gene (or operon) is:

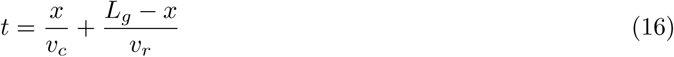

The speed of condensin, *v*_*c*_, and the speed of RNAP, *v*_*r*_, are assumed here to be a constant equal to their average speeds. If the probability of encountering an RNAP at any basepair position within the gene is *p*_*e*_, then the total probability of a condensin encounter with RNAP at (*j* + 1)^*th*^ basepair is *p*_*e*_(1 − *p*_*e*_)^*j*^. We will define the length of the gene, *L*_*g*_ = *N* ⋅ *l*, where *N* is the number of basepairs in the gene, and *l* =1 bp is the unit of length; equivalently, the encounter position (in units of base pairs) *x* = *l* ⋅ *j*, where *j* is a basepair number counter which starts from the beginning of the gene. Together, these relations give the average time, 〈*t*_*cooriented*_〉, to traverse a gene in the head-to-tail case as:

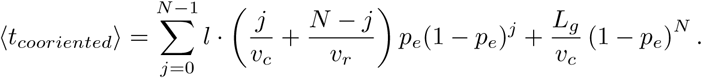

Averages are denoted by 〈*…*〉. After some algebra, the above expression simplifies to:

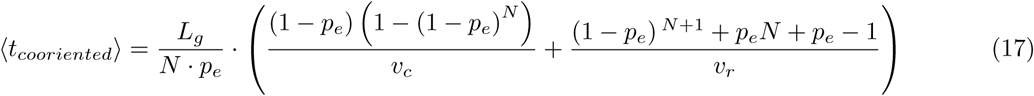

We can make a few simplifying approximations to clean up the expression. First, *N* + 1 ≈ *N* Second, we use the Taylor expansion approximation 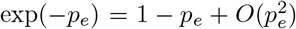, so that, (1 − *p*_*e*_)^*N*^ ≈ exp(−*p*_*e*_ · *N*) = exp(−〈*R*〉), where we define 〈*R*〉 = *N* ⋅ *p*_*e*_ to be the average number of RNA polymerases on the gene. Together, these approximations give:

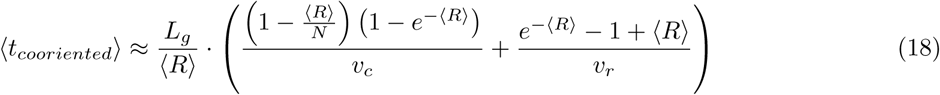

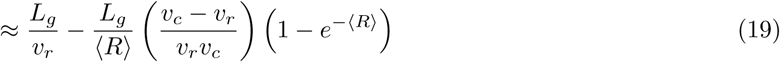

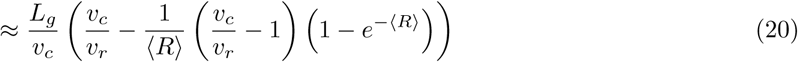

The value of 〈*t*_*cooriented*_〉 (from the unsimplified equation, Eq. 17) is plotted as a function of gene length *L*_*g*_, and RNA polymerase density per kilobase (〈*R*〉/1000*kb*) (Fig. 4B, left); we have assumed that *v*_*c*_ = 1500 *bp/s* and *v*_*r*_ = 100 *bp/s* for the calculation. The value of *p*_*e*_ is simply the RNA polymerase density on the gene (per base-pair). Interestingly, only two parameters (the average number of RNA polymerases for that gene) and the gene length are required to calculate the time to cross the gene. Most simply, the gene length only changes the time linearly, whereas the numbers of RNAP molecules on the gene increases the time more non-trivially.

To get a better intuition for how strongly transcription affects condensin translocation rates, we plug in some real values: For a typical operon of length 3 kb in a protein-coding locus, the average RNAP density is ~0.1 RNAP/kb (or an average of 0.3 RNAP for that operon), the fold-increase in time to cross the operon (compared to the case where the condensin is moves at its maximum rate *v*_*c*_) (i.e 〈*t*_*cooriented*_〉*v*_*c*_/*L*_*g*_) is 2.2-fold longer, suggesting that transcription does not strongly slow down the process of condensin extrusion in the head-to-tail case. For an rRNA locus, a 3 kb operon with 5 RNAP/kb, it takes 8.2 times longer to cross the locus compared to the “free extrusion” scenario.

### 5.3 SMC translocation opposing transcription (head-to-head interactions)

In the case of head-to-head collisions (Fig. 4A, right), condensin translocates across a gene (or operon) at its native high speed, *v*_*c*_, towards the transcription start site until it meets an RNAP. Upon encountering RNAP, condensin translocation towards the transcription start site is stopped; condensin is then pushed back to the transcription termination site by the transcribing RNAP at the speed of the RNAP transcription, *v*_*r*_. Once RNAP dissociates (i.e. when it reaches the end of the gene), condensin can attempt to cross the gene again; the condensin will only successfully cross the gene if no RNAPs are encountered during its run through the gene (Fig. 4A, right).

We assume that a translocating RNAP forms a completely impermeable barrier to condensin. The time for condensin to cross a gene of length, *L*_*g*_, is the sum of times for unsuccessful traversal attempts plus the time for a successful traversal.

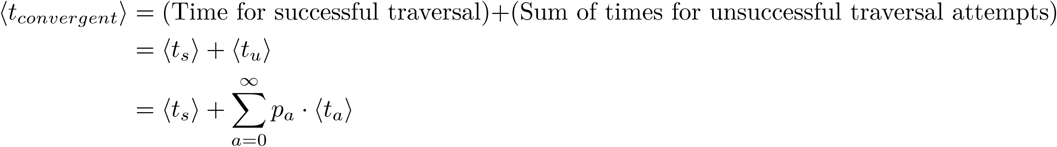

where *a* is the number of unsuccessful attempts, *p*_*a*_ is the probability of observing *a* unsuccessful attempts, *t*_*a*_ is the total time for *a* unsuccessful traversal attempts and *t*_*s*_ is the total time for a successful attempt. The sucessful traversal time is a constant:

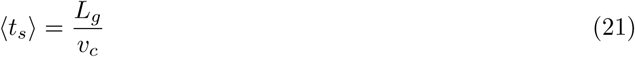

The number of attempts, *A*, that condensin will make before successfully traversing the gene (in the direction opposing transcription) is given by a Geometric distribution with the probability of encountering an RNAP at any position in the gene of 1 − (1 − *p*_*e*_)^*N*^:

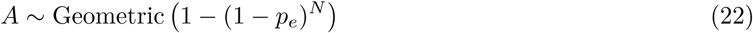

As before, *N* is the number of basepairs in the gene of length *L*_*g*_. Since *A* is geometrically distributed, the probability of *a* unsucessful attempts to cross the gene followed by a successful attempt on the (*a* + 1)th attempt is then: *p* = (1 − (1 − *p*_*e*_)^*N*^)^*a*^(1 − *p*_*e*_)^*N*^. Thus, the average time for unsuccessful traversal attempts is given by:

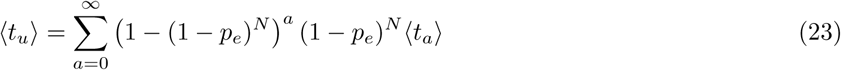

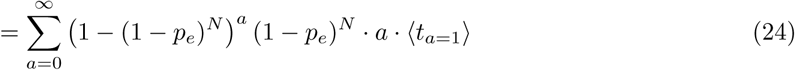

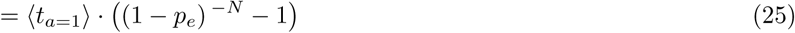

The second line follows by noting that 〈*t*_*a*_〉 = *a* ⋅ 〈*t*_*a*=1_〉 since the mean of the sum of *a* independently and identically distributed random variables is equal to sum of the means for any one variable (or *a* times the mean of one of the attempts).

We now seek to determine the form of 〈*t*_*a*=1_〉. The time of a single unsuccessful traversal attempt has duration

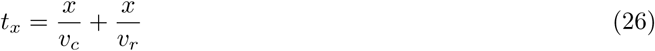

assuming that a condensin meets RNAP at nucleotide position, *x*, in the gene. Unlike in Section 5.2, Eq. 16, where the probability of condensin encountering an RNAP was given by a simple Geometric distribution (and hence it could pass through the gene with finite probability of not encountering RNAP), in this case, we stipulate that condensin must encounter an RNAP in the gene (i.e. since the probability is conditioned to be an unsuccessful traversal attempt). Thus, *x* (where *x* = *j* ⋅ *l*, *L*_*g*_ = *N* ⋅ *l*, *l* =1 bp) is given by a Truncated Geometric distribution,

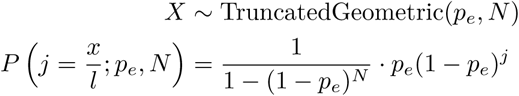

which has support *xϵ*[0, *L*_*g*_ − 1]. So, the average time for condensin to encounter an RNAP in a single attempt is related to the first moment 〈*x*〉

Thus,

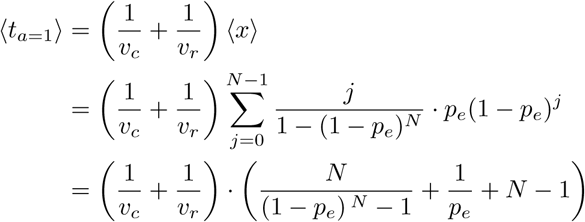

After some calculations, Eq. 23 becomes:

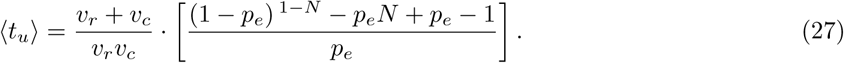

Putting this together, we finally arrive at the final exact gene-crossing time (using Eqs. 21 and 27):

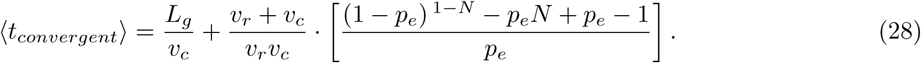

With some approximations as in the previous section (i.e. *e*^−*p*_*e*_^ ≈ 1 − *p*_*e*_, *N* − 1 *≈ N*, *p*_*e*_ ≪ 1, and setting 〈*R*〉 ≈ *p*_*e*_*N*), this expression becomes:

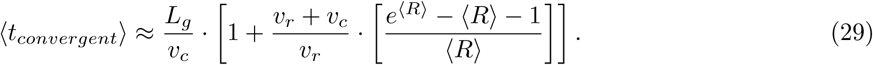

We can plot the average times for traversal in the convergent orientation (Eq.28) as shown in Fig. 4B, on the right. Interestingly, we see that in convergent motion, average times quickly escape physiologically possible conditions as a function of RNAP density. The physiological limit we defined as the 35 minute time mark, which is the experimentally observed time to fully “zip” a chromosome arm; in reality, the limit for any single genetic feature is smaller than this value. With long loci, like ribosomal RNA loci where transcription units are close to 10 kbp long, with as few as 0.6-0.8 RNA polymerases per kilobase, we would predict that the locus would become impenetrable to condensin.

For numerical comparison with the head-to-tail interaction case we calculate 〈*t*_*convergent*〉_*v*_*c*_/*L*_*g*_ (i.e. the fold-increase in time compared to the “no RNAP”/ “free-extrusion” scenario). For a typical operon of length 3 kb in a protein-coding locus, the average RNAP density is ~0.1 RNAP/kb (or an average of 0.3 RNAP for that operon), the fold-increase in time is 2.8-fold (compared to 2.2 for the head-to-tail case). This suggests that for most regularly transcribed genes, there is no strong difference in time to cross the locus in the head-to-head versus head-to-tail cases, consistent with our current results and previous results (Tran et al., 2018). In contrast, for an rRNA locus, a 3 kb operon with 5 RNAP/kb, it takes 314-fold longer to cross the locus compared to the free extrusion scenario. This 314-fold slow-down in the head-to-head scenario is in stark contrast to the 8.2-fold slow-down for the head-to-tail case. Thus, by increasing the RNAP density, there can be huge temporal penalties for crossing a locus in the head-to-head case with impermeable barriers.

### 5.2 Estimating the value of *ρ*, and why the optimal value varies from strain to strain

We can relate the value of *ρ* that we used in the *gene position and direction model* (Fig. 2A) to the moving barriers model derived above. The probability of crossing a gene in the “head-to-head” orientation is *f*_*conv*_ and in the “head-to-tail” orientation is *f*_*coord*_, and the probability of crossing a locus without a gene is *f*_*free*_; *f*_*free*_ + *f*_*conv*_ + *f*_*coord*_ = 1. In the *B. subtilis* genome, *f*_*free*_ < 0.1. Then, if one of the bidirectional extrusion motors (labelled *A*) is crossing a gene at genome position, *x*_*A*_, and the other motor (labelled *B*) is crossing a different locus at position, *x*_*B*_, then the instantaneous value of *ρ* is

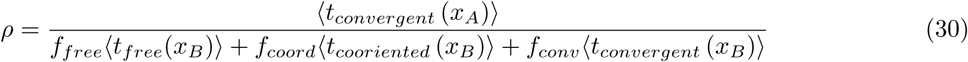

where 〈*t*_*free*_(*x*)〉 is the time to cross a locus without impediments (i.e. at the condensin motor’s maximal speed). We note that it is important to use consistent “locus lengths” (e.g. the time for 1 bp, or 100 bp) to calculate each of the average quantities to obtain good estimates of *ρ*.

Since, as we saw above, 〈*t*_*convergent*_〉 and 〈*t*_*cooriented*_〉 are functions of both gene length and RNAP density (i.e. encounter probability) for the locus pair *x*_*A*_ and *x*_*B*_, the parameter *ρ* that we measure from experimental fits will actually be an average quantity; the average is the weighted average over all combinations of gene lengths and RNAP densities encountered by the pair of *A* and *B* motors as extrusion occurs away from the *parS* loading site. This observation, interestingly, provides a rationale for why there are some variations in the optimal values of *ρ* obtained from strain to strain: as we move the *parS* site, we slightly shift the set of gene combinations (i.e. the distribution of lengths for *t*_*convergent*_ and *t*_*cooriented*_) encountered by the extruding motor along its path, which will result in different combinations of optimal *ρ*.

As a baseline minimum estimate of *ρ* from the moving barriers model (Fig. 4B), we note that since it measures the effect of motor *A* at an rRNA locus (i.e. crossing an ~10 kb operon with an RNAP density of 5-10 RNAP/kb, whereas motor *B* is crossing at a regular operon (~ 1-3 kb in length), with a time average density of ~0.05-0.1 RNAP/kb it follows that,

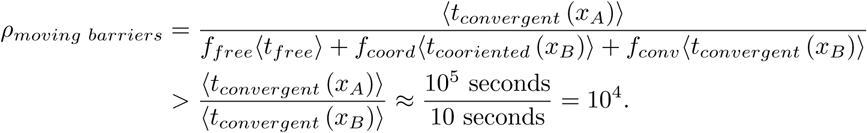

since 〈*t*_*free*_〉 < 〈*t*_*coordinated*_〉 < 〈*t*_*convergent*_〉.

## 6. Permeable moving barriers model

### 6.1 Problem statement

Let’s assume that RNAP is a partially permeable barrier to the motion/translocation of the SMC condensin complex. Our problem is to determine the average time that SMC spends at a locus position *x* within a gene body due to interactions with RNAP. Assuming that the amount of time that an SMC spends at that locus is proportional to its ChIP-seq density profile, we ultimately seek to predict the SMC ChIP-seq enrichment profile as a function of RNAP density (i.e. RNAP ChIP-seq). In addition, we will derive some quantitative intuition for a potential molecular mechanism behind the slowing down of SMC when travelling head-on against genes versus in the gene direction. We will consider below the two separate cases for the SMC ChIP-seq: (A) SMC extrusion in the direction of transcription, and (B) SMC extrusion in the direction opposing transcription. However, before analytically estimating the shapes of the SMC ChIP-seq profiles, we will obtain estimates for the value of the rate at which condensin bypasses RNAP.

### 6.2 First passage times, and estimating the permeability rate *µ*

The permeable moving barriers model is similar to other types of dynamical systems which have over the years garnered many different names such as: “Asymmetric Persistent Random Walks”, “Markov Jump Processes”, among others (Codling et al., 2008). The permeable moving barriers model can be mapped to the Telegrapher’s equation, and the problem of dynamic instability of microtubules (Bicout et al., 1997).

To obtain an estimate of *µ* (the rate of bypassing RNAP) we turn to the calculation of the effective speed of a condensin as it traverses a locus (e.g. the rRNA genes) in the head-to-tail direction. The effective speed of traversing a gene, *v*_*eff*_, is given by:

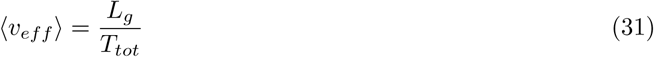

where *T*_*tot*_ is the total time to cross a locus of length *L*_*g*_. From Hi-C data (i.e. the length of the secondary diagonal trajectory as condensin passes the rRNA locus), we estimate that to cross 30 kb rRNA locus in the head-to-head orientation it takes about ~8-15 minutes, therefore, *L*_*g*_ = 30,000 bp, *T*_*tot*_ ≈ 480− 900 s, or *v*_*eff*_ ≈ 30 − 60 bp/s.

Analogously, the problem of traversing the total length *L*_*g*_ can be broken into segments of “steps forward” and “steps backward”. The average distance (the mean freepath) that a condensin will travel “forward” before encountering an RNAP is found as follows: We assume the condensin travels a distance *d* to the next RNAP in time *t* with speed *v*_*c*_. The length of the gap, *d* (in the operon’s frame of reference) between the condensin and RNAP shrinks from its initial (average) value of 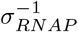 (the average distance between RNAPs) as RNAP moves towards the condensin at speed *v*_*r*_. Thus, 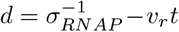 and *d* = *v*_*c*_*t*. These two equations can be solved for *t* and *d* resulting in 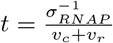, and 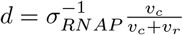. The average distance that condensin will travel “backwards” (pushed by and RNAP it encounters) is proportional to the RNAP speed and the mean bypassing time (i.e. inverse of the permeability rate): *v*_*r*_ ⋅ *µ*^−1^. Thus, the operon/gene length *L*_*g*_ can be composed of the sum of *n* increments of length *δL*_*i*_ (where *iϵ*[1, *n*]), where the average length is 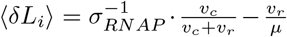. Similarly to the distance increments, we can define temporal increments for each of the combined “forwards” followed by “backwards” steps as *δT*_*i*_. The average 〈*δT*_*i*_〉 is readily calculated: In the forward direction (before meeting an RNAP), condensin crosses the gap between the previous RNAP (or the TSS) in an average time equal to 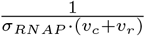 (as calculated above). In the backwards direction, it spends the mean bypassing time, which is equal to the permeability rate via 1*/µ*. Thus, mean time per segment is: 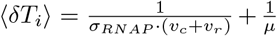. The effective condensin speed can be approximated (to zeroth order) as:

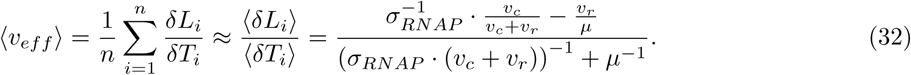

We can solve this equation for *µ*:

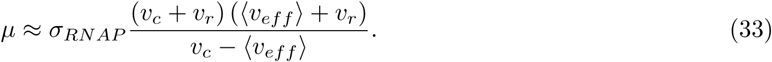

For the values of condensin speed, *v*_*c*_ ≈ 800 bp/s, RNAP speed *v*_*r*_ = 80 bp/s, 〈*v*_*eff*_〉 ≈ 30 − 60 bp/s, *σ*_*RNAP*_ ≈ 5 − 10 RNAP/kb, we obtain an estimate of *µ* ≈ 0.6 − 1.7 *s*^−1^, which is close to the values that we obtain via simulations (see below).

### 6.3 ChIP-seq profiles: SMC translocation cooriented with transcription (head-to-tail interactions)

We next move to estimating the SMC ChIP-seq enrichment profiles analytically. We start with the case of head-to-tail interactions. The amount of time an SMC spends at any position *x* within a gene body will be proportional to the number of times it crosses the locus weighted by the speed of crossing it. Without loss of generality, let’s define *χ*_*c*_ = *χ*_*c*_(*x*) as the probability that an SMC crosses locus *x* with the RNAP speed, *v*_*r*_ = *v*_*r*_(*x*) (which can be a position-dependent function). In other words, *χ*_*c*_(*x*) is the probability that SMC is “captured”/blocked by an RNAP at *x* or upstream of *x* (i.e. before *x*, towards the TSS) and it passes *x* moving towards the TES by a speed limited by the local transcription elongation rate. The value (1 − *χ*_*c*_(*x*)) is then the probability that SMC crosses locus *x* with its native speed, *v*_*c*_, which we will assume here is a constant in the absence of interactions with RNAP. The average amount of time an SMC spends at locus *x* is then given by the probability density *σ*_*SMC−cooriented*_(*x*):

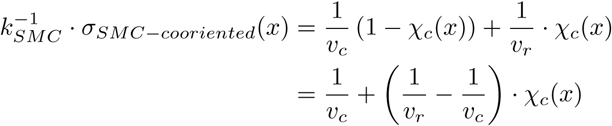

where *k*_*SMC*_ is the rate at which an SMC arrives at the TSS. The observed SMC ChIP-seq enrichment is 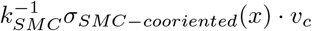,

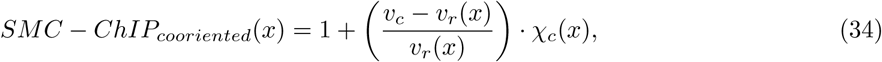

which has the interpretation that the ChIP-seq signal is enriched by a factor proportional to the difference of SMC to local RNAP speeds. Moreover, the ChIP-seq enrichment has amplitude proportional to the probability, *χ*_*c*_, that SMC is “captured” or “blocked” by the RNAP upstream of position *x*. We will discuss below the general form that *χ*_*c*_(*x*) can take, and consider specific example cases. For now, we simply note that *χ*_*c*_(*x*) will depend on the density of RNA polymerase (*σ*_*RNAP*_(*x*)), and can be related to a permeability rate, *µ*, which describes how likely it is for RNAP to block the SMC extrusion.

Interestly, and of note, we see that the ChIP-seq signal will depend on the position-dependent RNAP rate *v*_*r*_(*x*). Indeed, since the rate of RNAP transcription within the gene will determine the enrichment of the RNAP, it follows that the SMC ChIP-seq signal will be strongly (positively) correlated with the RNAP signal. For instance, in a gene body, wherever the RNAP ChIP-seq value increases (i.e. where *v*_*r*_(*x*) is lowest), we also expect that the SMC ChIP-seq value will increase, even while keeping *χ*_*c*_ constant.

### 6.4 ChIP-seq profiles: SMC translocation opposing transcription (head-to-head interactions)

For the case of SMC motion opposing transcription, we break up the problem into two contributions: one contribution is due to SMC moving unhindered by RNAP towards the TSS (Fig. 4A, row i), and another contribution from SMC moving towards the TES being pushed back by RNAP (Fig. 4A, rows ii-iii). For this derivation, we use the shorthand *χ*_*c*_ = *χ*_*c*_(*x*). Again, we define *χ*_*c*_(*x*) as the probability that an SMC is “captured”/blocked by an RNAP somewhere between the position *x* and the TSS and then crosses the locus *x* with the RNAP speed on its way back through *x* (since it is being pushed by RNAP); we assume, of course, that the SMC has already passed position *x* at least once at its native speed, *v*_*c*_, moving towards the TSS from the TES direction.

The probability that the SMC passes a locus *x* only once is the probability that it traverses the gene and “escapes” without being captured and brought back; it is equivalent to: (1 − *χ*_*c*_). The probability that SMC passes locus *x* twice is the probability it fails to exit the gene (i.e. is captured and brought back) times the probability it successfully escapes capture: *χ*_*c*_(1 − *χ*_*c*_). The probability that it passes three times is 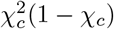, and so forth. Thus, the amount of time SMC spends at locus *x* is given by the density *σ*_*SMC−convergent*_(*x*):

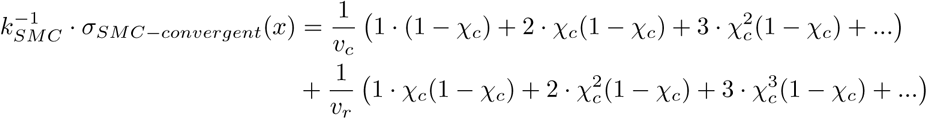

The terms involving *v*_*c*_ are understood to be the motion towards the TSS (unhindered by RNAP), and the terms involving *v*_*r*_ are understood as motion towards the TES as SMC is pushed back by RNAP at the local average RNAP speed (*v*_*r*_ = *v*_*r*_(*x*)). Collecting these terms, and after some straight-forward algebra, we get:

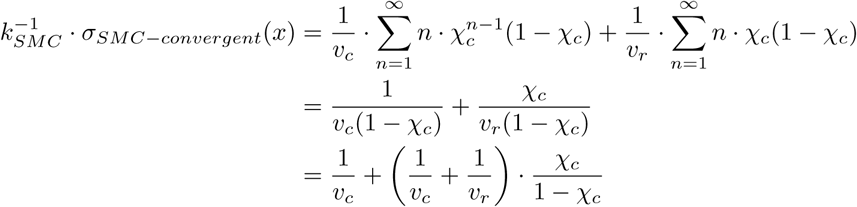

The observed SMC ChIP-seq enrichment is 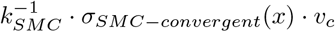,

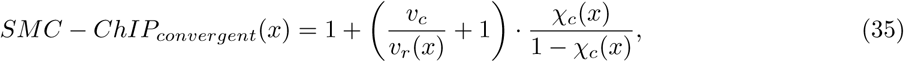

The final result for the SMC ChIP-seq enrichment within a gene can be interpreted as follows: it is the sum of the relative times to cross *x* in both the forward and reverse directions 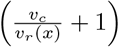 (i.e. the average time spent at position *x* per crossing attempt), multiplied by the average number of attempts to cross the locus, 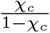. The latter term is formally equivalent to the expectation value of a negative binomial distribution.

### 6.5 SMC ChIP-seq profiles: comparing head-to-head and head-to-tail encoutners

Right away (following the two sections above), we see that within a gene body, we expect there to be a stronger enrichment of SMC in the head-to-head interaction case than the head-to-tail case:

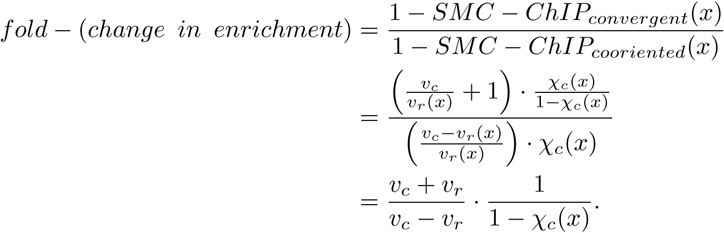

Thus, part of the observed change in enrichment will come from a relative difference in densities (i.e. 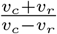) as condensin meets RNAP more frequently when it translocates opposite to transcription than with transcription. However, we expect that this value is relatively small (<1.3, even plugging in generous values for the condensin speed (*v*_*c*_ = 800 bp/s), and RNAP speeds (*v*_*r*_ = 80 bp/s)). Thus, the observed 1.6 to 2-fold change in SMC enrichment seen experimentally (i.e. *fold* − (*change in enrichment*) ≈ 2 for a 0.1-1 kb operon (Fig. S11B, left), and ≈ 1.6 for a 1-2 kb operon (Fig. S11B, right)) between the head-to-head and head-to-tail cases cannot be accounted for just based on the differences in condensin speed. Indeed, experimental values suggest that:

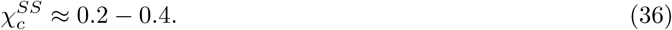

The capture probability, *χ*_*c*_(*x*), can be linked to the permeability rate as we will show below.

### 6.6 General forms for the capture probability, *χ*_*c*_(*x*) and the permeability rate, *µ*

To calculate *χ*_*c*_(*x*), we must consider separately the cases of head-to-tail and head-to-head encounters between the translocating SMC motor and the RNAP. The value *χ*_*c*_(*x*) will depend on the probability density of RNAP in a given operon, at a given position, *x*, the length of the operon (i.e. the position *x*), and the permeability rate, *µ*. We can compute *χ*_*c*_(*x*) using a Master equation, and we consider only the case of head-to-tail encounters for simplicity. To write the Master equation (i.e. the time-evolution equation of the state *χ*(*x*)), we consider and transitions between the two possible states. That is: (1) the state *χ*_*c*_(*x*) where condensin is hindered by RNAP at position *x*, and (2) the state (1-*χ*_*c*_(*x*)) where condensin is unhindered by RNAP at *x*. The transition rates between each state are illustrated in the figure below.

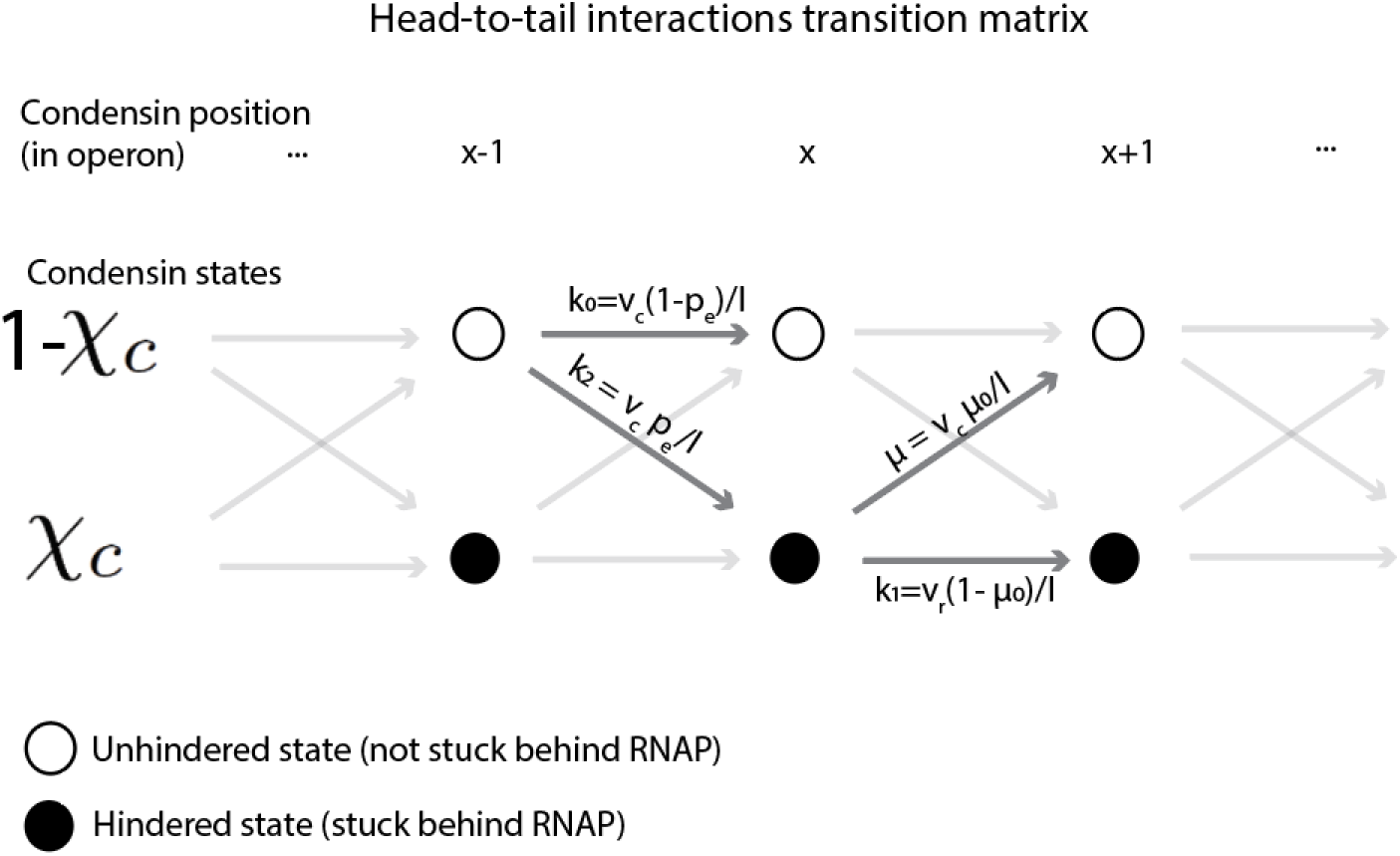

We arrive at these transition rates by considering 4 requirements. The first two requirements are for the transitions out of the state *χ*_*c*_(*x*).

#### Requirement 1

The rate at which condensin bypasses an RNAP (i.e. transitions from *χ*_*c*_(*x*) → 1 − *χ*_c_(*x*+ 1)) is: *µ* = *v*_*c*_ ⋅ *µ*_0_/*l*. We define the permeability “probability” as *µ*_0_, so the permeability probability per basepair is *µ*_0_*/l* (where *l* = 1 bp, as before) and *µ* is the value we ultimately aim to find from experiments. For simplicity, we consider only the case of constant *µ*_0_, but in its most general form the permeability probability could be a function of the position within an operon (i.e. *µ*_0_ = *µ*_0_(*x*)). The form for the permeability/bypass rate (*µ* = *v*_*c*_ ⋅ *µ*_0_*/l*) makes sense since it stipulates that the upper bound on condensin’s rate of escape is its speed, *v*_*c*_ (i.e. when *µ*_0_ = 1).

#### Requirement 2

When a condensin moves from a hindered state at one position (e.g. *x*) to a hindered state at the next position (e.g. *x* + 1) the rate is limited by the RNAP speed (i.e. *v*_*r*_/*l*), and will also depend on condensin not “escaping” the bound state (i.e. 1 − *µ*_0_). Together, this requires the transition *χ*_*c*_(*x*) → *χ*_*c*_(*x* + 1) to have a rate *k*_1_ = *v*_*r*_(1 − *µ*_0_)*/l*. The limiting cases make sense: when *µ*_0_ = 0 (i.e. when RNAP is totally impermeable), then the transition rate becomes the RNAP speed (*v*_*r*_/*l*) and the permeability rate becomes:*µ* = 0 *s*^−1^ (i.e. condensin cannot bypass RNAP). On the other hand, when *µ*_0_ = 1, condensin can only step out of *χ*_*c*_(*x*) by going back to the “unhindered” state, and it does so at the rate *µ* = *v*_*c*_/*l*.

The transitions out of the state 1 − *χ*_*c*_(*x −* 1) lead to the remaining 2 requirements.

#### Requirement 3

the total rate for condensin to move out of state 1 − *χ*_*c*_(*x*) is *v*_*c*_. This requirement stipulates that the transition 1 − *χ*_*c*_(*x* − 1) → 1 − *χ*_*c*_(*x*) takes place with a rate *k*_0_ = *β* ⋅ *v*_*c*_, where *β* remains to be determined, but 0 ≤ *β* ≤ 1. The factor *β* is necessary since the transition 1 − *χ*_*c*_(*x* − 1) → *χ*_*c*_(*x*) is also possible. To find the value of *β*, we have one last requirement.

#### Requirement 4

When an unhindered condensin takes a step, it has a probability *p*_*e*_ of encountering an RNAP at *x*. This requirement stipulates that the transition 1 − *χ*_*c*_(*x* − 1) → *χ*_*c*_(*x*) should occur with a rate *k*_2_. The probability, *p*_*e*_ is related to *k*_2_ and *k*_0_ (i.e. *β ⋅ v*_*c*_) via 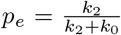. Thus, 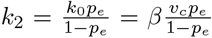. We can solve for *β* using the third requirement, which stipultes that: *k*_2_ + *k*_0_ = *v*_*c*_. After some substitutions, this results in *β* = 1 − *p*_*e*_. Thus, we arrive at the rates shown in the diagram (i.e. the condensin steps from unhindered state to unhindered state with rate *k*_0_ = *v*_*c*_(1 − *p*_*e*_) and from unhindered state to hindered state with rate *k*_2_ = *v*_*c*_*p*_*e*_).

We can now write down the Master equation for the time-evolution of the state, *χ*(*x*).

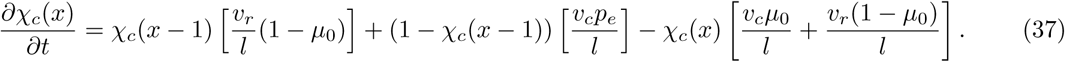

At the steady-state, 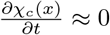. Moreover, for sufficiently large values of *x*, there is also a “spatial” steady-state (i.e. when 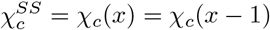. In these two conditions we obtain:

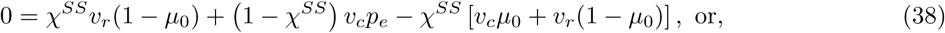

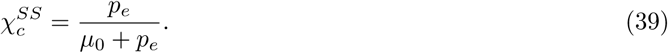

Finally, solving for *µ*_0_ and recalling that *µ* = *µ*_0_*v*_*c*_, we get:

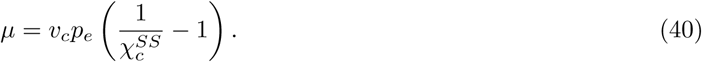

We can obtain *µ* from 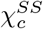 (estimated to be ~0.2-0.4 in the previous section), by noting that the average density of RNAP at non-rRNA is 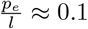 RNAP/kb (see Section 4.2). Thus,

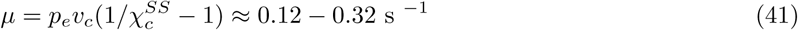

This estimate suggests that it takes up to ~8 seconds to bypass an RNAP transcribing a protein-coding gene.

### 6.7 Simulating SMC ChIP-seq and locus crossing times

We carried out simulations of the permeable moving barriers model to measure SMC occupancy as a function of distance within an operon and to calculate estimates of the locus crossing times. The simulations were performed on a discrete 1D lattice, with explicit RNAP and condensins. The RNAPs were mutually exclusive, and not allowed to occupy the same lattice site; similarly, condensins (if multiple were on the same operon) were mutually exclusive and could not occupy the same lattice site. Each discrete lattice site corresponded to 1 bp. Operons were simulated in the range of lengths 1 kb to 10 kb (or 1000 to 10,000 lattice sites), plus 100 lattice sites upstream and downstream of the operon. Simulations proceeded with a fixed time step equal to a condensin forward step; in this way, condensins were deterministic walkers, occupying sequential 1 bp lattice sites at a time, and the simulation time step was equal to the inverse of the condensin speed (assumed 800 bp/s, or 1.25 ms per simulation lattice site). RNAP forward steps were taken with a probability *p* = 0.1 simulation steps, corresponding to an average of 80 bp/s. RNAP were randomly initiated with exponential kinetics at the transcription start site with a probability *k*_*r*_ = 10^−3^ to 10^−5^, which gave rise to average numbers of RNAP per kilobase in the range of 0.1 to 10. RNAPs reaching the end of the operon (i.e. position 10,100 for a 10 kb operon) were dissociated immediately. Similarly to RNAPs, condensins were randomly initiated with exponential rates corresponding to probability 10^−4^ per simulation time step, this corresponds roughly to 1 SMC passing through a TSS (for the first time) every 12.5 seconds or a separation of roughly ~20 kb per SMC. The starting position of SMCs, however, was always at lattice position “0”. SMCs reaching the end of the simulation box (e.g. position 10,200 for a 10 kb operon) were dissociated. We estimate this is a ~ 10-fold overestimate of the density of SMCs, but we note (see below) that for the purposes of simulating SMC ChIP-seq profiles, it does not significantly change the shapes of the observed curves.

The simulation step rules proceeded as follows. If an RNAP took a tentative forward step and encountered another RNAP at the next lattice site, the step was aborted (i.e. due to the mutual exclusion). If a condensin moved a forward step and it encountered another condensin, the step was similarly aborted (due to mutual exclusion). However, if condensin took a tentative forward step and it encountered an RNAP, it took the step with probability *µ* (corresponding to the rate of bypassing RNAP). Only in this situtation were two molecules allowed to occupy the same lattice site. The typical simulation was run for 10,000,000 steps. In each simulation step, all particle positions were updated according to the rules above, and the average time to cross from the transcription start site to end site were computed.

To compute the SMC enrichment profiles, we created an array of positions (equal to the length of the simulation box) and added 1 count for each unique cluster of SMCs at each time point sampled at the position closest to the RNAP. Sampling was done once per simulation time step. Unique clusters were defined as sets of contiguous lattice sites occupied by successive SMCs. For instance, consider SMC at lattice positions in the set {*i=15,100,101,102,5056,5057,5090*}. This hypothetical scenario would constitute 4 clusters of SMCs (a single SMC at position 15, a cluster of 3 SMCs registered at position 102, a cluster of 2 SMCs registered at position 5057, and a single SMC at the position 5090. Thus, similarly to ChIP-seq, which would not distinguish between clusters of adjacent SMCs (and thus does not strictly speaking measure SMC enrichment), we account for the potential confounding factor of SMCs forming a consecutive train in our simulations. We note, however, that for the experimentally relevant densities of RNAP, permeability rates, and even our overestimate of the condensin density, the average number of SMCs per operon were typically less than 2. Thus, clustering of SMCs was minimal, and this justifies using the larger SMC density (i.e. to obtain more occurrences of operon traversal). SMC enrichment was performed by dividing the counts per lattice site by the mean of the first 100 lattice sites in the simulation box (i.e. the sites before the operon).

## References

1. Hirano T (2016) Condensin-based chromosome organization from bacteria to vertebrates. Cell 164(5):847–857.

2. Alipour E, Marko JF (2012) Self-organization of domain structures by DNA-loop-extruding enzymes. Nucleic Acids Res 40(22):11202–11212.

3. Fudenberg G, et al. (2016) Formation of chromosomal domains by loop extrusion. Cell Rep 15(9):2038–2049.

4. Sanborn AL, et al. (2015) Chromatin extrusion explains key features of loop and domain formation in wild-type and engineered genomes. Proc Natl Acad Sci 112(47):E6456–E6465.

5. Riggs AD (1990) DNA methylation and late replication probably aid cell memory, and type I DNA reeling could aid chromosome folding and enhancer function. Philos Trans R Soc Lond B Biol Sci 326(1235):285–297.

6. Naysmith K (2001) Disseminating the genome: joining, resolving, and separating sister chromatids during mitosis and meiosis. Annu Rev Genet 35:673–745.

7. Wang X, et al. (2015) Condensin promotes the juxtaposition of DNA flanking its loading site in Bacillus subtilis. Genes Dev 29(15):1661–1675.

8. Wang X, Brandão HB, Le TBK, Laub MT, Rudner DZ (2017) Bacillus subtilis SMC complexes juxtapose chromosomal arms as they travel from origin to terminus. Science 355(6324):524–527.

9. Terakawa T, et al. (2017) The condensin complex is a mechanochemical motor that translocates along DNA. Science 358(6363):672–676.

10. Ganji M, et al. (2018) Real-time imaging of DNA loop extrusion by condensin. Science 360(6384):102–105.

11. Eeftens JM, et al. (2017) Real-time detection of condensin-driven DNA compaction reveals a multistep binding mechanism. EMBO J 36(23):3448–3457.

12. Keenholtz RA, et al. (2017) Oligomerization and ATP stimulate condensin-mediated DNA compaction. Sci Rep 7(1):14279.

13. Kim H, Loparo JJ (2016) Multistep assembly of DNA condensation clusters by SMC. Nat Commun 7:10200.

14. Marko JF, Rios PDL, Barducci A, Gruber S (2018) DNA-segment-capture model for loop extrusion by structural maintenance of chromosome (SMC) protein complexes. bioRxiv 325373. doi:10.1101/325373.

15. Fudenberg G, Abdennur N, Imakaev M, Goloborodko A, Mirny LA (2017) Emerging evidence of chromosome folding by loop extrusion. Cold Spring Harb Symp Quant Biol 82:45–55.

16. Gassler J, et al. (2017) A mechanism of cohesin-dependent loop extrusion organizes zygotic genome architecture. EMBO J 36(24):3600–3618.

17. Haarhuis JHI, et al. (2017) The cohesin release factor WAPL restricts chromatin loop extension. Cell 169(4):693–707.

18. Rao SSP, et al. (2017) Cohesin loss eliminates all loop domains. Cell 171(2):305–320.

19. Schwarzer W, et al. (2017) Two independent modes of chromatin organization revealed by cohesin removal. Nature 551(7678):51–56.

20. Wutz G, et al. (2017) Topologically associating domains and chromatin loops depend on cohesin and are regulated by CTCF, WAPL, and PDS5 proteins. EMBO J 36(24):3573–3599.

21. Marbouty M, et al. (2015) Condensin- and replication-mediated bacterial chromosome folding and origin condensation revealed by Hi-C and super-resolution imaging. Mol Cell 59(4):588–602.

22. Le TBK, Imakaev M V., Mirny LA, Laub MT (2013) High-resolution mapping of the spatial organization of a bacterial chromosome. Science 342(6159):731–734.

23. Gruber S, Errington J (2009) Recruitment of condensin to replication origin regions by ParB/SpoOJ promotes chromosome segregation in B. subtilis. Cell 137:685–696.

24. Sullivan NL, Marquis KA, Rudner DZ (2009) Recruitment of SMC by ParB-parS organizes the origin region and promotes efficient chromosome segregation. Cell 137(4):697–707.

25. Wilhelm L, et al. (2015) SMC condensin entraps chromosomal DNA by an ATP hydrolysis dependent loading mechanism in Bacillus subtilis. Elife 4:e06659.

26. Minnen A, et al. (2016) Control of Smc coiled coil architecture by the ATPase heads facilitates targeting to chromosomal ParB/parS and release onto flanking DNA. Cell Rep 14(8):2003–2016.

27. Marbouty M, et al. (2014) Metagenomic chromosome conformation capture (meta3C) unveils the diversity of chromosome organization in microorganisms. Elife 3:e03318.

28. Tran NT, Laub MT, Le TBK (2017) SMC Progressively Aligns Chromosomal Arms in Caulobacter crescentus but Is Antagonized by Convergent Transcription. Cell Rep 20(9):2057–2071.

29. Wang X, et al. (2018) In vivo evidence for ATPase-dependent DNA translocation by the Bacillus subtilis SMC condensin complex. Mol Cell 6(71):841–847.

30. Miermans CA, Broedersz CP (2018) Bacterial chromosome organization by collective dynamics of SMC condensins. J R Soc Interface 15(147). doi:10.1098/rsif.2018.0495.

31. Le TBK, Laub MT (2016) Transcription rate and transcript length drive formation of chromosomal interaction domain boundaries. EMBO J 35(14):1582–1595.

32. Heinz S, et al. (2018) Transcription elongation can affect genome 3D structure. Cell 174(6):1522–1536.

33. Rowley MJ, et al. (2017) Evolutionarily conserved principles predict 3D chromatin organization. Mol Cell 67(5):837–852.

34. Uhlmann F (2016) SMC complexes: From DNA to chromosomes. Nat Rev Mol Cell Biol 17(7):399–412.

35. Busslinger GA, et al. (2017) Cohesin is positioned in mammalian genomes by transcription, CTCF and Wapl. Nature 544(7651):503–507.

36. Rowley MJ, et al. (2019) Condensin II Counteracts Cohesin and RNA Polymerase II in the Establishment of 3D Chromatin Organization. Cell Rep 26(11):2890–2903.

37. Merrikh H, Zhang Y, Grossman AD, Wang JD (2012) Replication–transcription conflicts in bacteria. Nat Rev Microbiol 10:449–458.

38. Touzain F, Petit M-A, Schbath S, El Karoui M (2011) DNA motifs that sculpt the bacterial chromosome. Nat Rev Microbiol 9(1):15–26.

39. Badrinarayanan A, Reyes-Lamothe R, Uphoff S, Leake MC, Sherratt DJ (2012) In vivo architecture and action of bacterial structural maintenance of chromosome proteins. Science 338(6106):528–531.

40. Zawadzka K, et al. (2018) MukB ATPases are regulated independently by the N- and C-terminal domains of MukF kleisin. Elife 7:1–26.

41. Banigan EJ, Mirny LA (2018) Limits of chromosome compaction by loop-extruding motors. bioRxiv. doi:10.1101/476424.

42. Mosteller RD, Yanofsky C (1970) Transcription of the tryptophan operon in Escherichia coli: Rifampicin as an inhibitor of initiation. J Mol Biol 48(3):525–531.

43. Griffith KL, Grossman AD (2008) Inducible protein degradation in Bacillus subtilis using heterologous peptide tags and adaptor proteins to target substrates to the protease ClpXP. Mol Microbiol 70(4):1012–1025.

44. Nicolas P, et al. (2012) Condition-dependent transcriptome reveals high-level regulatory architecture in Bacillus subtilis. Science 335:1103–1106.

45. Klumpp S, Hwa T (2008) Growth-rate-dependent partitioning of RNA polymerases in bacteria. Proc Natl Acad Sci 105(51):20245–20250.

46. Vogel U, Jensen KF (1994) The RNA chain elongation rate in Escherichia coli depends on the growth rate. J Bacteriol 176(10):2807–2813.

47. Lengronne A, et al. (2004) Cohesin relocation from sites of chromosomal loading to places of convergent transcription. Nature 430(6999):573–578.

48. Mizuguchi T, et al. (2014) Cohesin-dependent globules and heterochromatin shape 3D genome architecture in S. pombe. Nature 516(7531):432–435.

49. Davidson IF, et al. (2016) Rapid movement and transcriptional re-localization of human cohesin on DNA. EMBO J 35(24):2671–2685.

50. Wang MD, et al. (1998) Force and velocity measured for single molecules of RNA polymerase. Science 282(5390):902–907.

51. Golding I, Paulsson J, Zawilski SM, Cox EC (2005) Real-time kinetics of gene activity in individual bacteria. Cell 123(6):1025–1036.

52. Nora EP, et al. (2017) Targeted degradation of CTCF decouples local insulation of chromosome domains from genomic compartmentalization. Cell 169(5):930–944.

53. Guo Y, et al. (2015) CRISPR Inversion of CTCF Sites Alters Genome Topology and Enhancer/Promoter Function. Cell 162(4):900–910.

54. Diebold-Durand M-L, et al. (2017) Structure of Full-Length SMC and Rearrangements Required for Chromosome Organization. Mol Cell 67(2):334–347.e5.

55. Burmann F, et al. (2019) A folded conformation of MukBEF and cohesin. Nat Struct Mol Biol 26(3):227–236.

56. Nichols MH, Corces VG (2018) A tethered-inchworm model of SMC DNA translocation. Nat Struct Mol Biol 25(10):906–910.

57. Eeftens JM, et al. (2016) Condensin Smc2-Smc4 Dimers Are Flexible and Dynamic. Cell Rep 14(8):1813–1818.

58. Murayama Y, Uhlmann F (2014) Biochemical reconstitution of topological DNA binding by the cohesin ring. Nature 505(7483):367–371.

59. Murayama Y, Uhlmann F (2015) DNA Entry into and Exit out of the Cohesin Ring by an Interlocking Gate Mechanism. Cell 163(7):1628–1640.

60. Hu B, et al. (2011) ATP hydrolysis is required for relocating cohesin from sites occupied by its Scc2/4 loading complex. Curr Biol 21(1):12–24.

61. Shintomi K, et al. (2017) Mitotic chromosome assembly despite nucleosome depletion in Xenopus egg extracts. Science 356:1284–1287.

62. Lioy VS, et al. (2018) Multiscale Structuring of the E. coli Chromosome by Nucleoid-Associated and Condensin Proteins. Cell 172(4):771–783.e18.

## References

[1] Bicout, D. J. (1997). Green’s functions and first passage time distributions for dynamic instability of microtubules. Physical Review E, 56(6), 6656.

[2] Codling, E. A., Plank, M. J., & Benhamou, S. (2008). Random walk models in biology. Journal of the Royal Society Interface, 5(25), 813–834.

[3] Golding, I., Paulsson, J., Zawilski, S. M., & Cox, E. C. (2005). Real-time kinetics of gene activity in individual bacteria. Cell, 123(6), 1025–1036.

[4] Graham, T. G., Wang, X., Song, D., Etson, C. M., van Oijen, A. M., Rudner, D. Z., & Loparo, J. J. (2014). ParB spreading requires DNA bridging. Genes & development. 28: 1228–1238.

[5] Karp, P. D., Latendresse, M., Paley, S. M., Krummenacker, M., Ong, Q. D., Billington, R., … & Spaulding, A. (2015). Pathway Tools version 19.0 update: software for pathway/genome informatics and systems biology. Briefings in bioinformatics, 17(5), 877–890.

[6] Klumpp, S., & Hwa, T. (2008). Growth-rate-dependent partitioning of RNA polymerases in bacteria. Proceedings of the National Academy of Sciences, 105(51), 20245–20250.

[7] Le, T. B., Imakaev, M. V., Mirny, L. A., & Laub, M. T. (2013). High-resolution mapping of the spatial organization of a bacterial chromosome. Science, 342(6159), 731–734.

[8] Lindow, J. C., Kuwano, M., Moriya, S., & Grossman, A. D. (2002). Subcellular localization of the Bacillus subtilis structural maintenance of chromosomes (SMC) protein. Molecular microbiology, 46(4), 997–1009.

[9] Harwood, C. R., & Cutting, S. M. (1990). Molecular biological methods for Bacillus. Wiley.

[10] Imakaev, M., Fudenberg, G., McCord, R. P., Naumova, N., Goloborodko, A., Lajoie, B. R., … & Mirny, L. A. (2012). Iterative correction of Hi-C data reveals hallmarks of chromosome organization. Nature methods, 9(10), 999.

[11] Romero, P. R., & Karp, P. D. (2004). Using functional and organizational information to improve genome-wide computational prediction of transcription units on pathway-genome databases. Bioinformatics, 20(5), 709–717.

[12] Rudner, D. Z., Fawcett, P., & Losick, R. (1999). A family of membrane-embedded metalloproteases involved in regulated proteolysis of membrane-associated transcription factors. Proceedings of the National Academy of Sciences, 96(26), 14765–14770.

[13] Schroeder, J. W., & Simmons, L. A. (2013). Complete genome sequence of Bacillus subtilis strain PY79. Genome Announc., 1(6), e01085–13.

[14] Travers, M., Paley, S. M., Shrager, J., Holland, T. A., & Karp, P. D. (2013). Groups: knowledge spreadsheets for symbolic biocomputing. Database, 2013.

[15] Youngman, P. J., Perkins, J. B., & Losick, R. (1983). Genetic transposition and insertional mutagenesis in Bacillus subtilis with Streptococcus faecalis transposon Tn917. Proceedings of the National Academy of Sciences, 80(8), 2305–2309.

[16] Wang, X., Le, T. B., Lajoie, B. R., Dekker, J., Laub, M. T., & Rudner, D. Z. (2015). Condensin promotes the juxtaposition of DNA flanking its loading site in Bacillus subtilis. Genes & development, 29(15), 1661–1675.

[17] Wang, X., Brandão, H. B., Le, T. B., Laub, M. T., & Rudner, D. Z. (2017). Bacillus subtilis SMC complexes juxtapose chromosome arms as they travel from origin to terminus. Science, 355(6324), 524–527.

